# Neuronal loss reshapes survivor dynamics and limits mechanism inference in excitatory–inhibitory neural fields

**DOI:** 10.64898/2026.09.11.750823

**Authors:** Ronald Garcia Reyes, Pedro Antonio Valdés-Sosa

## Abstract

Does neuronal loss simply reduce measured activity, or also change how the surviving network behaves? We separate these effects in a next-generation excitatory–inhibitory neural field by writing the viable population measure as *q*_*a*_ = *λ*_*a*_*f*_*a*_, where *λ*_*a*_ is viable population mass and *f*_*a*_ is the normalized survivor distribution. Under state-independent thinning with fixed Cauchy heterogeneity, normalization commutes with the Ott–Antonsen/Montbrió–Pazó–Roxin reduction on the specified analytic invariant manifold. The mortality term disappears from conditional transport, but viable mass remains in recurrent coupling: loss can reshape survivor dynamics, not merely scale their contribution to tissue activity. Conversely, for otherwise identical constant homogeneous parameters, viability, pathway integrity and compensation give exactly conjugate conditional deterministic dynamics whenever 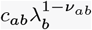 is preserved. Identical conditional activity therefore need not imply an identical biological mechanism. Equilibrium and oscillatory bifurcations, finite-population escape, and delayed propagation reveal consequences of these two principles. In particular, matched field simulations show that localized loss can increase whole-sheet firing through recurrent reorganization, while coherent-wave continuation quantifies viability-dependent propagation and phase relaxation. The framework distinguishes neuronal abundance from survivor state and places an exact limit on mechanism inference. Attributing activity changes to neuronal loss therefore requires information beyond conditional neural dynamics, such as tissue-level measurements or independent structural constraints, interpreted through an appropriate observation model.

## 1. Introduction

What does neuronal loss do to population dynamics? Fewer neurons contribute to activity, but fewer neurons also generate the recurrent input received by those that remain. A reduction in neuronal number therefore need not produce a proportional reduction in network activity. Survivors may change their firing, collective oscillations may appear or disappear, and a local loss may reorganize activity elsewhere. Interpreting an activity change requires separating this dynamical response from the direct loss of contributing cells.

We address that separation by writing the viable population measure as *q*_*a*_ = *λ*_*a*_*f*_*a*_. The viable fraction *λ*_*a*_ measures population mass relative to a reference population; *f*_*a*_ describes the normalized distribution of survivor states. Conditional firing rate *r*_*a*_ and phase moment *Z*_*a*_ refer to survivors, whereas *R*_*a*_ = *λ*_*a*_*r*_*a*_ and *M*_*a*_ = *λ*_*a*_*Z*_*a*_ describe tissue-level activity relative to the reference population. A high survivor rate can coexist with a weak tissue contribution. Crucially, viable mass also weights recurrent sources, so it is not a scale factor applied after solving an otherwise unchanged network.

The first theoretical question is whether this separation can retain an exact microscopic-to-macroscopic description. Classical excitatory–inhibitory (E/I) rate models prescribe interactions between populations^1^. Next-generation models instead derive their collective variables from quadratic integrate-and-fire (QIF) or theta-neuron dynamics through the Ott–Antonsen/Montbrió– Pazó–Roxin (OA/MPR) construction^2–5^. Their field extensions retain this connection in spatially coupled populations^6–8^. They offer a precise setting in which to distinguish a change in neuronal abundance from a change in the state law of the surviving neurons.

Under state-independent thinning, normalization removes the mortality term from conditional transport. We establish when this operation commutes with the OA/MPR reduction on an analytic invariant manifold, including the role of a fixed quenched trait coordinate when common excitability parameters vary. This is an invariance result; attraction from more general initial distributions is a distinct problem with its own hypotheses^9–11^. Relative to the variable-density neural field of Garcia Reyes and Martinez-Montes^12^, the contribution is the conditional law, its explicit closure conditions, independent E/I masses, and the resulting observation identities. A state-dependent hazard supplies a complementary counterexample: selective survival reshapes the conditional distribution and can destroy the reduced family’s invariance.

The second question is whether biologically different mechanisms remain distinguishable in the reduced dynamics. They need not. With all other parameters fixed, constant homogeneous viability, pathway integrity and compensation enter through products 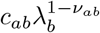. Preserving these products gives an exact linear conjugacy of the conditional deterministic systems, not just similar stationary rates. Corresponding trajectories, spectra, periodic dynamics and deterministic responses coincide. No increase in the precision of that same conditional observation can identify which mechanism produced it. This structural ambiguity is stronger than weak sensitivity or practical uncertainty, distinctions central to parameter inference in neural population models^13–15^.

Viability, intrinsic excitability, compensation, pathway integrity and propagation speed consequently remain separate generative coordinates even when some of their observations coincide. Models of activity-dependent degeneration and altered excitability^16, 17^, and of distributed-delay structural coupling^18^, motivate this separation without calibrating it. The present model is nondimensional and theoretical, not a model fitted to a disease stage or a clinical dataset.

We first derive conditional survivor dynamics and their tissue-level observables, then establish the exact mechanism-equivalence class and its consequences for observation design. The remaining analyses ask what changes when viable source strength is allowed to vary: E/I bifurcations describe reorganization of invariant states; finite-population theory separates sampling from noise-driven escape; and matched delayed fields show how local loss can increase global activity. Finally, coherent wave-trains and their Bloch spectra isolate propagation and phase relaxation as properties of a defined invariant solution. Computation tests these consequences of the two theoretical results; implementation and convergence controls are collected in the Supplementary Methods and Results. Detailed continuation and normal-form controls, finite-population validation, first-passage inference, spatial discretization tests, invariant-torus calculations and coherent-wave spectral convergence are cross-referenced from the corresponding analyses below.

## 2. A conditional-survivor E/I neural field

We represent neuronal number by viable mass and the state of the remaining neurons by a normalized conditional distribution. Recurrent input depends on both (Fig. 1).

**Fig. 1.**
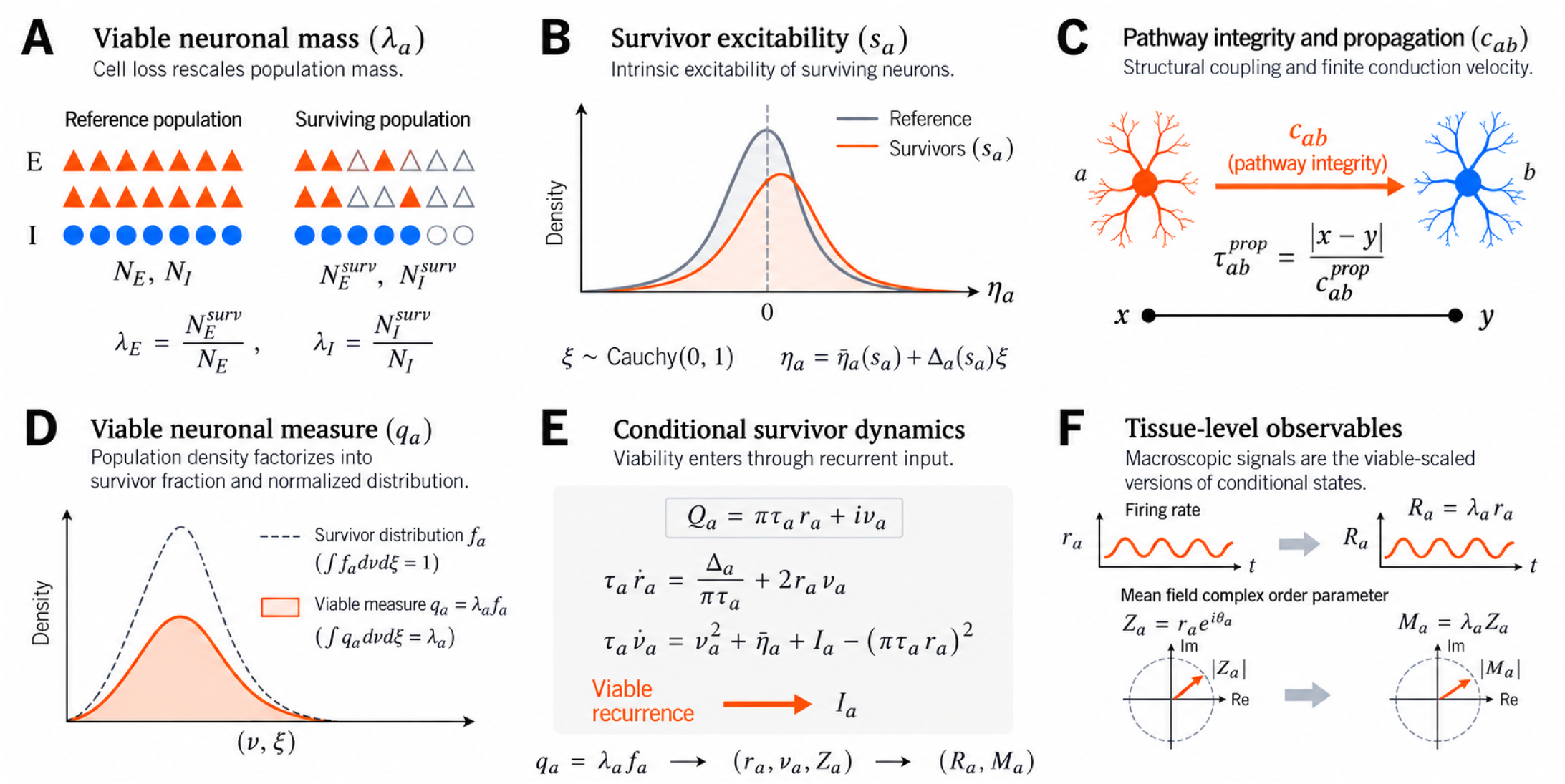
Loss, excitability and pathway changes act at distinct stages of the conditional-survivor model. **(A)** Class-selective thinning changes viable E/I counts. **(B)** Excitability modulation changes the excitability law of surviving neurons. **(C)** Integrity attenuates a pathway, whereas propagation speed changes transmission time. **(D)** The central factorization *q*_*a*_ = *λ*_*a*_*f*_*a*_ distinguishes the viable measure from its normalized survivor distribution. **(E)** Viable presynaptic mass feeds back into conditional survivor dynamics through recurrent input. **(F)** Conditional states (*r*_*a*_, *Z*_*a*_) and tissue-level observables (*R*_*a*_, *M*_*a*_) = (*λ*_*a*_*r*_*a*_, *λ*_*a*_*Z*_*a*_) describe different observation levels; the phase-plane radius is the magnitude of the conditional phase moment, not the conditional firing rate. In the artwork’s polar label, the radial factor must therefore be read as |*Z*_*a*_|, not *r*_*a*_. The text uses *c*_*ab*_ for transmission from class *b* to class *a*. Neuron and pathway drawings are schematic, not anatomical measurements.

### 2.1 Population mass and fixed heterogeneity

Let *a* ∈ {*E, I}* denote neuronal class and ***x*** spatial location. Reference density 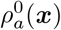 specifies the reference population against which mass is measured. Define

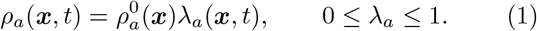

The conditional variables are defined where *λ*_*a*_ *>* 0. The E/I reference ratio is 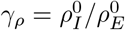, with (*γ*_*E*_, *γ*_*I*_) = (1, *γ*_*ρ*_) in the homogeneous reference model. Different viable fractions constitute *class-selective loss*. A hazard that depends on state within a class constitutes *state-dependent selection*. These terms will not be used interchangeably.

Each neuron receives a quenched label *ξ*_*aj*_ ~ *g*, with *g*(*ξ*) = [*π*(1+*ξ*^2^)]^−1^, held fixed throughout its evolution. A common survivor excitability control *s*_*a*_(***x***, *t*) may change intrinsic excitability through

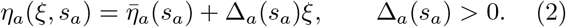

This control modulates the center or width of the intrinsic distribution. The QIF dynamics are

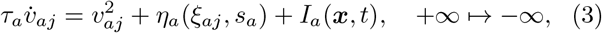

with fixed *τ*_*a*_ *>* 0. Under *θ* = 2 arctan *v* the same dynamics read

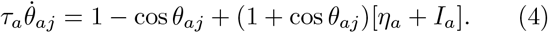

This compactification links spike/reset dynamics to phase moments^3, 19^.

### 2.2. Viable mass and conditional transport

Let *q*_*a*_(*v, ξ*, ***x***, *t*) be the unnormalized viable measure relative to the reference class population. At each location it obeys

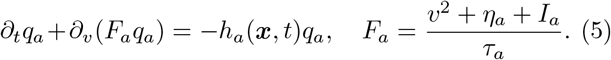

The neuronal-loss hazard *h*_*a*_ is independent of *v* and *ξ* within a class. Reset flux is conservative: spikes do not remove neurons.

#### Proposition 1

(Survivor factorization). *For conservative reset transport, positive initial mass and locally integrable state-independent hazard, the viable measure factors as*

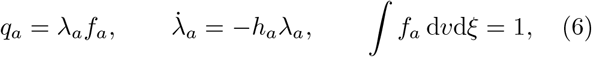

*and the normalized survivor distribution satisfies* ∂_*t*_*f*_*a*_ + ∂_*v*_(*F*_*a*_*f*_*a*_) = 0 *while λ*_*a*_ *>* 0.

The factorization identifies the normalization that separates neuronal number from survivor state. The mortality term cancels because it acts equally within a class, while the transport velocity still depends on viable recurrent input. The remaining question is whether this normalization is compatible with an exact conditional population reduction.

The QIF voltage coordinate is compactified by identifying its two infinities at the spike/reset point. Equivalently, *θ* = 2 arctan *v* lies on the circle. This avoids treating the outgoing firing flux as a loss of neurons. For a population at a fixed spatial location, let *q*(*v, ξ, t*) be the viable measure relative to the initial reference population. Its total mass is

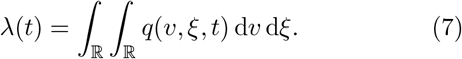

All integrals involving a density can equivalently be stated weakly for a measure. Conservative transport means that the voltage boundary fluxes agree at the identified infinities. The integral of ∂_*v*_(*Fq*) therefore vanishes. Integrating the killed transport equation gives

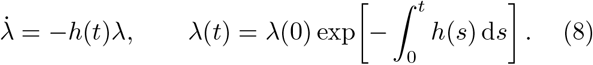

If *h* ≥ 0 is locally integrable and *λ*(0) *>* 0, the fraction remains positive at every finite time. Substituting *q* = *λf* and using Eq. (8) yields

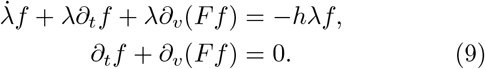

Conversely, a normalized solution *f* of this conservative equation, together with Eq. (8), reconstructs a solution *q* of the killed equation. This is a reversible correspondence as long as *λ >* 0. It does not remove the dependence of *F* on the viable recurrent source.

When *h* = *h*(*v, ξ, t*), the same calculation instead gives

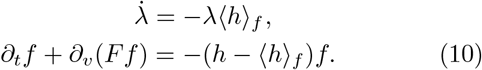

In particular, integrating over voltage shows that the quenched marginal *g*_*t*_(*ξ*) = ∫ *f* (*v, ξ, t*)d*v* obeys

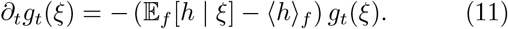

State-dependent selection can thus change both the conditional voltage law and the heterogeneity law. Different state-independent hazards in E and I classes do neither within a class; they change the two class masses separately.

### 2.3. Lorentzian ansatz and the fixed quenched coordinate

Conservative conditional transport alone does not supply a finite-dimensional closure. The Lorentzian/OA construction supplies that closure on a specified analytic family, with the quenched trait held fixed as excitability is controlled.

Write *f* (*v, ξ, t*) = *φ*(*v* | *ξ, t*)*g*(*ξ*), with *g*(*ξ*) = [*π*(1 + *ξ*)]^−1^. Initially require ∫ *q*(*v, ξ*, 0)d*v* = *λ*(0)*g*(*ξ*); the hazard must be independent of *ξ* as well as voltage to preserve this survivor marginal. The time constant *τ* is fixed within each neuronal class. On the Lorentzian manifold,

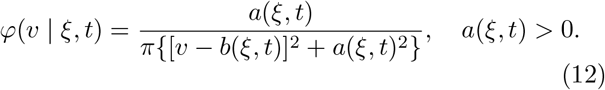

Inserting this ansatz into conservative QIF transport and equating powers of *v* − *b* gives

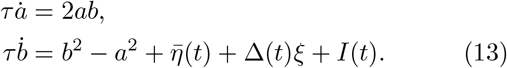

Here *I* is independent of the individual voltage and quenched label within the class, but may be generated self-consistently by all other populations and spatial locations. For *w* = *a* + i*b* these equations become

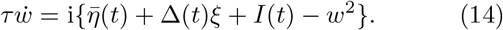

The individual firing flux is *a/*(*πτ*); the voltage location is a principal-value quantity. Assume that *w*(*ξ, t*) extends analytically into the lower half-plane, has no singularity there that obstructs contour closure, and has growth such that the integral over the closing semicircle vanishes. These are assumptions on the initial data and solution throughout the time interval under consideration. They are not conclusions for arbitrary initial densities. With

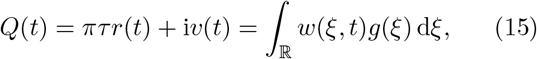

the clockwise lower contour evaluates the integral at *ξ* = −i:

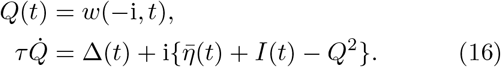

Taking real and imaginary parts gives the conditional MPR equations. The Lorentzian/OA construction is standard^2, 3^; the survivor theorem identifies the assumptions under which the population-mass factor can be separated without changing this conditional construction.

The pole is fixed in *ξ*, rather than in a time-varying physical excitability coordinate. If 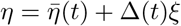 were used as the kinetic coordinate, its deterministic drift would be

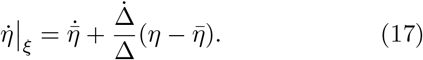

The corresponding density equation would contain an *η*-transport term. Using fixed labels already accounts for this coordinate motion. It generates no additional 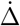 term in Eq. (16). Time-varying common excitability parameters are compatible with exact reduced constructions under their stated assumptions^20^; arbitrary heterogeneity laws generally need different or higher-dimensional reductions^21^.

The thinning transformation and conditional reduction commute in the following precise sense. Applying scalar thinning to the viable kinetic measure and then normalizing gives the same *f* as normalizing first and solving the conservative conditional problem with the same viable recurrent source. Evaluating the conditional analytic ansatz at the fixed pole therefore gives the same *Q* in both constructions. The mass *λ* is retained as an independent input to the recurrent functional.

#### Theorem 1

(Commutation of thinning and conditional MPR reduction). *Assume Proposition 1 and, for each class and location: (i) the initial survivor marginal of the fixed label is g*(*ξ*); *(ii) excitability has the affine form Eq*. (2), *with* Δ_*a*_ *>* 0 *and fixed τ*_*a*_; *(iii) the initial conditional voltage law belongs to the Lorentzian/OA family; (iv) its complex parameter admits lower-half-plane analytic continuation and contour closure throughout the time interval; and (v) recurrent input is common within the class and independent of the individual voltage and label. Then scalar thinning and conditional reduction commute. The survivor variables obey exactly on this invariant analytic manifold*

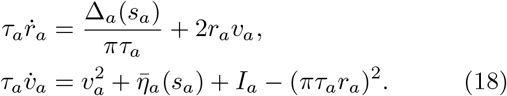

*The recurrent functional retains the viable presynaptic source. The conditional phase and tissue observables are*

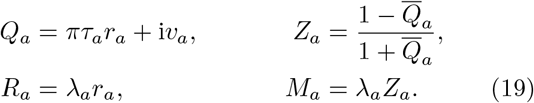

The reduced survivor equations contain no explicit mortality term, but they retain the viable population in recurrent input. The observation identities also retain viable-mass factors. Normalization therefore removes the sink term, not the dynamical or observational effects of neuronal loss.

*Proof*. State-independent thinning preserves the fixed-label survivor marginal and leaves conservative conditional transport. The Lorentzian parameter *w*_*a*_ = *a*_*a*_ + i*b*_*a*_ therefore satisfies its conservative Riccati equation with the same self-consistent viable input. The fixed Cauchy pole at *ξ* = −i gives *Q*_*a*_ = *w*_*a*_( −i) and 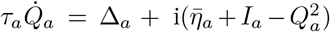. Its real and imaginary parts give Eq. (18). Normalization multiplies only the population measure, and the residue acts on the conditional analytic family; their order leaves the same conditional state and viable source. The phase transformation gives Eq. (19). The derivation above supplies the conditional transport and fixed-pole evaluation.

Exactness here concerns the stated invariant analytic manifold; attraction from arbitrary initial distributions is a separate question, as illustrated by attraction and beyond-manifold theories^4, 9–11, 22, 23^. The fixed *ξ* coordinate permits time-varying Δ_*a*_(*s*_*a*_) without 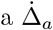 residue term. A physical-excitability coordinate instead requires the transport term associated with its motion.

### 2.4. Selective survival and the boundary of conditional closure

State-independent class loss preserves the conditional heterogeneity law. State-dependent loss instead selects among survivor states or traits and changes the conditional distribution itself. The following counterexample identifies why the four-variable conditional neuronal closure need not remain invariant under such selection.

A positive OA phase density has moments *Z*_*n*_ = *Z*^*n*^, *n* ≥ 1, and |*Z*| *<* 1. Consider a legitimate hazard *h*(*θ*) = *h*_0_ + *ϵ* cos *θ*, with *h*_0_ ≥ |*ϵ* |. At an instant when the density is on the OA manifold, the selection contribution to its moments is

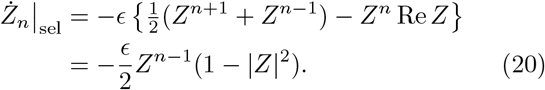

Tangency to *Z*_*n*_ = *Z*^*n*^ would require *Ż*_*n*_ = *nZ*^*n*−1^*Ż*_1_, introducing an extra factor *n*. Already the second moment fails this condition generically. Thus a state-dependent hazard can invalidate the OA closure even when its first-moment equation appears simple. The failure is a loss of tangency to the OA family, not a failure of population-mass book-keeping. Special structured hazards may preserve other reduced families, but require their own invariance analysis.

### 2.5. Viable recurrence, compensation and propagation

Each surviving presynaptic neuron contributes to the recurrent source. Compensation changes the source amplitude, pathway integrity scales its transmission, and propagation speed sets its travel time.

Let *J*_*ab*_ be a signed coupling from presynaptic class *b* to postsynaptic class *a, c*_*ab*_ its pathway integrity, and 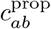 its propagation speed. The compensated viable source is

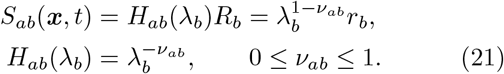

The conditional problem is defined for positive viable fractions; the *λ* = 0 state has no conditional survivor distribution. The source is filtered in space and time:

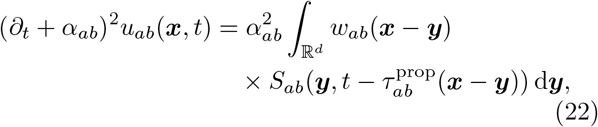

On a periodic domain, *S*_*ab*_ is periodically extended over R^*d*^. Thus different image paths retain different propagation delays, even when they connect the same pair of locations in the periodic cell. A single minimum-distance delay multiplying a periodized spatial weight is not the operator in Eq. (22).

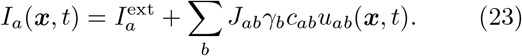

In the fixed-parameter numerical fields, *c*_*ab*_ is homogeneous and viability can vary in space. More general spatial reference densities would be included inside the source integral. Viability and compensation are evaluated at emission time in Eq. (22), not retrospectively at arrival time.

For the homogeneous undelayed system,

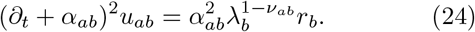

The four filters add eight states to the four conditional neuronal states. The normalized alpha response has unit zero-frequency (DC) gain. Compensation changes how the presynaptic mass enters recurrence; it does not restore missing cells or make a tissue observation equal to its conditional counterpart.

### 2.6. Dimensional and biological scope

All model parameters are nondimensional. A conversion to physical time requires a separately chosen time scale; therefore an angular frequency *ω* is reported in model-time^−1^, while a cycle frequency is *ω/*(2*π*). Conditional rates are per survivor, tissue-level observables are per reference neuron of the relevant class, and a physical source density would include 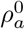. For example, a summed firing contribution per reference excitatory density is *R*_*E*_ +*γ*_*ρ*_*R*_*I*_, not *R*_*E*_ + *R*_*I*_ when reference densities differ.

These parameters describe population size, intrinsic excitability and transmission, not clinical disease stages. Except for the explicitly imposed slow-control protocol, numerical experiments hold viability fixed during each realization. Random removal is quenched before microscopic simulation, and finite-count trials have fixed viable counts. The continuum theorem permits time-dependent state-independent hazards, but the finite-size studies do not simulate ongoing stochastic neuronal deaths.

### 2.7. Reference parameterization

The reference parameters in the nondimensional convention above are

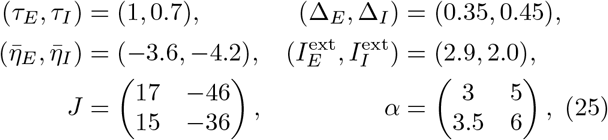

with *γ*_*I*_ = 0.25, *c*_*ab*_ = 1, *ν*_*ab*_ = 0, and zero additional survivor excitability shift unless otherwise specified. A shared configuration specifies these parameters. The two viable fractions are independently controlled; within each class thinning is independent of phase and quenched trait. The numerical design and organization of the computational details are described in Supplementary Sec. 1.

## 3. Conditional and tissue-level observables

The theorem separates probability dynamics from population mass. An observation must specify whether it is normalized by the viable or reference population. The conditional firing rate *r*_*a*_ and conditional phase moment *Z*_*a*_ describe survivors, whereas the tissue-level firing rate *R*_*a*_ = *λ*_*a*_*r*_*a*_ and tissue-level coherent moment *M*_*a*_ = *λ*_*a*_*Z*_*a*_ refer to the reference population.

The phase transformation *θ* = 2 arctan *v* gives

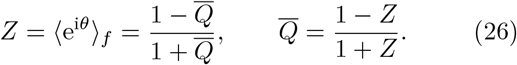

This maps the positive-rate half-plane to the unit disk. Consequently,

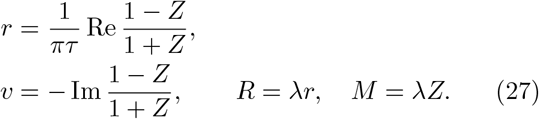

The conditional phase moment *Z* measures phase concentration. A large magnitude can reflect neurons concentrated near a resting phase rather than rhythmic synchronization, coincident spiking or strong tissue oscillation. The Cauchy voltage location *v* is not an ordinary integrable voltage mean. In a finite simulation, voltage inferred through Eq. (27) is derived from *Z* and does not constitute an independent validation of the closure.

Along matched loss, *λ*_*E*_ = *λ*_*I*_ = *ℓ*, a stable equilibrium or periodic conditional state can remain strongly structured while multiplication by *ℓ* reduces its tissue-level firing rate (Fig. 2). Conditional rates measure spikes per viable neuron; tissue-level firing rates measure spikes per reference neuron. Similarly, *Z*_*a*_ is the conditional phase moment, while *M*_*a*_ combines its phase organization with viable mass. The latter remains within a disk of radius *λ*_*a*_, rather than the unit disk.

**Fig. 2.**
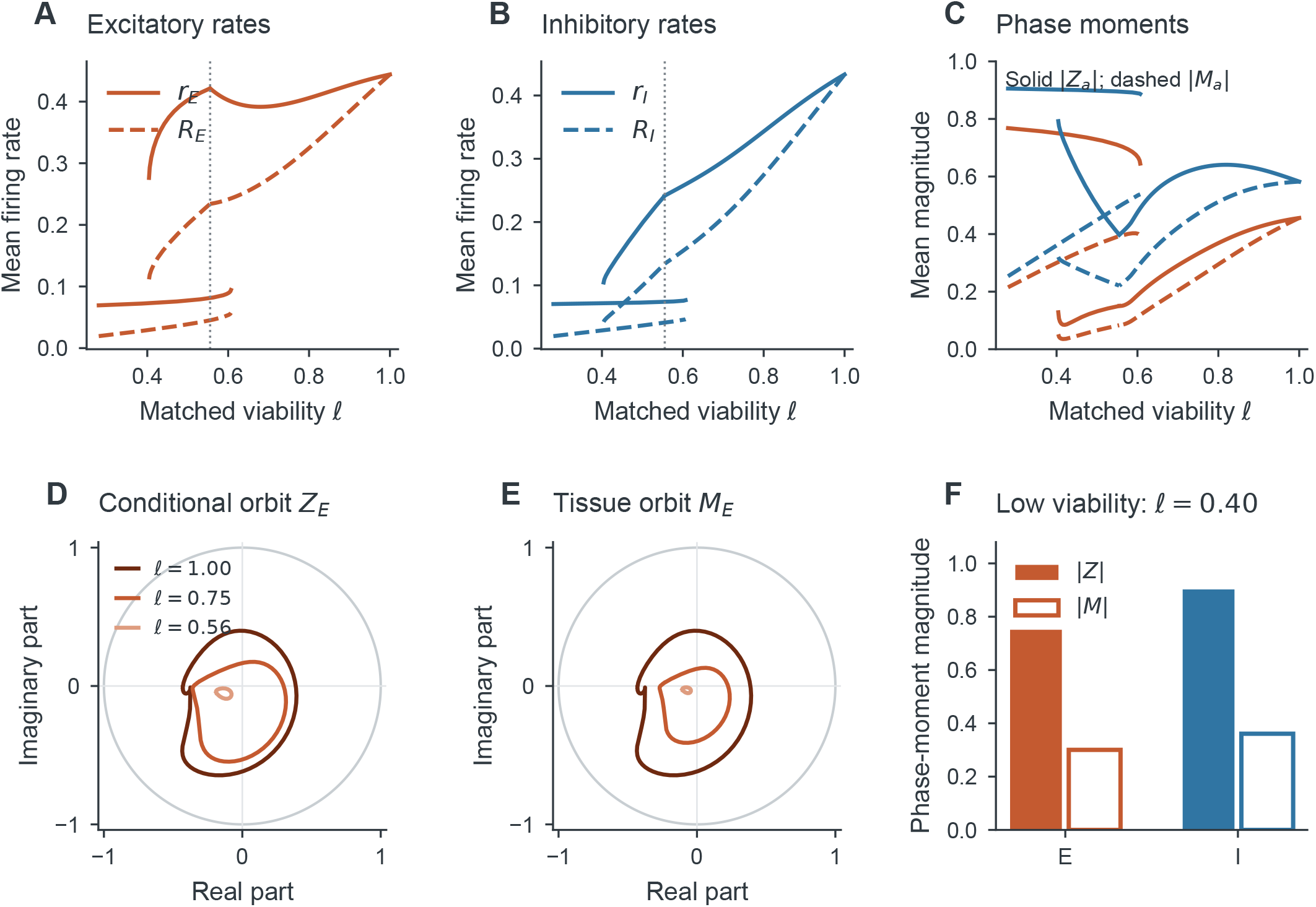
Survivor organization and tissue-level activity use different normalizations. **(A**,**B)** Conditional E and I firing rates (solid) and tissue-level firing rates (dashed) along matched viability. Periodic branches show cycle means; equilibrium branches show stable stationary values. **(C)** Mean magnitudes of the conditional phase moments and tissue-level coherent moments. **(D**,**E)** The same E-population periodic orbits as *Z*_*E*_ and *M*_*E*_ = *ℓZ*_*E*_ at three continued viable fractions. **(F)** At *ℓ* = 0.4, strong conditional phase concentration coexists with a much smaller tissue-level coherent moment. Concentration near a low-rate resting state is distinct from rhythmic synchronization. All panels use continued states or the corresponding direct trajectory.

The tissue-level coherent moment *M*_*a*_ is a model observable, not a forward solution for electroencephalography (EEG), magnetoencephalography (MEG) or local field potentials (LFPs). Source geometry and orientation, conductivity, synaptic or transmembrane currents, volume conduction and sensor mixing are needed before a measured field could be predicted. The exact normalization identities specify which population-level quantities such a forward model would combine.

Separating these normalizations specifies what an observation measures. It also exposes an inverse question: can the measured dynamics distinguish loss of viable neurons from changes in the pathways and compensation that remain?

## 4. Mechanistic distinction and observational equivalence

If viability, pathway integrity and compensation are biologically distinct, can population recordings distinguish them? The separation of mass and state does not by itself ensure that they can. With all other neuronal, synaptic and input parameters held fixed, a change of synaptic coordinates identifies an exact equivalence among constant homogeneous mechanisms.

### Proposition 2

(Conditional deterministic equivalence class). *For fixed homogeneous parameters with c*_*ab*_ *>* 0, *two choices of viability, integrity and compensation that preserve every product* 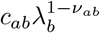 *have conjugate conditional deterministic dynamics under z*_*ab*_ = *c*_*ab*_*u*_*ab*_. *With corresponding initial synaptic states they have identical conditional neuronal trajectories and spectra. Their tissue observables and effective viable counts may differ*.

The answer is therefore no within this equivalence class: identical survivor dynamics need not have the same biological cause. The result concerns whole conditional trajectories, not merely a fitted firing rate. Intrinsic changes of 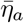 or Δ_*a*_ generally do not belong to this product equivalence class, although a stationary firing rate can still be matched.

### 4.1. Exact deterministic product equivalence

For constant homogeneous parameters define

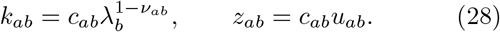

If *c*_*ab*_ *>* 0, the map 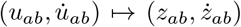 is invertible. The synaptic and input equations become

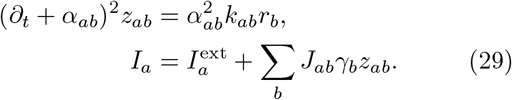

Thus two parameter choices with equal *k*_*ab*_ are linearly conjugate in their conditional deterministic states. Matching transformed initial conditions gives identical *r, v* trajectories; Jacobians at corresponding equilibria are similar, and monodromy matrices on corresponding periodic orbits are similar. This is stronger than equality of a stationary mean. Tissue observables and finite effective counts can still differ.

When integrity varies in time, the product rule introduces additional terms:

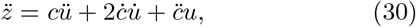

and a spatially varying source factor cannot generally be pulled outside a nonlocal delayed convolution. At *c* = 0 the transformation is not invertible. The conjugacy is thus a constant-parameter, homogeneous result; these other settings require analysis of the additional temporal or spatial terms.

### 4.2. Observation design and parameter redundancies

The inverse problem depends on what is measured. Stationary summaries may match even when relaxation and forced responses differ; tissue-level measurements add mass information only for the population being observed. We use a local design matrix to distinguish such weakly informative measurements from exact parameter redundancies.

Let ***g***(***θ***) collect the chosen model observations. In scaled parameter coordinates 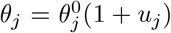 and fixed observation scales 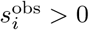, define

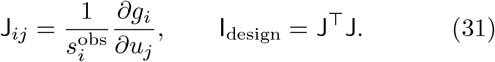

This transpose construction requires a real observation vector: each complex frequency response must contribute separate real and imaginary components to ***g***, making J real. If an empirical observation covariance is available, a statistical information construction uses 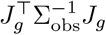, with *J*_*g*_ = *D*_***θ***_***g***, under the specified measurement likelihood. Here identity covariance after fixed scaling supplies only a design diagnostic. Singular values and right singular vectors identify parameter combinations that are poorly separated in this chosen observation space. This distinction between structural redundancy and practical inference is central to identifiability analysis, including neural population models^13–15^.

An exact redundancy is

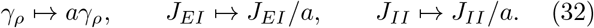

For *a >* 0, it preserves all conditional deterministic input products. In fractional coordinates its tangent is proportional to 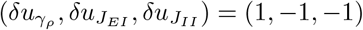, where each subscript on *u* names the corresponding parameter. Neither additional precision nor additional statistics of the same conditional dynamics can remove this gauge. Separating its members requires information that does not share the symmetry, such as an independent reference-density or coupling measurement.

Perturbations test dynamical information absent from stationary summaries. For a stable equilibrium, a small harmonic input ***b***e^i*ωt*^ has linear response

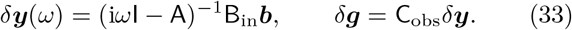

Here B_in_ denotes the input matrix, distinct from the state Hessian B in Eq. (40). Responses can reveal directions absent from summary measurements without guaranteeing practical estimability under empirical noise. Observation rank is always reported with the measurement set, parameter scales, finite-difference refinement and rank tolerance.

For an example of Proposition 2, consider the inhibitory pathways. At (*λ*_*E*_, *λ*_*I*_) = (0.6, 0.9) with *ν*_*aI*_ = 0.5, three changes have the same effective inhibitory presynaptic factor: reducing *λ*_*I*_ to 0.81, setting 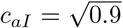, or with-drawing compensation to *ν*_*aI*_ = 0. They yield identical conditional rates and spectra when synaptic initial conditions are matched by the conjugacy. Tissue-level I firing nevertheless differs because only the first intervention changes I viable mass. Viable counts also change the mesoscopic noise covariance.

An inhibitory intrinsic-center change can match the same *r*_*E*_, but gives a different *r*_*I*_ and leading eigenvalue. Thus matching one rate is weaker than matching the dynamics. Numerical values for these comparisons are given in Supplementary Sec. 7. Perturbation responses can help separate such alternatives, provided that their observation and uncertainty model is stated.

In a nine-parameter local design, stationary summaries plus a relaxation eigenvalue give numerical rank five. Adding complex responses at four frequencies increases the rank to eight. The remaining null direction combines the inhibitory reference density and its two outgoing couplings, proportional to (1, −1, −1), as predicted by the exact multiplicative redundancy in Eq. (32). Right singular vectors, coordinate scales and the nonzero singular values are shown in the main observation-design figure; the full nine-value spectra, including zeros, are given in Supplementary Table S2. Scaling and finite-difference tests are specified in Supplementary Sec. 2. This local design result concerns the specified observations and scales; practical inference additionally depends on the recording model and uncertainty^14, 15^.

### 4.3. Frequency-resolved mechanism comparisons

Product-equivalent mechanisms have identical conditional transfer functions, whereas mechanisms matched at one stationary rate need not. For a unit excitatory external-current perturbation, we evaluate Eq. (33) across angular frequencies 0.01 ≤ *ω* ≤ 30. Fig. 3 displays the response amplitude and phase. The viable-fraction, pathway-integrity and compensation changes preserve their conditional transfer functions over the entire range. The intrinsic-excitability match has a different response, although it shares one stationary rate with the other mechanisms.

**Fig. 3.**
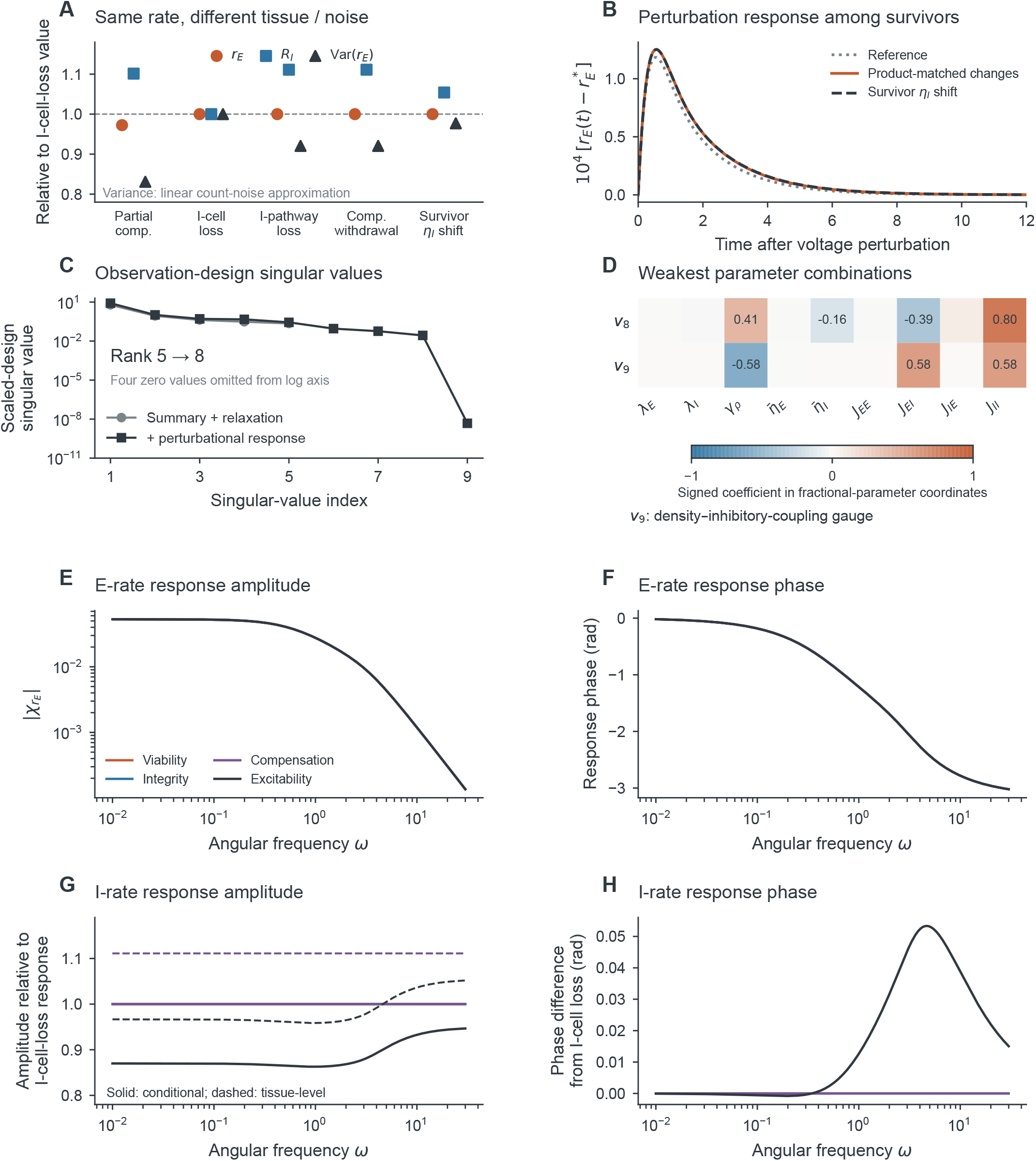
Distinct mechanisms have identical conditional responses, but different tissue weights can distinguish them. **(A)** Conditional E rate, tissue-level I rate and linear count-noise variance relative to the I-cell-loss case. **(B)** Conditional E response to the same voltage perturbation. **(C)** Singular values of the summary/relaxation and response-augmented designs. **(D)** Their two weakest augmented-design right singular vectors; *v*_9_ is the exact density–coupling gauge. **(E**,**F)** E response amplitude and phase to unit E current over 401 frequencies. **(G**,**H)** I amplitudes and phases relative to the I-cell-loss response. Solid lines show conditional amplitudes and dashed lines tissue-level amplitudes. Viability, integrity and compensation responses overlap conditionally; tissue-level I amplitudes distinguish different I viable fractions. The stationary-rate-matched excitability change differs dynamically. The design uses fixed scales and identity working covariance, not an empirically calibrated noise model.

The identity 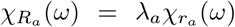 gives the tissue-level response at fixed viable fraction. Here all product-equivalent cases share *λ*_*E*_ = 0.6, so tissue-level E activity cannot distinguish them. Their different inhibitory viable fractions do separate the tissue-level I amplitudes. Multiplication by a positive viable fraction leaves response phase unchanged. Which population and normalization are observed is consequently part of the mechanism-identification problem.

The equivalence identifies the recurrent products that determine conditional dynamics. We next vary E and I viability while holding the other mechanisms fixed, asking how those products reshape the network’s invariant states.

## 5. E/I viability and bifurcation structure

Independent E and I loss changes the recurrent coupling and can move the system between stationary and oscillatory states. We locate the equilibrium and Hopf boundaries, then calculate the local nonlinear dynamics near folds, generalized-Hopf points and a double-Hopf intersection.

### 5.1. Equilibria under independent E/I viability

Define 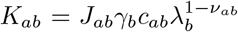 for fixed homogeneous parameters. At an equilibrium of the full synaptic system,

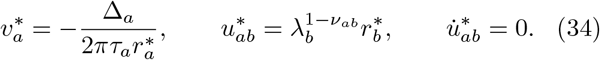

The two positive rates solve

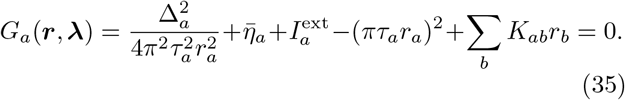

The numerical coordinates *x*_*a*_ = log *r*_*a*_ enforce positive rates. Their exact first derivatives are

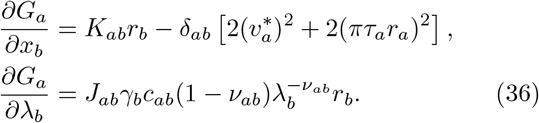

The indicator *δ*_*ab*_ here is the Kronecker delta, distinct from the scalar fold-distance coordinate used below. A singular derivative of *G* can identify an equilibrium fold, but the stability of that equilibrium must be computed from the full twelve-dimensional dynamics, including the synapses.

### 5.2 Stability of the coupled neuronal and synaptic state

Equilibrium rates locate a state, but its stability depends on relaxation through all recurrent pathways. We retain the synaptic variables explicitly and compute the full state derivative. Use the state order

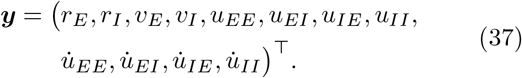

Let A = *D*_***y***_*F* and *L*_*ab*_ = *J*_*ab*_*γ*_*b*_*c*_*ab*_. The neuronal blocks have entries

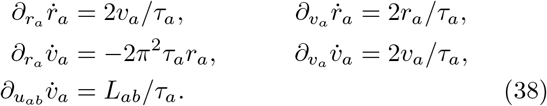

For each pathway, the synaptic block is

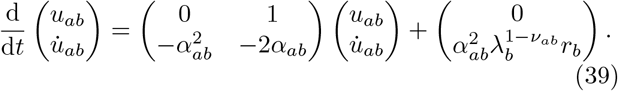

These entries determine the full Jacobian without differentiating trajectories. For fixed parameters the vector field is quadratic in the state. Define its bilinear second derivative by 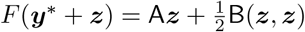. For arbitrary directions ***u, w*** its only nonzero components are

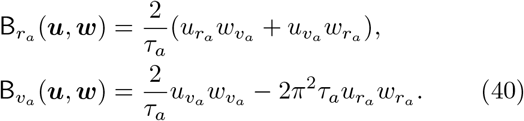

The third state derivative vanishes. This does not imply a vanishing cubic normal form: quadratic terms feed back through the center-manifold embedding.

### 5.3. Following fold and Hopf boundaries

To distinguish genuine stability boundaries from a finite parameter grid, we continue the equilibrium constraints and their critical eigenmodes. Pseudo-arclength permits these curves to turn in either viable fraction. For an *m*-equation curve ℱ (***z***) = 0 in *m* + 1 coordinates, the unit tangent ***t***_*n*_ lies in the nullspace of *D*ℱ(***z***_*n*_), with orientation chosen continuously. A predictor is ***z***_pred_ = ***z***_*n*_ + Δ*s****t***_*n*_. The corrector solves

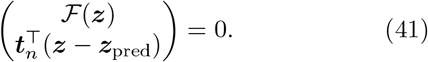

This continuation follows turning branches that a parameter grid alone can miss^24, 25^. The continuation and normal-form implementation is described in Supplementary Sec. 2; coordinate choices, termination rules and reference values are given in Supplementary Sec. 7.

For fold curves the augmented equations are

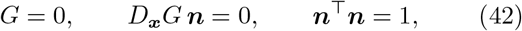

in (*x*_*E*_, *x*_*I*_, *λ*_*E*_, *λ*_*I*_, ***n***). Hopf curves solve equilibrium equations together with Re *µ*_*H*_ = 0 and Im *µ*_*H*_ = *ω*, tracking the same positive-frequency eigenpair. The transverse spectrum distinguishes a stability boundary from a Hopf bifurcation of an already unstable equilibrium.

### 5.4. Reference bifurcations along three loss paths

Continued folds delimit a region of low/high-rate coexistence (Fig. 4). Representative parameter points connect this geometry to a low-rate equilibrium, two stable equilibria separated by a saddle, a high-rate equilibrium, and a stable oscillation. Stability is assigned from the full state spectrum and periodic-orbit calculations.

**Fig. 4.**
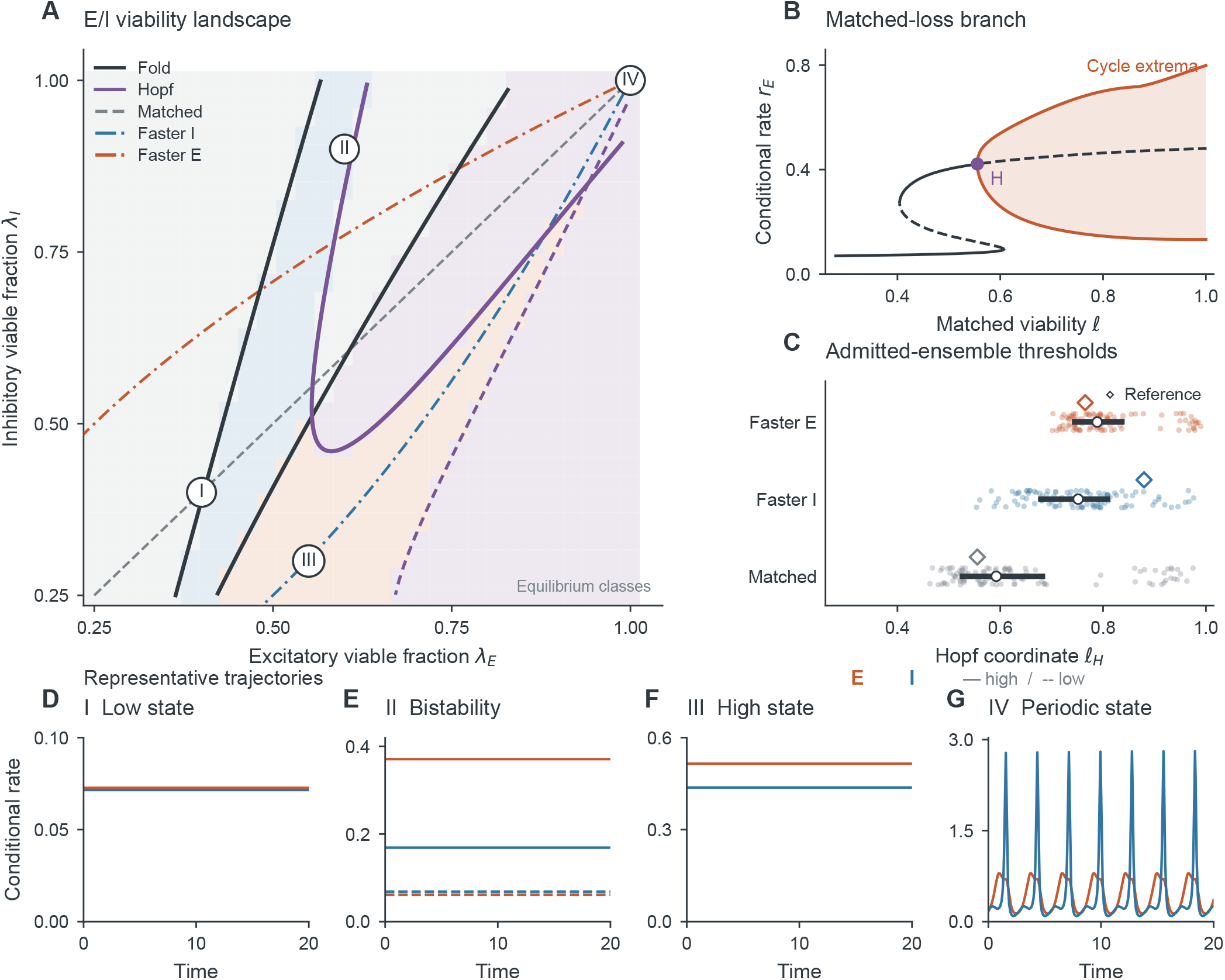
E/I viability reorganizes invariant states, and class-specific transition ordering depends on the recurrent parameters. **(A)** Continued fold and Hopf curves on the exploratory equilibrium grid, with matched, faster-I and faster-E loss paths. **(B)** The matched periodic branch, calculated by shooting rather than a finite-window amplitude threshold. **(C)** Regular pathwise Hopf crossings among 128 parameter sets supporting an oscillatory intact state; 87 have comparable crossings on all three paths. Marginal regular-path counts are 104, 99 and 99 for matched, faster-I and faster-E paths; topology changes and unresolved members remain separate categories. **(D–G)** Independently simulated low-rate, bistable, high-rate and periodic conditional states at the marked points in **A**. Higher-order structure is resolved in Fig. 5; the near-Hopf transient appears in Supplementary Fig. S1.

The high-state fold on *λ*_*I*_ = 0.9 occurs at *λ*_*E*_ ≃ 0.53852. With the nullvector convention in Eq. (45), its coefficients are *A*_*F*_ ≃ −3.13025 and *B*_*F*_ ≃ 1.82023. These coefficients fix the local coordinate, potential and noise projection used in the escape analysis.

The three loss paths are parameterized by a decreasing coordinate *ℓ*:

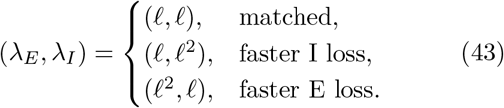

Their reference high-state Hopf crossings occur at approximately *ℓ* = 0.5553, 0.8797 and 0.7652, respectively. Thus faster I loss encounters this transition earlier in the reference model. All three crossings are transverse and supercritical; their coefficients and frequencies are reported in Supplementary Sec. 7.

Periodic shooting, defined in Eqs. (52) and (53), follows the matched stable orbit from the intact state down to the Hopf neighborhood. The stable branch approaches zero amplitude at Hopf. Just beyond the boundary, at *ℓ* = 0.55, an oscillatory perturbation instead decays slowly (Supplementary Fig. S1). Following the invariant periodic solution separates this long transient from sustained oscillation.

### 5.5. Parameter dependence of class-specific transition ordering

To test whether the reference ordering persists under parameter variation, coupling magnitudes, heterogeneity widths, intrinsic centers, synaptic rates and the reference density ratio were independently perturbed by up to 10%. We retain 128 parameter sets supporting an oscillatory intact state; the dynamical and continuation criteria are given in Supplementary Sec. 7.

All three paths had comparable regular Hopf crossings in 87 of these parameter sets. Among these, faster I loss crossed first in 21 cases (24.14%, bootstrap 95% interval 16.09–33.33%), and faster E loss in 66 (75.86%, 66.67–83.91%). Thirty-eight members showed verified path topology changes and three retained unresolved path calculations. They remain explicit categories rather than being counted as evidence for either ordering. The distributions in Fig. 4 summarize regular local Hopf crossings in a dynamically selected neighborhood, not global cycle-loss thresholds or a probability distribution over biological populations.

### 5.6. Local fold geometry and the soft mode

Near a fold, the stable state approaches a saddle and relaxation slows. The local reduction quantifies both effects and supplies the escape barrier used in Sec. 7. At a simple zero eigenvalue choose real right and left null vectors ***e, ℓ*** such that ∥***e***∥_2_ = 1 and ***ℓ***^⊤^***e*** = 1. In a transverse scalar parameter *δ*,

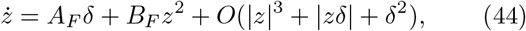

where

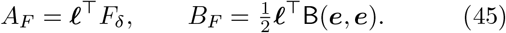

A regular fold requires a one-dimensional nullspace, a spectral gap to the remaining eigenvalues, and nonzero *A*_*F*_ and *B*_*F*_ for the stated transverse direction. Along a two-parameter fold curve, the vector ***ℓ***^⊤^*F*_***λ***_ identifies possible transverse directions; a particular coordinate can be tangent even when the fold is regular.

Choose the orientation *B*_*F*_ *>* 0 and take *A*_*F*_ *δ <* 0. Put 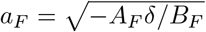. The center-stable and center-unstable coordinates are −*a*_*F*_ and +*a*_*F*_. With stable transverse modes, as at the reference fold, these correspond to a stable state and a saddle. With

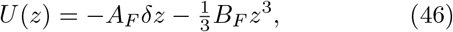

the barrier and local relaxation rate are

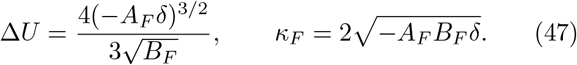

The potential is a one-dimensional normal-form construction near the fold, not a global potential for the E/I network. Projection outside that local neighborhood need not give quantitatively accurate escape actions.

### 5.7. Hopf criticality and the generalized-Hopf point

A Hopf boundary alone does not determine whether emerging cycles are stable or whether a small coexistence region is possible. The cubic coefficient determines criticality; where it vanishes, a fifth-order calculation determines the nearby cycle branches. At a simple Hopf pair let

A***q*** = i*ω****q***, A^⊤^***p*** = −i*ω****p***, ∥***q***∥_2_ = 1 and ***p***^†^***q*** = 1. Under the convention *F* = A***z*** + B(***z, z***)/2 + C(***z, z, z***)/6, the first Lyapunov coefficient is

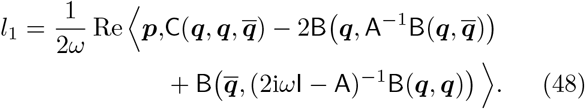

Here C = 0 for the reference model; the eigenvector normalization above fixes the coefficient’s amplitude convention. For an equilibrium branch ***y***^∗^(*s*), the crossing derivative is

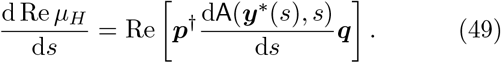

Its total derivative includes equilibrium motion, not just an explicit parameter derivative at a fixed state. A regular supercritical stability transition requires the crossing, *l*_1_ *<* 0 under this convention, and stable transverse modes.

Following the high-order normal-form approach to generalized Hopf bifurcations^26^, where *l*_1_ = 0, fifth-order coefficients are obtained by solving polynomial invariance equations, rather than numerically differentiating a simulated amplitude. Write the center embedding and reduced flow using ordinary, not factorial-weighted, coefficients:

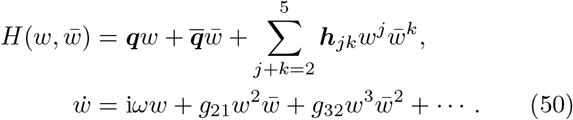

At each degree, known lower-order coefficients supply a forcing ***f***_*jk*_. Nonresonant terms solve

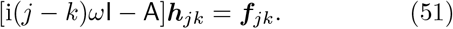

For *j* − *k* = 1, a bordered solve adds ***q****g*_*jk*_ on the left and imposes ***p***^†^***h***_*jk*_ = 0. Conjugate coefficients enforce reality. The reported normalization is *l*_1_ = Re *g*_21_*/ω* and *l*_2_ = Re *g*_32_*/ω*. These are coefficients in rescaled time *s* = *ωt*, rather than the original-time coefficients *g*_21_ and *g*_32_ in the displayed flow. The second equality fixes the fifth-order convention; numerical values from other conventions need not coincide. Independent cubic/quintic normal forms and a nonlinear coordinate shear test the implementation.

A generalized Hopf point additionally requires a regular two-parameter unfolding and *l*_2_ ≠ 0. A double-Hopf spectral point requires two distinct imaginary pairs and independent parameter crossings. Its nonlinear classification additionally requires cross-mode coupling coefficients and a check for low-order resonances. These quantities and the associated invariant objects are calculated below.

### 5.8. Periodic branches and their stability

Normal forms predict local periodic branches, whereas shooting locates them in the full system and Floquet multipliers determine their stability. For an autonomous flow *ϕ*_*T*_ (***y***_0_; *s*), the periodic-orbit equations are

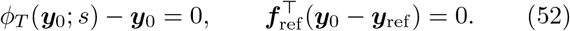

The second equation removes time-translation freedom. Its reference is updated locally along the branch to avoid tangencies to a fixed global phase plane. The fundamental matrix is integrated with the orbit:

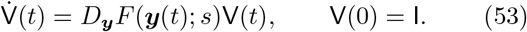

The monodromy V(*T*) gives Floquet multipliers. The derivative of the shooting residual with respect to log *T* is *TF* (***y***(*T*); *s*); parameter sensitivity obeys the inhomogeneous variational equation. In general, V(*T*) is not the exponential of the time-integrated Jacobian because Jacobians at different times do not commute.

An autonomous orbit has a neutral multiplier +1. Stability depends on all other multipliers. At a cycle fold a second real multiplier reaches +1; the phase-neutral direction must be separated when refining this event. Raw coalescing eigenvalues are sensitive to numerical roundoff, so the augmented shooting and neutral-lifted multiplier residuals are also reported. Passing through zero amplitude near a Hopf point can parameterize the same physical periodic family twice with different phase; this is not itself a fold of cycles. Periodic solutions and their stability are mathematical objects separate from finite-network sample trajectories^27, 28^.

### 5.9. A resolved coexistence region near generalized Hopf

The reference Hopf curve contains a generalized-Hopf point at

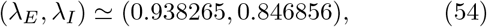

where *ω* 2. ≃ 68098, *l*_1_ vanishes numerically, and *l*_2_ = −0.00646343. Its negative sign organizes the stable outer cycle near the change in Hopf criticality. The convention is given in Eqs. (50) and (51); coefficient and continuation checks are reported in Supplementary Sec. 9.

At *λ*_*I*_ = 0.84975, the Hopf is weakly subcritical. Continued periodic solutions connect small unstable cycles to larger stable cycles through a fold at *λ*_*E*_ ≃ 0.9405982. The coexistence interval is narrow: the fold lies approximately 5.803 × 10^−7^ in *λ*_*E*_ from Hopf, in close agreement with the quintic prediction. Fig. 5C,D magnifies this separation and its stability change. The nearby intersection of two Hopf curves requires a further step: the two oscillatory modes must be analyzed together.

**Fig. 5.**
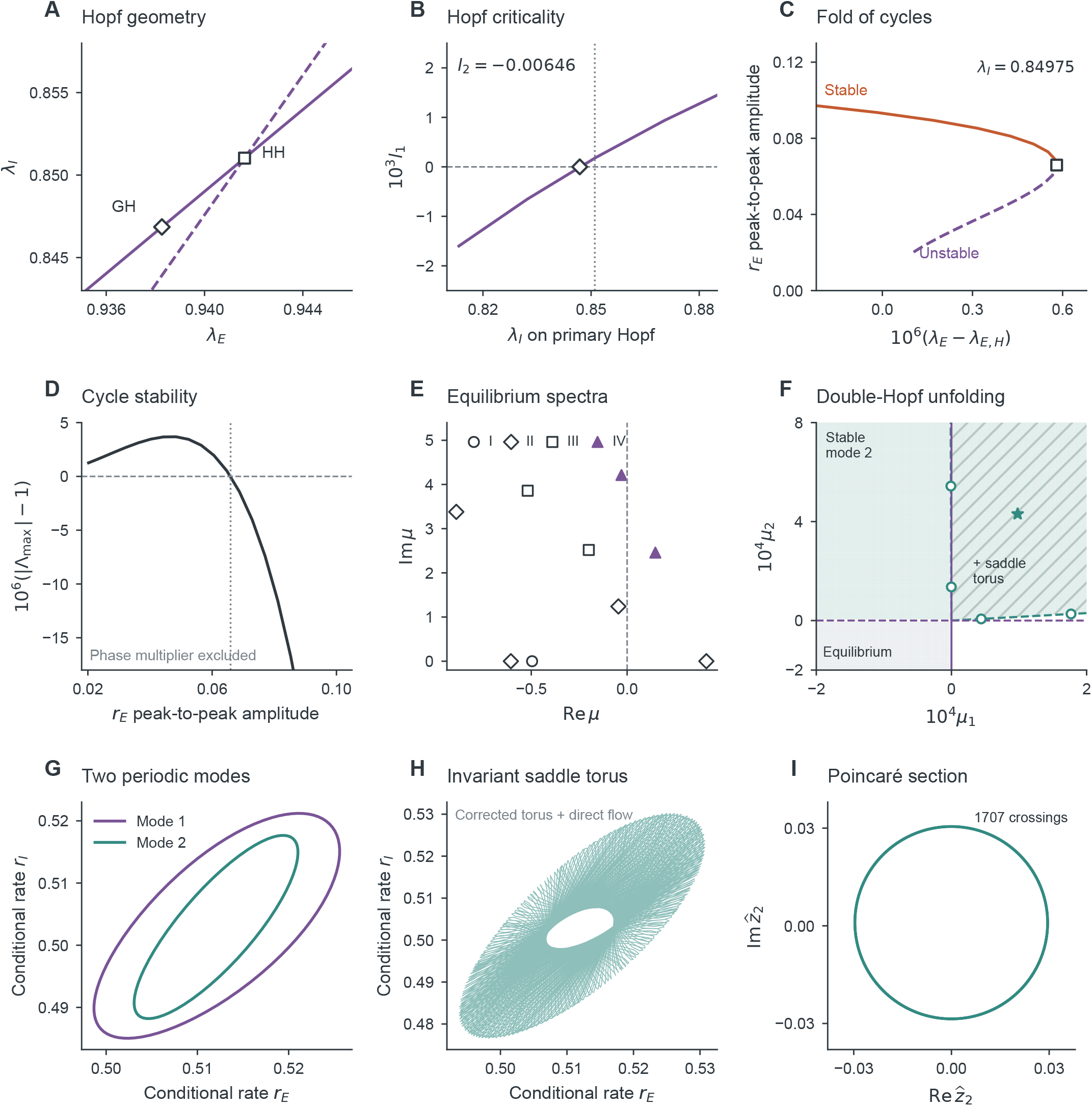
Viability changes Hopf criticality and organizes interacting oscillatory modes. Geometry and stability **(A–E)** connect to the local double-Hopf unfolding and its saddle torus **(F–I). (A)** Continued Hopf curves, generalized Hopf (GH) and double Hopf (HH). **(B)** First Lyapunov coefficient on the primary curve and the GH fifth-order coefficient. **(C)** Stable and unstable periodic branches meet at the resolved cycle fold on *λ*_*I*_ = 0.84975; the horizontal coordinate magnifies the separation from Hopf. **(D)** The largest nontrivial Floquet modulus crosses one at that cycle fold. **(E)** Leading upper-half-plane equilibrium eigenvalues at the four conditions in Fig. 4; only a restricted window of the twelve-state spectrum is shown. **(F)** The *cubic local* unfolding in the linear crossing coordinates: gray denotes a stable equilibrium, pale green a stable mode-2 cycle, and hatching the additional positive mixed-amplitude saddle. Dashed teal rays are the two predicted secondary bifurcation boundaries; white circles are full-state shooting points in the displayed window. Unshaded white regions lack a stable state in the cubic truncation, without excluding a full-system attractor. The star marks the corrected torus. **(G)** Full-state periodic orbits at the two amplitude-0.04 secondary points; these are different parameter points. **(H)** An independently integrated 200-time-unit rate projection of the corrected saddle torus. **(I)** Positive modal-phase crossings over 4,000 time units, projected onto the second modal coordinate. Invariant-surface residuals and normal variational calculations establish the numerical torus and its saddle stability; projections alone do not.

**Fig. 6.**
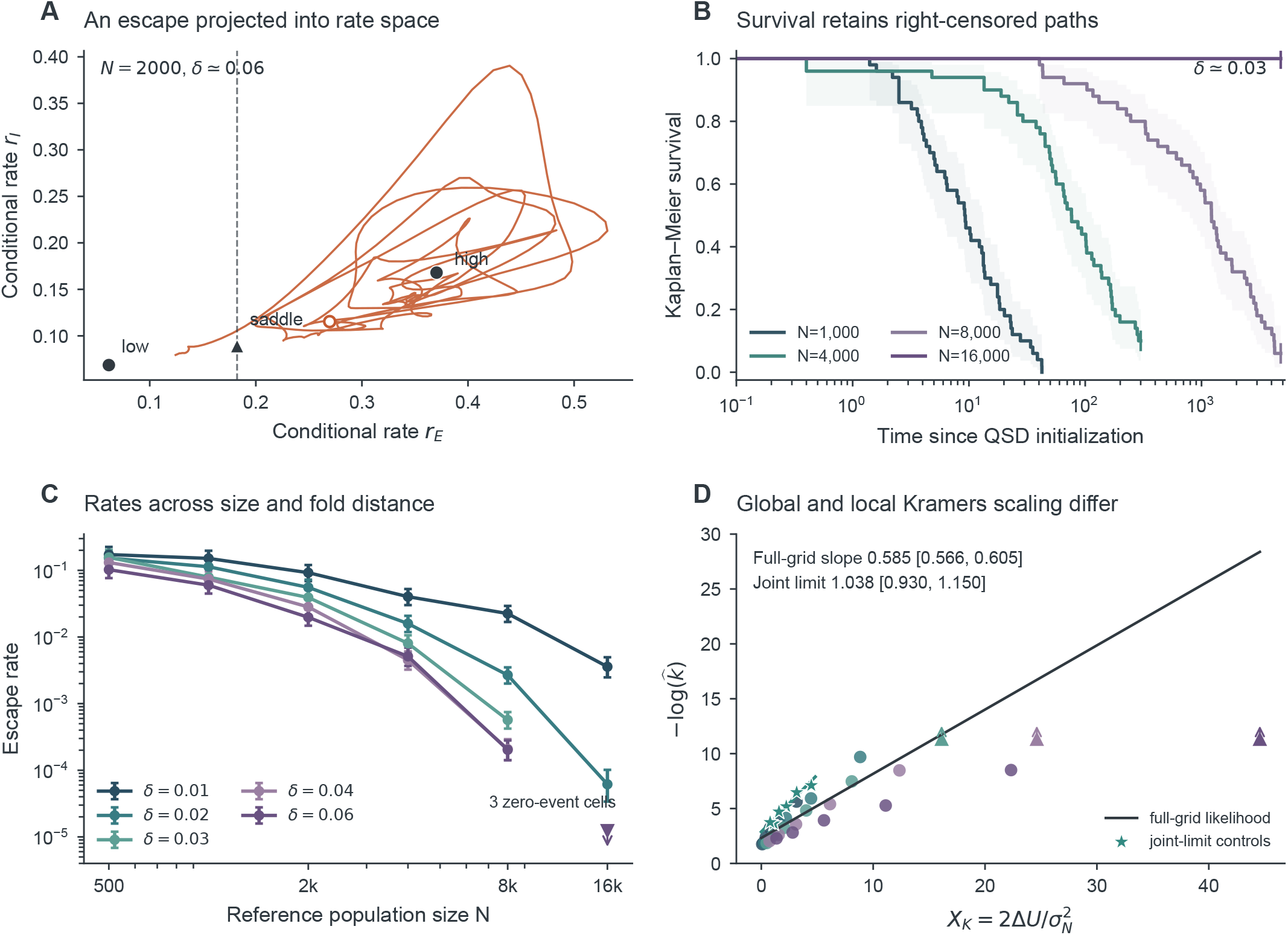
Finite-count escape approaches the local Kramers exponent only in the joint near-fold, weak-noise regime. **(A)** A 12-dimensional count-process path projected onto conditional E/I rates, with high and low equilibria, saddle, empirical-QSD initial state and saddle-rate detector threshold. **(B)** Kaplan–Meier curves at *δ* = 0.03 for four sizes, with Greenwood log-log nominal 95% bands and censoring ticks. **(C)** Event/exposure rates over the 30-cell admissible grid with exponential working-model profile intervals; downward triangles mark one-sided upper bounds for three zero-event cells. **(D)** Negative log rate against the local barrier exponent, including zero-event lower bounds. The axis notation 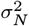 denotes the same fold-projected variance 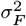, emphasizing its population-size dependence. The full-grid likelihood slope is 0 585; six separately labeled joint-limit controls give 1 038. The displayed fits have a free intercept and no prefactor offset. Each cell retains 50 paths; intervals are conditional on empirical reservoirs and the adaptive-horizon protocol in Supplementary Sec. 8.

### 5.10. Double-Hopf interaction: two periodic modes and a saddle torus

Because two Hopf curves intersect, neither oscillatory mode alone describes the local dynamics. Their coupled amplitudes predict a mixed-mode saddle; periodic branches and an invariant torus test that prediction in the full system. The curves meet at

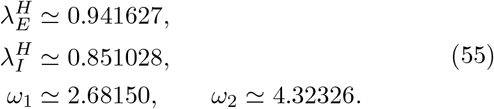

The remaining eight eigenvalues have real parts below −1.1273. The frequency ratio is approximately 1.61226. The minimum absolute homological detuning through degree five is 0.6020, separating this point from the low-order resonances relevant to the reduction. Full-precision coordinates and coefficients are retained in Supplementary Table S5; derived quantities use those values, not the rounded main-text matrices.

Using the full-state Jacobian A and Hessian B, choose ∥*q*_*j*_∥_2_ = 1 and 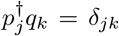, with conjugate eigenvectors included in the real center space. The eigenvector phases are fixed by making each largest component positive real. In a common original model time, the cubic normal form is

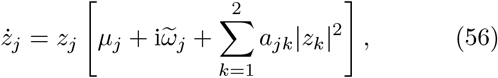

for *j* = 1, 2, to leading order in parameter displacement and cubic order in amplitude. With *δ****λ*** = ***λ*** − ***λ***^*H*^, the frequency detuning is

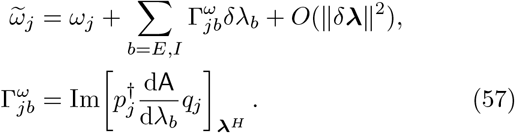

The total derivative follows the equilibrium branch. Its imaginary components are tabulated with the equilibrium-branch derivatives in Supplementary Sec. 11. The local parameter map, including equilibrium motion, is

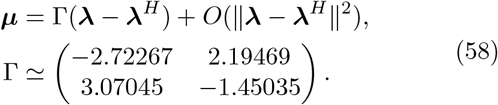

Its determinant is approximately −2.78989, so the two crossing directions are independent. The cubic coefficients are 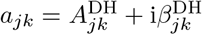, with

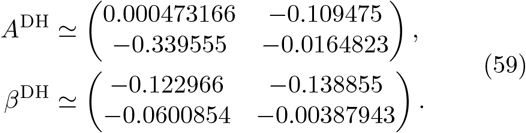

These are ordinary polynomial coefficients, not factorial-weighted coefficients or coefficients in separately rescaled modal times. In particular, the pure-mode convention in Eq. (48) gives *l*_1,*j*_ = Re *a*_*jj*_*/ω*_*j*_. The squared modal amplitudes *s*_*j*_ = |*z*_*j*_|^2^ satisfy

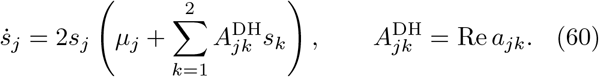

The signs 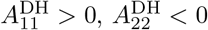 and det *A*^DH^ ≃ −0.0371805 624 gives case IV of the opposite-identify opposite pure-mode criticalities. The superscript DH identifies the real and imaginary cubic coefficient families. They are distinct from the synaptic rates *α*_*ab*_ and the transport rates *β*_*ab*_ introduced in the wave calculation. Writing 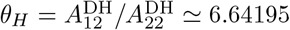 and 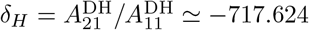 gives case IV of the opposite-self-coupling (“difficult”) classification^29^, Sec. 8.6. In this case, mode 1 exists for *µ*_1_ *<* 0 and is radially unstable; mode 2 exists for *µ*_2_ *>* 0 and is stable when *µ*_1_ *< θ*_*H*_*µ*_2_. Here *θ*_*H*_ is a coupling ratio, not a neuronal phase, and *µ*_*j*_ are real modal growth rates, not the complex eigenvalues used in spectral plots. The equilibrium is stable when both *µ*_*j*_ *<* 0. Where ***s*** = − (*A*^DH^)^−1^***µ*** is positive, the mixed-amplitude state has radial derivative 2 diag(***s***)*A*^DH^ and negative determinant. It is therefore a saddle in the amplitude system, not an attracting two-frequency state. The predicted secondary Hopf (Neimark–Sacker) boundaries are

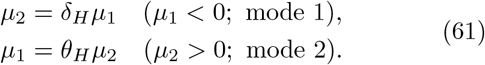

The negative cross-couplings describe mutual suppression of the modal growth rates; together with the opposite self-coupling signs, they determine the mixed state’s saddle character. These statements classify small-amplitude objects near this parameter point. Remote attractors, including possible large attracting tori, require separate analysis.

#### Coefficients of the interacting modes

The coefficients follow from a four-mode center embedding. Put 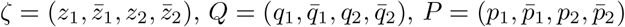 and Λ = i(*ω*_1_, −*ω*_1_, *ω*_2_, −*ω*_2_). With ordinary coefficients,

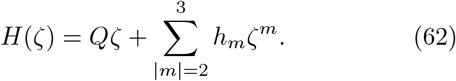

Matching 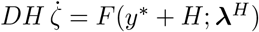 degree by degree gives

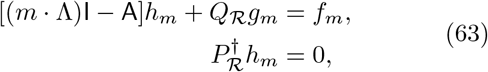

where ℛ contains the resonant outputs; both the added term and gauge constraint are absent for a nonresonant monomial. The forcing *f*_*m*_ contains already determined lower-order terms. Reality fixes the conjugate coefficients, while all four phase-equivariant cubic interactions are retained^30, 31^.

An independent explicit calculation uses Eq. (40). Define *R*(*σ*) = (i*σ*I − A)^−1^. The relevant quadratic coefficients are

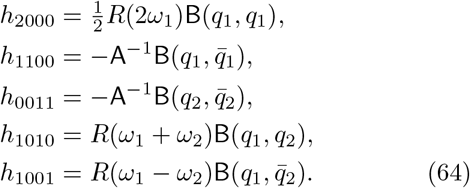

For a general third derivative C, the first self- and cross-couplings are

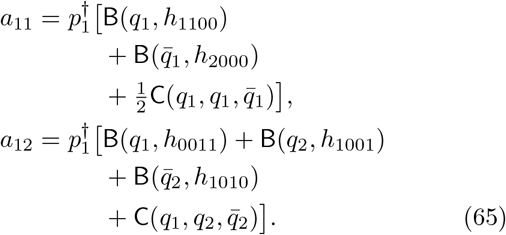

Interchanging modal labels gives *a*_22_ and *a*_21_. Although C = 0 for this quadratic neuronal field, the center embedding generates nonzero cubic coefficients. The polynomial and explicit calculations agree to 5.3 × 10^−16^.

#### Interacting periodic modes in the full system

Both secondary branches are realized in the full twelve-state system. Periodic shooting follows the two small-amplitude families to secondary bifurcations. Their critical multipliers cross the unit circle at finite angles, distinguishing interaction with the second oscillatory mode from a cycle fold. Continued slices verify the predicted crossing signs. Coordinates, multipliers and amplitude-dependent comparisons with the normal form are given in Supplementary Sec. 9; shooting and normal-form tests are described in Supplementary Sec. 11 and Supplementary Fig. S7.

#### Invariant torus and normal stability

The mixed-amplitude saddle predicts a two-frequency invariant object, not an attracting oscillation. To test that prediction, we solve a two-angle invariance equation for an embedding *Y* : T^2^ → ℝ^12^:

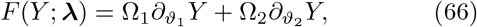

with two integral phase conditions and unknown frequencies^32^. Near (*λ*_*E*_, *λ*_*I*_) = (0.94202, 0.85156), the corrected frequencies are approximately Ω_1_ = 2.68112 and Ω_2_ = 4.32347. Angular-grid refinement and full-system evolution from the surface support the invariance calculation. Its section and spectrum visualize the resulting two-frequency motion (Fig. 5H,I).

Existence and stability require different checks. Eigen-functions of the full torus variational operator satisfy

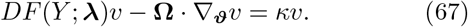

The two numerically resolved real normal exponents are *κ*_+_ ≃3.9093 × 10^−4^ and *κ* ≃ −4.1955 ×10^−4^. The associated eigenfunctions are normal to the torus after tangent projection; an integrated variational calculation confirms their signs. Thus the computed invariant object is a *saddle torus*. The unfolding predicts local coexistence with a stable mode-2 periodic state. Together, these resolution-converged calculations provide numerical evidence for an invariant saddle torus. The invariance and normal-variational discretizations, phase conditions and refinement tests are detailed in Supplementary Sec. 11.

### 5.11. Finite-rate passage and history dependence

Frozen bifurcation structure describes invariant states at fixed parameters. Biological change or imposed control instead occurs at a finite rate, so a trajectory need not track those states quasistatically. The competition between control rate and modal relaxation creates dynamic lag.

#### Tracking a moving state

To quantify tracking, for a stable frozen equilibrium define

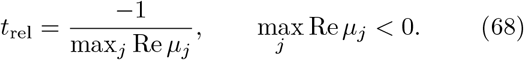

This is the leading modal relaxation time of the full coupled system. The local term 2*v*_*a*_*/τ*_*a*_ alone does not determine it. For a slowly moving equilibrium, ***z*** = ***y*** − ***y***^∗^(*λ*(*t*)) satisfies to first order

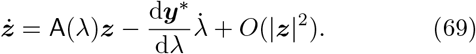

Relaxation slows near a stability boundary and quasistatic tracking can fail. The observed down/up tipping difference includes dynamic lag and the imposed ramp rate; it is not a stationary noise-driven dwell-time statistic.

The imposed matched-viability protocol decreases from 1 to 0.34 and then increases to 1, at 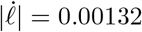. The observed down/up transition separation is 0.268884 under the specified rate-based detector. The passage protocol is detailed in Supplementary Sec. 6. Supplementary Fig. S5 compares the dynamic path with frozen equilibria and periodic-orbit extrema. The triangular protocol is imposed; its increasing limb is an external control operation, not a prediction of neuronal regeneration. It lies outside the irreversible thinning law 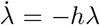 with *h* ≥ 0. The observed hysteretic separation combines frozen-state coexistence with finite-rate lag; it cannot be assigned to static bifurcation hysteresis alone.

Finite-rate control changes the trajectory through a deterministic landscape. Finite populations raise a different question: how sampling and stochastic emission affect the approximation and persistence of its states.

## 6. Finite populations: sampling and escape

The deterministic theory separates abundance from survivor state and identifies mechanisms with the same conditional dynamics. Finite populations introduce two different questions: how a quenched network approximates that mean field, and how fluctuations in emission can drive transitions between its attracting states. The first concerns sampling; the second concerns first passage.

### 6.1. Conditional observables in finite QIF networks

Let 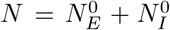 be the reference count and *n*_*a*_ the viable count in class *a*. Integer rounding gives the achieved fraction 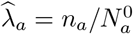. For an observation bin Δ*t*,

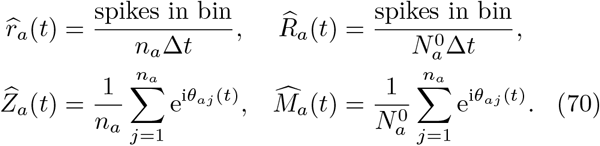

These definitions preserve the tissue identities exactly. Independent phases sampled from a fixed law satisfy

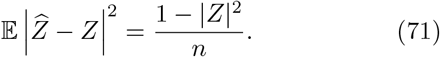

Recurrence introduces correlations, so this identity is a sampling benchmark, not a universal error law for a finite recurrent network.

The QIF comparisons support approach to the conditional theory while exposing a distinct source of persistent error: clipping Cauchy traits changes their limiting distribution. Clipping places probability mass at the cutoff and therefore defines a different continuum population. In the E-dominant-loss condition, stationary quadrature predicts an inhibitory-rate bias of about −0.0104 for the cutoff used. Increasing *N* cannot remove a change in the underlying trait law. Unbounded-trait simulations instead compare like with like and show decreasing errors in the primitive rate and phase observables. The complete comparisons and integration controls appear in Supplementary Fig. S2 and Supplementary Sec. 4. Trait sampling, count rounding and solver tests are specified in Supplementary Sec. 3; Supplementary Fig. S3 separates integration, clipping and sampling errors.

Nor does one finite-size exponent describe every observable. In the positive Cauchy tail, stationary single-cell firing grows as 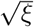, giving a logarithmically divergent quenched rate variance; phase moments remain bounded. Their sampling asymptotics need not agree. The measured finite-range slopes combine quenched sampling, temporal exposure and nonlinear network response, and provide no evidence for a universal pooled *N* ^−1/2^ law. This problem is separate from mesoscopic shot noise^33, 34^, to which we turn next.

## 7. Finite-count metastability near a fold

We now model fluctuations in presynaptic emission at fixed viable counts. Unlike the quenched QIF networks, this mesoscopic model has stochastic emission that can drive escape from a stable high-rate state. Projection onto the soft fold mode gives a local escape-rate prediction.

### 7.1. From viable counts to shared-source covariance

In the formulas below, 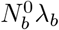 is the viable count; numerical trials use 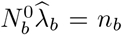 after rounding. Approximate population spike counts over a short bin by conditionally independent presynaptic processes

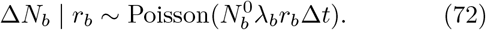

Each presynaptic count drives both outgoing pathways. The compensated tissue source increment 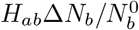 has conditional variance

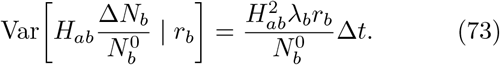

The diffusion approximation to the synaptic acceleration has a noise factor G_noise_ with two independent columns and entries

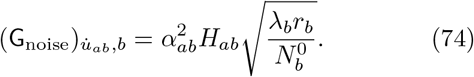

Other entries vanish. Its covariance is 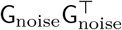; treating outgoing pathways as independent would incorrectly remove shared presynaptic covariance. This Poisson mesoscopic approximation separates emission fluctuations from quenched sampling^33–35^. Population-process escape theory provides the corresponding framework for metastability^36^; the count law is an additional modeling assumption, not a consequence of the deterministic OA reduction.

At a stable equilibrium, the Gaussian linear-noise covariance C_*y*_ solves

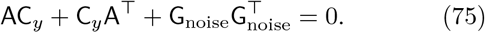

This provides a local small-fluctuation prediction, not an escape-rate estimate. For equal deterministic product *k*_*ab*_, the effective noise entering a pathway is proportional to 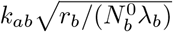, illustrating why equal conditional deterministic dynamics can have different finite-count variability.

### 7.2. Soft-mode projection and the local escape barrier

Projecting the emission noise onto the slow direction between the stable state and saddle gives a local scalar approximation. Using the left fold eigenvector,

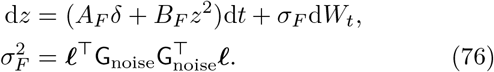

For the fitted local predictor, the noise factor uses fold-state rates and the fold left nullvector, together with each condition’s achieved viable fractions and reference counts. It does not substitute the high-state rates. State-dependent diffusion away from that local evaluation is an approximation error. For the additive gradient convention in Eq. (76), Kramers theory predicts

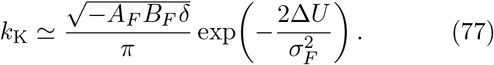

The factor two in the exponent follows from noise written as *σ*_*F*_ d*W*, not 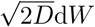. This convention is essential when comparing slopes^37, 38^.

With fixed class proportions, 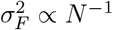, while Δ*U* ∝ *δ*^3/2^. The relevant joint variable is therefore *Nδ*^3/2^, not population size alone. The local fold approximation needs small *δ*; activated escape also needs a well-localized quasistationary distribution (QSD) and a dwell time long compared with the relaxation time. These conditions may conflict close to the fold at small *N*. A broad parameter grid can order rates correctly while failing to reproduce a unit exponent slope or the absolute prefactor.

### 7.3. First-passage survival and censor-aware inference

Testing an exponentially small escape rate requires accounting for trajectories that have not escaped by the observation horizon. Here “survival” refers to remaining in the high-rate regime, not to the viable fraction of neurons. We use event counts and total exposure, retaining right-censored paths and zero-event conditions. Kaplan–Meier curves and hazard diagnostics assess the event process^39^; the likelihood, risk-set definitions and uncertainty construction are given in Supplementary Sec. 8.

For condition *j*, define 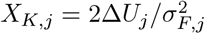 and fit

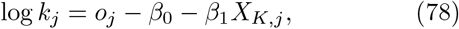

where *o*_*j*_ = 0 or the logarithm of the analytic Kramers prefactor. The likelihood uses each condition’s event count and summed exposure, including zero-event conditions. A unit slope is the asymptotic benchmark under this coordinate and diffusion convention. Residuals, profile intervals and stratified trajectory bootstrap intervals are retained. Bootstrap uncertainty conditional on estimated QSD reservoirs does not include all error in the finite-particle QSD approximation.

The first-passage experiment compares the shared-source count process with the fold predictor using quasi-stationary initialization and a persistent saddle-rate crossing rule. Unescaped trajectories contribute exposure, including in zero-event conditions. The count-process design and sensitivity tests are given in Supplementary Sec. 5 and Supplementary Fig. S4; reservoir construction, event detection and adaptive-horizon inference are detailed in Supplementary Sec. 8.

### 7.4. Local barriers require a joint limit

The fold reduction yields the barrier Δ*U* ∝ *δ*^3/2^ and projected variance 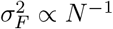. Consequently the Kramers exponent scales as 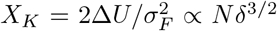, subject to both local center-manifold validity and weak projected noise. A long deterministic relaxation time alone does not guarantee this joint regime. We fit event and exposure data with a censored exponential likelihood, retaining zero-event cells, rather than regressing logarithms of observed escape counts alone^37, 38^.

For all 30 admissible grid cells, a free-intercept fit of negative log rate against *X*_*K*_ gives slope 0.5849 (profile 95% interval 0.5657–0.6045). A near-fold/weak-noise subset of 14 cells gives 0.9348 (0.8743–0.9973). Six additional joint-limit controls at *δ* = 0.005 or 0.0025 and *N* = 16000– 128000 give 1.0383 (0.9298–1.1499); including the analytic prefactor as an offset gives 1.0677 (0.9602–1.1786). Conversely, a timescale-separated selection without the locality restriction still gives slope 0.3637. The subset selection rules and fits are specified in Supplementary Sec. 5.

The broad grid fails to reproduce the local Kramers slope quantitatively, whereas the joint near-fold/weak-noise controls are consistent with the predicted unit exponent. Locality in deterministic fold distance and weakness of projected noise are both essential; timescale separation alone is insufficient. This agreement supports the exponent scaling, leaving the absolute prefactor and a global large-deviation principle unresolved. Comparisons must retain the same escape detector, quasi-stationary initialization and analysis horizon because these define the measured event process.

The count model treats viability as a class-level control. In tissue, loss may instead be localized, changing both the delayed source emitted at each location and the tissue-level weight of its survivor activity. We now turn to that field problem.

## 8. Viable source weighting in delayed neural fields

Spatially localized loss changes both the recurrent source emitted at each location and its contribution to tissue-level activity. We use delayed line and sheet fields to separate these effects, then compare intact and lesioned fields with identical pre-intervention histories.

### 8.1. The distributed-delay spatial operator

Synaptic filtering and axonal propagation contribute separate distributions of transmission times. The causal impulse response of the second-order synapse is

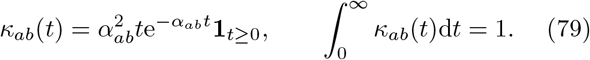

Its mean delay is 2*/α*_*ab*_ and its Laplace transform is 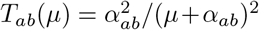. This synaptic filtering is distinct from propagation delay 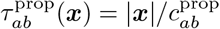. Distributed propagation affects homogeneous temporal modes as well as spatial patterns^40, 41^.

Suppressing pathway indices, write *ℓ*_ker_ for the kernel length and *c*^prop^ for the propagation speed. The kernel length is distinct from the matched viable fraction *ℓ*; with pathway indices restored it is *ℓ*_*ab*_. For the normalized line kernel 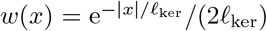, direct integration gives, with 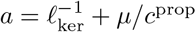,

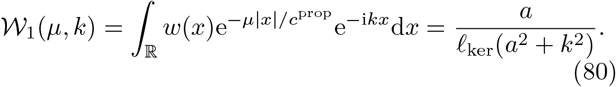

For the plane radial kernel 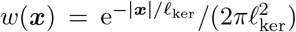, polar coordinates and the Bessel transform give

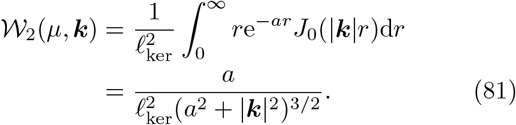

The integral representation initially requires Re *a >* 0. In this domain, 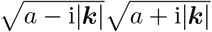 implements the analytic square root, with its value positive for real *a >* 0. In particular,

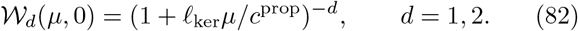

Setting wave number to zero does not erase distributed propagation delays. The undelayed homogeneous ordinary differential equation (ODE) is recovered when *c*^prop^→ ∞, or when the delay is explicitly omitted, rather than merely by taking *k* = 0.

For periodic numerical domains, the plane kernel is image-periodized before delay binning. Each image retains its actual path length. Minimum-distance wrapping of a kernel would define a different propagation model. The discrete weights are normalized; the pre-normalization DC quadrature defect and the omitted image-tail bound are reported separately. Refining spatial grid, integration timestep and delay-shell width addresses distinct error sources.

### 8.2. Characteristic matrix at a stationary homogeneous state

The transforms determine which temporal and spatial modes can destabilize a homogeneous stationary field. Eliminating the synaptic variables gives the neuronal characteristic system while retaining both kinds of delay. A perturbation proportional to e^*µt*+i***k***·***x***^ has synaptic increment

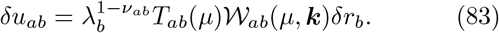

The neuronal characteristic equations are

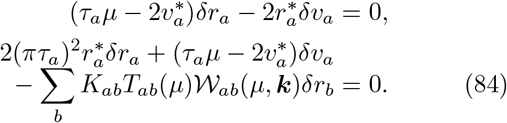

Their four-dimensional determinant is analytic in the stated Laplace domain except at eliminated synaptic poles. The one-dimensional transform is rational; the nonzero-wave-number radial transform has algebraic branch points under continuation beyond that domain. At a filter pole, the unreduced characteristic system must be used rather than treating a pole or a pole-zero cancellation as a root. The full field has infinitely many spatial modes.

At fixed wave number the one-dimensional exponential kernel, and the radial kernel at zero wave number, admit finite-dimensional realizations; general distributed-delay kernels need not do so. Multistart searches locate characteristic roots within a finite search domain. A root with positive real part establishes instability, even when the rightmost root has not been located.

The local observation identity can already be seen in a heterogeneous line field: structured survivor activity persists through a deficit while its tissue-level firing is attenuated (Supplementary Fig. S8). The paired maps compare two normalizations of one trajectory. To isolate recurrent reorganization, the sheet experiments below also compare that trajectory with a matched intact field.

### 8.3. A localized viability deficit in two dimensions

On a sheet, a localized deficit can also change propagation direction and spatial phase geometry. The delayed field is simulated on periodic spatial domains. The reference sheet is a periodic square of side *L* = 2*π*, with radial kernels

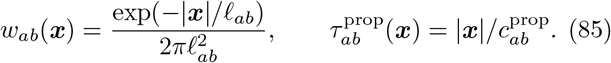

The spatial scales and speeds are

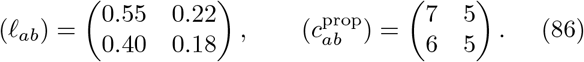

The class-specific lesion is 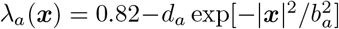, with (*d*_*E*_, *d*_*I*_) = (0.48, 0.30) and (*b*_*E*_, *b*_*I*_) = (0.58, 0.65). Widths have model-space units and velocities model-space per model-time units. A smooth prescribed perturbation seeds the field; the same deterministic construction is used for resolution comparisons.

Image periodization retains the actual length of each propagation path. The image-tail bound, quadrature defects and separate spatial, timestep and delay-shell refinements are given in Supplementary Sec. 10 and Supplementary Fig. S6.

The radial transform differs from the line-kernel transform:

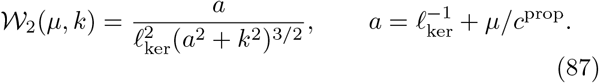

In particular *W*_2_(*µ*, 0) = (1 + *ℓ*_ker_*µ/c*^prop^)^−2^: a homogeneous perturbation retains the distribution of propagation times. At the homogeneous reference state with matched viability 0.82, a multistart characteristic-root search finds a growing pair at *k* = 0, *µ* ≃ 0.080564 + 2.19910i, and a larger searched growth rate near |*k*| = 1.5, approximately 0.20131, in the continuous-wave-number dispersion calculation. The latter magnitude is not an admissible mode on the 2*π* square lattice; the admissible *k* = 0 positive root already establishes instability of that stationary state. It does not determine which nonlinear pattern develops.

### 8.4. Local survivor activity and tissue-level patterns

Paired observations of one sheet trajectory show that substantial conditional activity can coexist with a much smaller local tissue contribution (Supplementary Fig. S9). This directly illustrates the observation identity, while a matched intact field is needed to isolate the lesion’s dynamical effect.

Spatial means need care when viability is heterogeneous. The displayed ⟨*r*_*a*_⟩_*x*_ averages local conditional rates over space. A whole-sheet survivor-normalized rate instead equals ⟨*λ*_*a*_*r*_*a*_⟩_*x*_/ ⟨*λ*_*a*_⟩_*x*_. Likewise, the whole-sheet survivor phase moment is ⟨*λ*_*a*_*Z*_*a*_⟩_*x*_/ ⟨*λ*_*a*_⟩_*x*_. These aggregate observations answer different questions. In particular, a spatial mean of phase-moment magnitudes differs from the magnitude of their spatial mean.

Short-horizon discretization tests and long-window summary diagnostics are reported separately in Supplementary Sec. 13. The central dynamical question is whether changing the source weights reorganizes the survivor state itself. The following matched intervention answers that question.

### 8.5. Local loss can increase whole-sheet activity

The decisive comparison is between fields that differ only in viability at the intervention. The intact and lesioned systems start from the same neuronal and synaptic state and the same emitted-source history. Signals already in flight retain their original weighting. Any subsequent difference is consequently a response to the altered viable source, not to a second intervention in the incoming history. The matched-state and emitted-history construction is specified in Supplementary Sec. 12.

A diagonal plane at matched viability 0.82 provides a persistent reference. With the deficit specified above, both the spatial mean conditional E rate and the whole-sheet tissue-level E firing rate increase (Fig. 7). Over the late observation window, the conditional mean rises from approximately 0.344 to 0.374, while the tissue-level mean rises from 0.282 to 0.301. The exact window, values and stability diagnostics are reported in Supplementary Sec. 14.2.

**Fig. 7.**
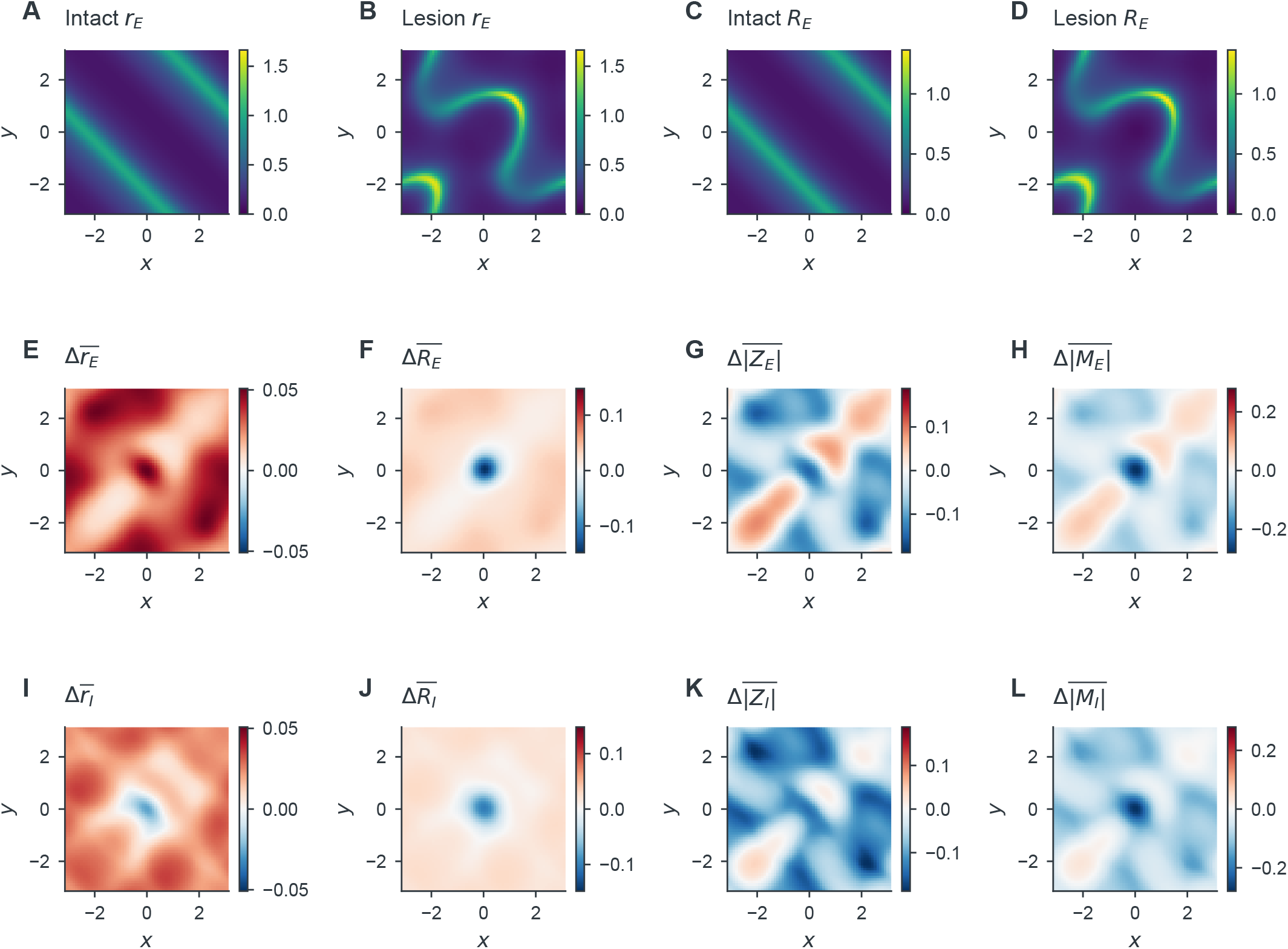
Localized neuronal loss increases whole-sheet E activity through recurrent reorganization. Intact and lesioned fields share the full pre-intervention state and emitted history. **(A–D)** Conditional and tissue-level E firing rates at *t* = 800, with common scales within each observable pair. **(E–H)** Lesion-minus-intact differences in the time means of local *r*_*E*_, *R*_*E*_, |*Z*_*E*_|, |*M*_*E*_| over 400–800; **(I–L)** the corresponding I quantities. Signed differences share symmetric scales within each observable column. Local mean magnitudes remain distinct from magnitudes of spatially averaged complex moments. The increase in mean E activity coexists with the local identity *R*_*E*_ = *λ*_*E*_*r*_*E*_.

This result separates two operations that would otherwise appear contradictory. At a fixed survivor state, *R*_*E*_(***x***, *t*) = *λ*_*E*_(***x***)*r*_*E*_(***x***, *t*) decreases when local viability decreases. In a recurrent field, however, that state does not remain fixed: loss changes the sources driving activity across the sheet. The resulting reorganization can outweigh local attenuation in the spatial average. Less local neuronal mass therefore need not imply less global activity. The I-population phase observations also change, demonstrating that the response is not captured by an E-rate scale factor alone.

### 8.6. Spatial organization beyond a plane

The delayed field also supports planar, radial and rotating activity under different initial conditions. These categories require different evidence: a radial image can be transient, whereas persistent rotation requires phase winding, tracked cores and angular rotation in the unfiltered firing rate. Selected conditions retain opposite-charge rotating cores over many cycles, including long continuations. Core motion and creation distinguish this activity from a single rigidly rotating spiral solution. The phase geometry, complete initial-condition study and refinement tests are in Supplementary Sec. 14 and Supplementary Fig. S10. For the main question, the persistent plane provides the controlled reference needed to isolate how a deficit changes an incident crest.

### 8.7. Crest timing and tissue amplitude respond differently

The matched planar field also separates a change in crest timing from a change in tissue amplitude. A depth– width study shares the intact state and emitted history at *t* = 800 across 21 conditions, including one intact reference. Arrival is measured along the physical propagation coordinate 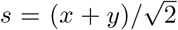, with *t*_*d*_(*s*) and *t*_0_(*s*) the lesioned and intact arrival times of the same incident crest. We distinguish

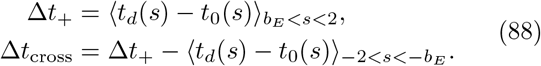

The downstream shift includes any upstream phase change caused by the simultaneous intervention. Subtracting that upstream shift gives the additional crossing shift. Both describe an oscillatory crest, not a causal information transmission time.

Across the tested deficits, the field exhibits arrival shifts, tissue-amplitude attenuation and local phase deformation, without a condition meeting the criteria for pinning or loss of the incident crest. One condition remains unresolved because its arrival fit lacks sufficient linearity. For the reference deficit, the downstream shift is approximately 0.156 time units, while the additional crossing shift is about 0.0412: treating the former as a transit delay would conflate two different responses. Crest tracking and qualification criteria are specified in Supplementary Sec. 12; the full parameter design, exact values and spatial panels appear in Supplementary Sec. 15 and Supplementary Fig. S11.

These heterogeneous fields reveal recurrent reorganization, but their evolving patterns need not be traveling invariant solutions. A coherent-wave calculation is needed to assign propagation speed and spectral stability to a defined object.

## 9. Coherent wavetrains and spectral stability

The heterogeneous two-dimensional fields show how viability affects evolving spatial patterns, but those patterns need not be coherent traveling solutions. We therefore return to a homogeneous one-dimensional field and solve directly for a wavetrain. The boundary-value problem determines its profile and speed; stability calculations then test perturbations of that same wave.

### 9.1. A comoving boundary-value problem

For a homogeneous one-dimensional wavetrain write ***y***(*x, t*) = ***Y*** (*ζ*), *ζ* = *x* − *c*_w_*t*, with period *L*. A source at displacement *s* and propagation delay *s /c*^prop^ is sampled at comoving coordinate

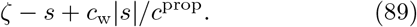

For spatial Fourier wave number *k*_*n*_ = 2*πn/L*, the temporal exponent is *µ*_*n*_ = −i*k*_*n*_*c*_w_. Thus synapses are eliminated exactly within a Fourier-collocation representation as

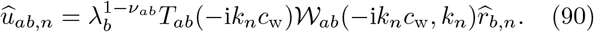

The four neuronal profiles solve

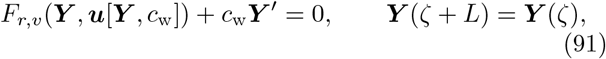

with an integral phase condition 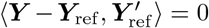. The wave speed is an unknown of the boundary-value problem, rather than an estimate from finite-time morphology. The continuation corrector solves these equations with an arclength condition; collocation resolution is checked independently, following neural-field methods for computing periodic and traveling solutions^7, 25, 27, 42^.

The boundary-value problem determines a traveling solution. Its stability requires linearizing the dynamics about that same solution, retaining the synaptic and propagation states eliminated during the profile calculation.

### 9.2. An exact transport representation for wave stability

For the line exponential kernel, a local transport system represents the distributed delay exactly and permits full-state linearization. For each pathway introduce two fields 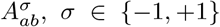, with 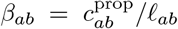 and 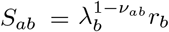:

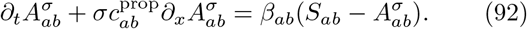

The average 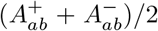 drives the alpha filter. Its Fourier–Laplace transfer function is

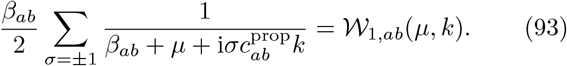

Together with four neuronal variables and eight alpha-filter variables, the eight directional fields form an exact 20-component partial differential equation (PDE) representation of the periodized one-dimensional model, with compatible transport histories. The representation does not replace the two-dimensional radial kernel by a one-dimensional exponential.

In the moving coordinate *ζ* = *x* −*c*_w_*t*, each neuronal and synaptic equation gains *c*_w_∂_*ζ*_; a transport equation has advective coefficient 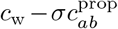. Linearizing all 20 components about the corrected wave and applying Fourier collocation yields a real 20*n* × 20*n* matrix. All its eigenvalues are computed. The translation vector must approximate a nullvector, providing a check beyond a small eigenvalue. Odd grids avoid the even-grid Nyquist derivative ambiguity. Numerical linearization and spectral convergence tests are given in Supplementary Sec. 10 and Supplementary Fig. S6.

If *µ* is a comoving eigenvalue, evolution over a full domain traversal gives

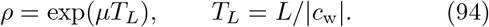

The computed wave has spatial wave number two, so its minimal observed period is *T*_*L*_/2. The reported monodromy over *T*_*L*_ uses two such cycles and returns the moving coordinate to the identity on all *L*-periodic perturbations. This spectrum tests perturbations on the fixed domain. Longer-scale modulations require the Bloch calculation below.

### 9.3. Viability-dependent waves and fixed-domain stability

To follow the wave as viability changes, we solve for its profile and speed using Eq. (91), with exact Fourier elimination of the synaptic and distance-delay filters within the collocation representation. A direct homogeneous-field trajectory supplies a seed, and pseudo-arclength continuation follows the family over matched viability approximately 0.70085–0.94745.

At *ℓ* = 0.82, the refined speed is *c*_w_ ≃ 0.871843. The wave has spatial wave number two on the 2*π* domain, hence wavelength *π* and minimal temporal period *π/*|*c*_w_|. Profile and spectral refinement are reported in Supplementary Sec. 9.

For a line exponential kernel, two counterpropagating transport fields per pathway exactly realize the propagation operator (Eqs. (92) and (93)). The resulting 20-component PDE permits linearization about the wave in its moving frame. The refined leading nontrivial pair is

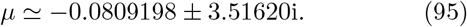

Its negative real part describes decay transverse to translation. The corresponding full-domain-traversal Floquet modulus is approximately 0.5581; this traversal spans two minimal periods. Twelve continued wave states also have no unstable nontrivial collocation eigenvalues. These spectra support numerical stability on the fixed periodic domain. The next calculation asks whether this wave belongs to a broader wavelength family and whether perturbations outside the original periodic cell are also damped.

### 9.4. Wavelength continuation and Bloch sidebands

The fixed-domain wave has primitive wavelength *P* = *π*, although its representation occupies *L* = 2*π*. In this subsection and its figure, *λ* = *λ*_*E*_ = *λ*_*I*_ = *ℓ* denotes matched viability, using *λ* rather than the loss-path coordinate *ℓ*.

We first vary the wavelength to establish a connected family, then test perturbations not restricted to the original cell. An independent adjoint phase-diffusion calculation checks the long-wave part of that spectrum.

#### A connected wavelength family

Let ***Y***_***\****_ denote the full 20-component wave profile. Pseudo-arclength continuation in (***Y***_***\****_, *c*_w_, *P*), with an integral phase condition, gives a connected wavelength family at matched viability *λ* = 0.82. Across the computed range 1.58913 ≤ *P* ≤ 6.21641, the speed increases from 0.603917 to 1.800822; at *P* = *π*, it is 0.871843. This establishes a connected family; stability is a separate property, tested next at its reference wavelength across viable fractions.

#### Perturbations beyond the periodic cell

A perturbation periodic on the doubled cell need not be periodic on one wavelength, and neither restriction covers every longer-scale modulation. To resolve these distinct spectral problems, let ***Y***_∗_(*ζ*) denote the full 20-component wave, with *ζ* = *x* − *c*_w_*t*, and write a perturbation as

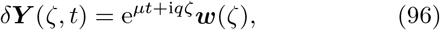

where ***w***(*ζ* + *P*) = ***w***(*ζ*) and −*π/P* ≤ *q* ≤ *π/P*. If ℒ_0_ is the linearized comoving transport generator, then the Bloch generator is obtained by replacing every spatial derivative by ∂_*ζ*_ + i*q*:

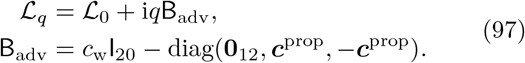

Here ***c***^prop^ lists the pathway propagation speeds in the order EE, EI, IE, II. The advective matrix B_adv_ is distinct from the state Hessian used in the normal-form calculation. The first twelve entries comprise the neuronal variables and both alpha-filter stages; the remaining entries are the two directional transport fields for each pathway. Thus the Bloch shift acts on all dynamical components, including those without laboratory-frame advection. Because the coefficients are real, complex conjugation maps ℒ_*q*_ to ℒ _−*q*_, so 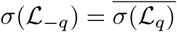. In particular, ℒ_*q*_ and ℒ _−*q*_ have the same spectral abscissa, which permits restricting the stability scan to nonnegative *q*. For the two-copy wave, cell folding gives

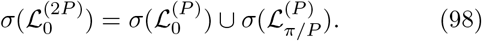

Consequently a doubled-cell periodic calculation samples only the periodic and antiperiodic sectors of the primitive cell, rather than its complete sideband spectrum.

#### An independent long-wave stability check

The translation-derived branch determines whether slow phase modulations grow or decay. Adjoint perturbation theory evaluates its curvature independently of a fit to sampled Bloch eigenvalues. Translation invariance supplies ℒ_0_***ϕ*** = 0 with ***ϕ*** = ∂_*ζ*_***Y***_***\****_. For a simple translation eigen-value, choose an adjoint nullvector ***ψ*** satisfying ⟨***ψ, ϕ*⟩** = 1, using the period-averaged complex *L*^2^ inner product. Perturbation of this eigenpair yields

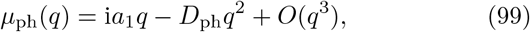

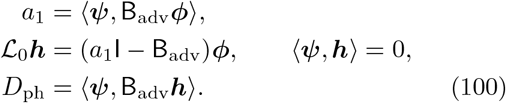

Here *a*_1_ is the linear phase coefficient in the comoving frame, not the wave speed. Along a differentiable wavelength family, the temporal angular frequency is *ω*(*k*) = *kc*_w_(*k*) with *k* = 2*π/P*, so

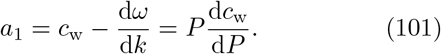

This links the small-*q* spectrum directly to the nonlinear wavelength continuation. A positive *D*_ph_ damps sufficiently long-wavelength phase modulations of this simple branch. Other spectral branches must still be checked independently.

#### Sampled sideband damping and phase relaxation

For *P* = *π*, complete finite-dimensional Bloch spectra at *λ* = 0.705, 0.76, 0.82, 0.88, 0.94 show no resolved positive nontranslation eigenvalues over the sampled Brillouin zone (Fig. 9). At *λ* = 0.82, the co-periodic transverse pair has real part −0.0809198, whereas the zone-edge spectral abscissa is −0.0952044. The adjoint phase-diffusion coefficient increases from 0.0329848 at *λ* = .705 to 0.156090 at *λ* = .94, with *D*_ph_ = 0.130413 at *λ* = .82. Direct small-*q* spectral fitting agrees within 2.4 ×10^−8^. The increasing phase-diffusion coefficient identifies a viability-dependent change in long-wave relaxation on this branch. Fourier and Floquet-number refinement support numerical sideband damping over the sampled Bloch spectrum. The wavelength continuation, Bloch sampling and adjoint calculations are detailed in Supplementary Sec. 16; Supplementary Fig. S12 reports their convergence. A continuum spectral-stability theorem remains outside this calculation.

**Fig. 8.**
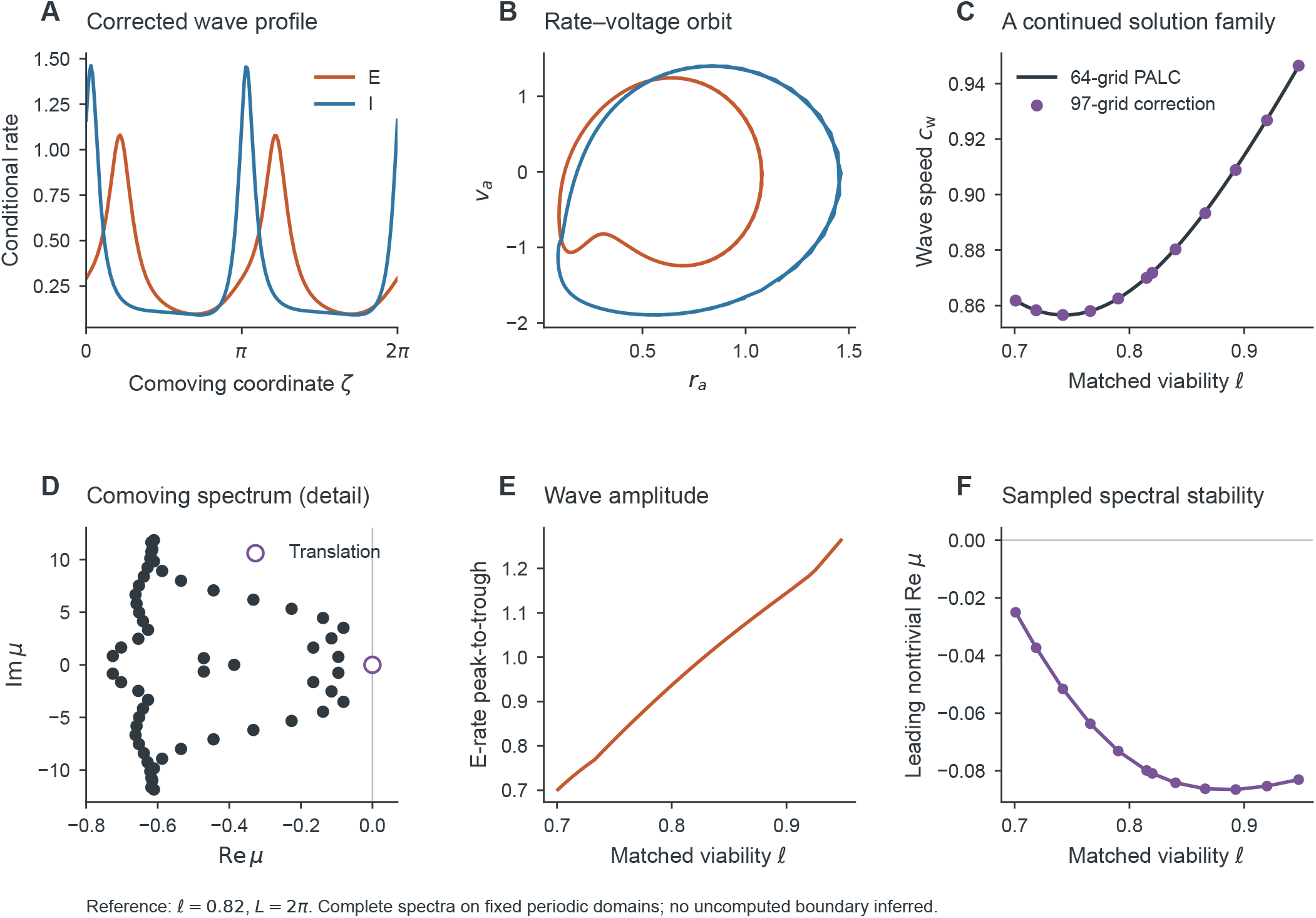
The continued wavetrain changes speed with viability and is numerically stable at the sampled fixed-domain states. **(A**,**B)** The 193-point corrected E/I wave profile and rate–voltage orbit at matched viability 0.82. **(C)** Wave speed along the 50-point 64-grid pseudo-arclength family, with 12 independently corrected 97-grid samples. **(D)** The rightmost portion of the complete 193-grid comoving spectrum; the open symbol identifies the translational neutral mode. **(E)** Continued 64-grid E-rate peak-to-trough amplitude. **(F)** Leading nontrivial eigenvalue real part at the 12 stability samples. The numerical classification concerns *L* = 2*π*-periodic perturbations, including the primitive-cell periodic and antiperiodic sectors, not all Bloch sidebands. Velocities are in model-space units per model time, without physiological calibration.

**Fig. 9.**
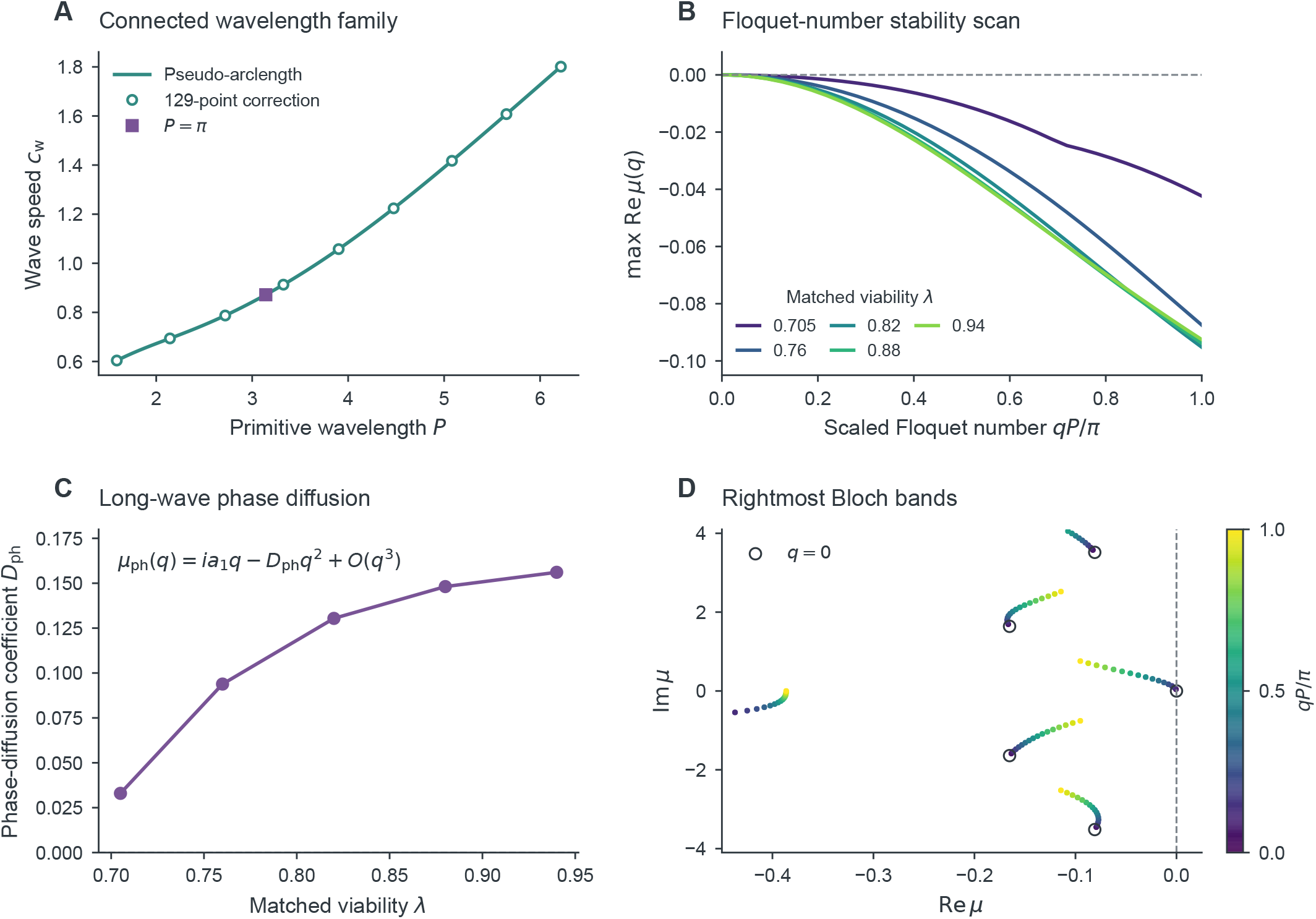
The coherent-wave family exhibits sampled sideband damping and viability-dependent long-wave phase relaxation. **(A)** Phase-conditioned pseudo-arclength continuation at matched viability *λ* = 0.82; open circles show independent 129-point profile corrections. The square marks *P* = *π*. **(B)** Rightmost eigenvalue real part versus scaled Floquet number for five viabilities at *P* = *π*; the neutral point at *q* = 0 is the translation mode. Each point is computed from a complete 980-dimensional spectrum. **(C)** Long-wave phase-diffusion coefficient from 97-mode spectral fits, independently checked by the adjoint formula. **(D)** Rightmost Bloch bands at *λ* = 0.82, colored by Floquet number; open circles denote co-periodic eigenvalues. The displayed spectral window is restricted to Re *µ >* −0.45 and |Im *µ*| *<* 4.1, while the stability tests use the complete spectra.

## 10. Discussion

The theory separates two questions that an activity measurement alone cannot resolve: how many neurons contribute, and how those neurons behave. Viable mass affects both the observation and the recurrent input that generates the observed state. Yet different mechanisms can preserve that input exactly. These two results explain why neuronal loss need not produce a proportional activity decrease, and why a change in activity need not identify neuronal loss.

### 10.1. Population mass is neither excitability nor an observation gain

The factorization *q*_*a*_ = *λ*_*a*_*f*_*a*_ assigns different mathematical roles to abundance and conditional state. State-independent thinning changes mass without selecting among states or quenched traits within a class. On the analytic invariant manifold, normalization and conditional OA/MPR reduction commute. The surviving neurons therefore retain the conditional state law, while their recurrent input changes with viable presynaptic mass. Viability is neither an intrinsic-current parameter nor a scale factor applied after solving the network dynamics.

The distinction is also observational. Conditional firing *r*_*a*_ and phase concentration *Z*_*a*_ describe survivors; *R*_*a*_ = *λ*_*a*_*r*_*a*_ and *M*_*a*_ = *λ*_*a*_*Z*_*a*_ describe their contribution relative to the reference population. High survivor firing can coexist with a small tissue-level firing rate. Likewise, a concentrated conditional phase distribution can have a small tissue-level coherent moment. Phase concentration around a resting state is not itself rhythmic synchronization.

Exactness here means invariance under the specified analytic assumptions, not attraction from every initial distribution. Attractiveness results and descriptions beyond the OA manifold address related but distinct questions^10, 11^. Selective survival changes the conditional law itself: the failure-of-tangency example shows why a scalar viable fraction cannot generally absorb state-dependent selection.

### 10.2. A generative distinction need not be identifiable

The constant-product conjugacy is stronger than a stationary-rate match. Corresponding synaptic coordinates give identical conditional neuronal trajectories, equilibrium spectra, periodic stability and deterministic forced responses. Thus viability, pathway integrity and compensation can remain structurally indistinguishable even with noiseless conditional recordings. The density–coupling gauge gives a second exact redundancy. Increased precision within an unchanged observation channel cannot break either symmetry.

Identifiability depends jointly on the parameterization, observation operator, perturbations, structural constraints and measurement uncertainty^13–15^. Frequency-resolved responses distinguish alternatives that match only a stationary summary, but not mechanisms related by the exact conjugacy. Tissue-level measurements can supply mass information when they observe a population whose viable fraction differs. Independent cell counts, pathway measurements or calibrated coupling constraints can supply information absent from conditional dynamics. The local design calculation identifies promising observation directions; practical inference from neural recordings still requires a likelihood and an explicit measurement model.

### 10.3. Selective E/I loss changes the dynamical landscape

Changing E and I mass alters different recurrent pathways. Its consequences extend beyond the amplitude of an existing state to the arrangement of equilibria, oscillations and coexistence regions. The generalized-Hopf point changes Hopf criticality and organizes a narrow cycle fold. At double Hopf, two oscillatory modes interact through cross-coupling; their mixed-amplitude saddle corresponds to a numerically computed saddle torus in the full system. These structures connect viable source strength to the organization of the network’s possible dynamics.

The ordering of class-specific Hopf transitions changes across parameter sets supporting an oscillatory intact state. The reference ordering is a property of the surrounding recurrent parameters, not a universal hierarchy of E and I vulnerability. Finite-rate control adds lag to this frozen landscape, combining state coexistence with incomplete relaxation. That history dependence is distinct from stochastic escape and from a biological claim of regeneration.

### 10.4. Two different finite-population problems

A finite quenched QIF network approximates a deterministic conditional law; a mesoscopic emission process fluctuates around and can escape its attractors. These are different limits, not a single universal finite-*N* correction. For the first problem, clipping Cauchy traits changes the continuum distribution, so increasing population size cannot remove cutoff bias. Heavy-tailed firing contributions and bounded phase moments also have different sampling behavior. The comparisons support observable- and sampling-dependent convergence, without a universal pooled *N* ^−1/2^ law.

For the second problem, shared presynaptic counts determine the covariance projected onto the soft fold mode. The local barrier scales as *δ*^3/2^ and the noise variance as *N* ^−1^, making *Nδ*^3/2^ the relevant asymptotic combination. The broad-grid departure from the unit Kramers slope is informative: deterministic locality and weak projected noise are both required. The joint-limit controls support the exponent, while the absolute prefactor and global escape action remain open. The Poisson emission approximation is separate from microscopic stochastic QIF reductions^43^ and from large-deviation treatments of other neuronal population processes^36^.

### 10.5. Local loss and global propagation

Spatial viability weights the delayed source emitted at each location and the tissue-level observation made there. Distributed propagation times remain even at zero wave number; a spatially homogeneous perturbation is not generally an undelayed one. The exact line-kernel transport representation and the radial sheet transform preserve this distinction.

The matched lesion experiment makes the recurrent effect particularly clear. At a fixed survivor state, lower viable fraction reduces local tissue output. Once the recurrent field adjusts, however, that state is no longer fixed: the tested lesion increases both mean conditional E firing and whole-sheet tissue-level E firing. Local mass loss and global activity can therefore change in opposite directions. Identical pre-intervention states and emitted histories are essential to attributing this reorganization to the lesion.

The spatial analyses distinguish evolving morphology from coherent propagation. Phase geometry, persistent cores and unfiltered angular rotation support rotating activity in selected conditions, while radial and some planar patterns are transient. Moving and disappearing cores need not constitute a rigid spiral. Across the tested deficit range, crest timing, tissue amplitude and local phase deform without qualified pinning or propagation failure. Stronger propagation effects found for other forms of field heterogeneity^44, 45^ remain possible outside this range. Because intervention can shift an incident crest upstream as well as downstream, its phase-equivalent timing shift is not a causal information delay.

A continued wavetrain supplies a sharper invariant object. Its boundary-value problem determines profile and speed; wavelength continuation and Bloch sidebands test perturbations beyond the original periodic cell. Positive adjoint phase diffusion agrees with long-wave spectral damping at the sampled viable fractions. These calculations establish numerical stability over the sampled spectra; continuum spectral bounds, disconnected wave families and two-dimensional transverse stability remain further questions^27, 28, 46^.

### 10.6. Extensions require explicit changes of assumptions

Refining the two-class population structure would separate laminar and cell-type-specific vulnerability from changes in interpopulation coupling. Transcriptomic or cellular measurements could constrain vulnerability priors, as motivated by spectrolaminar modeling^47^, while within-class selection would require an evolving conditional trait law. Adaptation, electrical coupling and non-Cauchy heterogeneity alter the cellular or distributional assumptions and require corresponding reduction arguments.

Replacing homogeneous geometry by structural connectomes and distributed-delay tensors^18^ would distinguish pathway loss from propagation changes. Myelination-dependent conduction would enter velocity or delay distributions separately from integrity^48^. These are extensions of a nondimensional mechanism model, not a clinical calibration.

Finally, *M*_*a*_ is not an EEG, MEG or LFP forward model. Those measurements require synaptic or transmembrane currents, source geometry and orientation, conductivity, volume conduction and sensor mixing. Higher-order spectra may constrain nonlinear interactions^49^, but statistics of identical conditional trajectories cannot identify their distinct biological causes without additional observation or structural information.

## 11. Conclusion

Neuronal loss changes both the amount of tissue contributing to activity and, through recurrent source strength, the dynamics of surviving neurons. Its effect is not generally a multiplicative reduction of neural activity. Conversely, distinct viability, pathway-integrity and compensation mechanisms can generate exactly the same conditional deterministic dynamics. Inferring neuronal loss from activity therefore requires an observation model and independent structural constraints.

## Acknowledgments

This work was supported by the National Key R&D Program of China (2024YFE0215100), the Brain Science and Brain-Inspired Intelligence Technology National Science and Technology Major Project (2022ZD0208500), and the National Natural Science Foundation of China (W2411084). Additional funding was provided by the Key Research and Development Projects of the Science and Technology Department of Chengdu (2024-YF08-00072-GX) and the Chengdu Science and Technology Bureau Program (2022-GH02-00042-HZ). Institutional support was received from the University of Electronic Science and Technology of China (UESTC) through the CNS Program (Y0301902610100201).

## Conflict of interest

The authors declare no conflicts of interest.

## Data and code availability

Simulation and analysis code, parameter configurations and numerical outputs are provided in the computational repository on *GitHub*. All data are simulation outputs; no human or animal data were collected.

## Supplementary Materials

## Supplementary Methods and Results

### 1. Numerical design

This supplement specifies numerical approximations, condition-level sampling, integration and continuation controls, and uncertainty calculations. The main article contains the mathematical derivations and principal results. Configurations and source routines specify the parameters, initial conditions and pseudorandom seeds. Numerical tables accompany the source code; the computational documentation identifies the retained and regenerable trajectories.

### 2. Continuation, normal forms and observation design

The full deterministic state has four neuronal and eight synaptic variables. Log-rate equilibrium continuation enforces positive rates, while all stability assignments use the complete state Jacobian. A singular-value null tangent, predictor, augmented hyperplane corrector and step-halving failure handling implement pseudo-arclength. No boundary is obtained by smoothing the exploratory grid. Fold-nullvector normalization and critical-pair residuals are evaluated at each continuation point. High-order coefficients follow the polynomial invariance equations in the main article, with ∥*q*∥_2_ = 1, *p*^†^*q* = 1, ordinary polynomial coefficients, and reported *l*_1_ = Re *g*_21_*/ω, l*_2_ = Re *g*_32_*/ω* in time *s* = *ωt* [1, 2].

The shooting unknowns include the initial state and log period. DOP853 integrates the orbit, full variational matrix and parameter sensitivity. The phase section is updated locally to avoid tangencies to a fixed global section. The subcritical cycle fold is checked by a neutral-lifted multiplier condition and 18 tolerance controls. Table S1 summarizes continuation accuracy and sampling; condition-level values are given in Secs. 7 and 9.

**Supplementary Table S1.** Continuation accuracy and sampling. Complete tolerance settings are supplied with the configurations.

| Calculation | Result or verification |
| --- | --- |
| Exploratory equilibrium count | 841/841 grid counts agree with continued folds. |
| High-state fold, $\lambda_I = 0.9$ | $\lambda_E = 0.538519620099$; full-Jacobian smallest singular value $1.57 \times 10^{-14}$. |
| Matched physical periodic branch | 90 computed points; intact period 2.801836, amplitude 0.666636. |
| Generalized Hopf | (0.9382654414, 0.8468556406); $l_2 = -0.00646343$; homological residual $< 9 \times 10^{-16}$. |
| Cycle fold, $\lambda_I = 0.84975$ | $\lambda_E = 0.9405981518$; augmented residual $1.51 \times 10^{-14}$. |
| Cycle-fold distance from Hopf | $5.80277 \times 10^{-7}$; quintic prediction differs by 0.0405%. |
| Admitted parameter ensemble | 128/157 proposals; 87 complete regular path triples, 38 topology changes, 3 unresolved members. |
| Comparable path ordering | Faster-I first 21/87; faster-E first 66/87. Among all admitted members these are 21/128 and 66/128, with the remaining category explicit. |
| Observation design | Nine fractional parameter directions; rank 5 with summaries/relaxation, rank 8 after frequency responses; one exact density-coupling gauge. |

The ensemble samples independent relative perturbations within ± 10% of reference coupling magnitudes, trait widths and centers, synaptic rates and density ratio. Admission requires the specified intact oscillatory regime. Pathwise regular sample counts are 104/99/99; their marginal distributions differ from the 87-member complete-path ordering sample. These crossings are local equilibrium Hopf events, not a global search of coexisting attractors.

The design matrix uses signed fractional reference parameter coordinates and fixed observation scales: 0.1 for rate/response rows and one for eigenvalue rows, with identity working covariance. Additional responses are evaluated at angular frequencies 0.5, 1, 2 and 4. The exact null direction 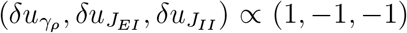 is reported. Here *u*_*γ*_, *u*_*J*_ and *u*_*J*_ are the fractional parameter coordinates, not absolute changes in density or coupling. No singular value is artificially floored before interpretation [3, 4]. Table S2 gives the complete singular-value spectra for both observation sets, including the null directions.

**Supplementary Table S2.** All nine observation-design singular values at fractional central difference step 10^−4^. Rank uses threshold 10^−5^. Zeros are retained here but cannot be displayed on the logarithmic figure axis.

| Index | Summary and relaxation | With frequency responses |
| --- | --- | --- |
| 1 | 6.1344831 | 8.1820890 |
| 2 | 0.8705800 | 1.0325293 |
| 3 | 0.4186408 | 0.5106227 |
| 4 | 0.3060790 | 0.4693688 |
| 5 | 0.2252679 | 0.2756589 |
| 6 | 0 | 0.0907199 |
| 7 | 0 | 0.0575651 |
| 8 | 0 | 0.0272700 |
| 9 | 0 | $4.7627 \times 10^{-9}$ |

For the response-augmented design, halving the fractional difference step from 2 × 10^−4^ to 10^−4^ and 5 × 10^−5^ reduces the ninth singular value from 1.9042 × 10^−8^ to 4.7627 × 10^−9^ and 1.1825 × 10^−9^. This is finite-difference residue along the exact gauge, not a ninth identifiable direction. Both designs retain their stated ranks over these three steps.

**Supplementary Fig. S1.**
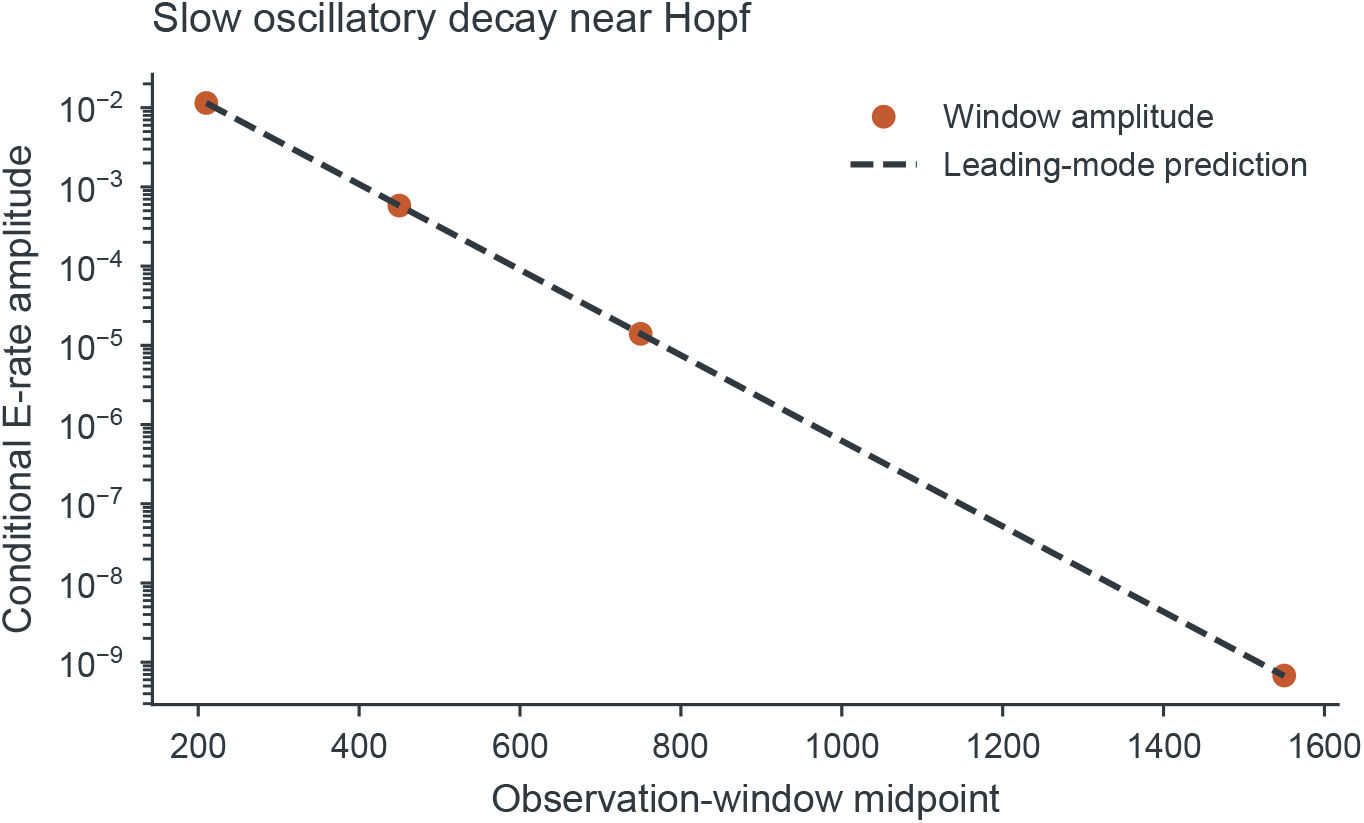
A near-Hopf oscillatory transient at matched viability 0.55. Points show the peak-to-trough conditional E-rate amplitude in successive windows of the same trajectory. The dashed prediction uses the leading linear decay exponent and is anchored to the first amplitude. The decreasing amplitude is consistent with asymptotic decay, not a stable periodic orbit.

### 3. Direct microscopic validation and numerical error controls

Direct simulations use the quadratic integrate-and-fire (QIF) microscopic model underlying the conditional Montbrió–Pazó–Roxin (MPR) equations [5]. All parameters are loaded from the single reference configuration. As in the main article, 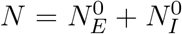 is the total reference count and *n*_*a*_ is the viable count in class *a*. With *γ*_*ρ*_ = 0.25, reference counts are 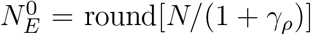 and 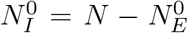; viable counts are 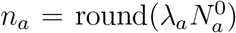. The achieved fraction 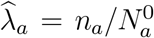 is used consistently in the deterministic comparison, tissue-level firing rate 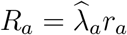, tissue-level coherent moment 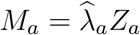, and projected noise. No minimum-count floor is imposed. Table S3 separates population-size comparisons from integration and sampling controls.

The six viability pairs are (0.4, 0.4) for matched low viability, (0.5, 0.8) for E-dominant loss, and (0.8, 0.5) for I-dominant loss. The other conditions are intact (1, 1), near-fold (0.56, 0.9), and near-Hopf (0.56, 0.56).

**Supplementary Table S3.** Direct-QIF design and actual run counts. Every run has 100 units of burn-in and 400 units of observation. Controls are separate labeled runs even where a parameter value coincides with a size-study setting.

| Design | Sizes or control values and replication | Runs |
| --- | --- | --- |
| Clipped-trait size study | $N = 400, 800, 1600, 3200, 6400, 12800$; six conditions; 20 seeds at each of the first five sizes, 10 at the largest | 660 |
| Clipped-trait timestep | $N = 1600$; $\Delta t = 0.01, 0.005, 0.0025, 0.00125$; two conditions, 10 paired seeds | 80 |
| Trait clipping | $N = 1600$; cutoff 20, 30, 50, 80; two conditions, 10 seeds | 80 |
| Trait sampling | $N = 1600$; three sampling laws; two conditions, 10 seeds | 60 |
| Crossed numerical resolution | $N_E^0 = 400, 800, 1600$ crossed with $N_I^0 = 100, 200, 400$; one condition, 10 seeds | 90 |
| Unbounded-trait size study | Same six total sizes; three conditions, 10 seeds per cell | 180 |
| Unbounded-trait timestep | $N = 1600$; $\Delta t = 0.01, 0.0025$; two conditions, 10 seeds; reference 0.005 runs above | 40 |
| Total | Network model-time exposure = 595000 | 1190 |

The two control conditions are E-dominant loss and intact; the adaptive unbounded study additionally includes matched low viability. The default trait-clipping cutoff is 80; timestep and sampling refinements are specified below. Observation was configured to extend if necessary to contain 100 mean-field cycles; all completed records used 400 units, with at least 100.546 cycles in oscillatory comparisons. Between-realization variability, first–second-half differences, basin occupancy, and phase-aligned versus unaligned oscillatory errors are recorded separately. The independent measured channels are spike-count rate and complex phase moment; tissue-level quantities and voltage reconstructed from *Z* are not additional independent validation channels. The unweighted primitive mean-error summary is

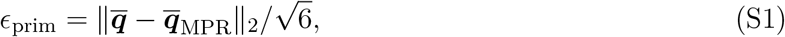

where the six entries of ***q*** are the two conditional rates and the real and imaginary parts of the two phase moments; all quantities use model units.

#### Sampling and refinement protocol

All runs use 100 model-time units of burn-in and at least 400 observation units, with an explicit oscillation-window check. The clipped-trait integration timestep is 0.005, phase statistics are sampled every 0.05, and uncertainty is separated into 20-unit temporal blocks and between-realization variation. Supplementary controls compare steps 0.01, 0.005, 0.0025, and 0.00125, Cauchy clipping levels 20, 30, 50, and 80, and three sampling designs: random reference traits, reference quantiles followed by thinning, and direct survivor quantiles.

The clipped-trait solver uses the midpoint theta step and exact homogeneous synaptic propagation with end-of-step impulses. The synaptic splitting is not assigned the midpoint method’s nominal order: successive measured errors are used instead. Initial phases are sampled from the mean-field marginal wrapped-Cauchy law independently of excitability; this is not an exact stationary joint phase–trait initialization.

Clipping is winsorization with endpoint atoms, not conditional truncation. Independent stationary quadrature for the clipped law isolates a continuum bias: at cutoff 80, E-dominant loss changes the limiting inhibitory rate by −0.0103800 relative to unbounded MPR. The unbounded quadrature recovers the MPR stationary rates and phase moments to approximately 10^−12^. The unbounded-trait solver uses the exact frozen-current Riccati flow in projective coordinates, counting all complete spike rotations, including multiple spikes within one step. Positive-, negative-, and zero-current single-neuron solutions are checked analytically. Synaptic/recurrent splitting remains approximate. Cross-solver complex-phase differences decrease from 4.54 × 10^−4^ at step 0.01 to 1.29 × 10^−5^ at 0.00125.

With condition-specific intercepts and proportional class counts, separate coefficients of log *n*_*E*_ and log *n*_*I*_ are unidentifiable in the unrounded design (rank 7 for 8 columns). Rounding only creates numerical full rank with condition number 2.67 × 10^4^. The crossed control instead varies computational sample resolution with representative-neuron weights that hold macroscopic *γ*_*ρ*_ fixed; it is not a physical reference-density manipulation. The common finite-range primitive-error slopes are −0.311 (clipped-trait) and −0.419 (unbounded-trait), with seed-cluster bootstrap 95% intervals [−0.363, −0.262] and [−0.490, −0.349]. Neither is an established asymptotic exponent. In particular, unbounded Cauchy traits and single-neuron rates proportional to 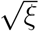 produce a logarithmically divergent second quenched rate moment. Conditional Poisson *N* ^−1/2^ count noise therefore does not imply an ordinary universal quenched central-limit rate. The projected count-noise predictor also fails to outperform the minimum-count predictor in held-out stable conditions (Supplementary Fig. S3).

### 4. Finite-QIF comparisons and sampling results

We first ask how a finite QIF population approximates the conditional theory, holding neuronal loss fixed before each realization. The comparison must separate sampling of the survivor distribution from changes to that distribution caused by numerical trait cutoffs. Otherwise an apparent finite-size error can persist even in the continuum limit.

#### 4.1. Finite-population observables

Finite-network observables and their independent-sampling benchmark are defined in main-text Eqs. 70 and 71, using achieved viable fractions. Recurrent finite networks need not satisfy the independence assumption. This formula is a sampling benchmark, not a general convergence theorem. Temporal autocorrelation similarly motivates block rather than pointwise uncertainty.

#### 4.2. Comparing finite networks with the conditional theory

The microscopic study includes 970 clipped-trait phase-based QIF realizations and 220 unbounded-trait QIF realizations. The principal size design uses *N* = 400, 800, 1600, 3200, 6400, 12800 reference neurons and six conditions: matched low viability (0.4, 0.4), E-dominant loss (0.5, 0.8), I-dominant loss (0.8, 0.5), intact populations, a near-fold point (0.56, 0.9), and a near-Hopf point (0.56, 0.56). The first five sizes have 20 independent realizations per condition and the largest has ten. Reference E/I counts are integer-rounded at the specified density ratio and then thinned. Each comparison uses the achieved viable counts and fractions, including the finite reference-density ratio, rather than the nominal values.

Temporal and between-realization uncertainty are assessed separately. Supplementary Methods specify the observation windows, integration refinements, trait cutoffs and sampling designs that isolate these error sources.

#### 4.3. A cutoff changes the limiting model

The clipped-trait experiment shows decreasing errors with size, but also a persistent inhibitory-rate bias in the E-dominant-loss condition (Supplementary Fig. S2). Clipping a Cauchy variable to a finite interval creates endpoint atoms; it does not produce the unbounded Cauchy law used by the exact reduction. An independent stationary quadrature agrees with the unbounded MPR solution to approximately 10^−12^ and predicts a clipped-at-80 inhibitory-rate shift of approximately −0.01038. Increasing the number of neurons alone cannot remove this bias.

Unbounded-trait validation uses a projective/Riccati integrator whose single-neuron substep is exact for a frozen recurrent current and can count multiple spikes. It comprises 180 size runs over three conditions and 40 timestep controls; recurrent coupling still introduces splitting error. An independent cross-integrator check reduces complex-phase trace difference from about 4.54 × 10^−4^ to 1.29 × 10^−5^ under step refinement. For E-dominant loss, the primitive-observable root-mean-square error (RMSE) decreases from 0.04889 at *N* = 400 to 0.009951 at *N* = 12800. Intact-state values decrease from 0.009600 to 0.004552. Realization variability and occasional basin-dependent outliers remain visible. These comparisons separate finite-population error from the bias introduced by clipping the trait distribution.

#### 4.4. What a finite-size slope measures

The composite primitive error includes E/I rates and real and imaginary parts of E/I phase moments. Derived *R*_*a*_ and *M*_*a*_ are not counted again as independent channels. A pooled descriptive size fit in the clipped-trait common-size design has slope −0.3113 (95% Wald interval −0.3595 to −0.2632; seed-cluster bootstrap interval −0.3633 to −0.2617), with substantial observable and condition dependence. This number combines quenched sampling, finite temporal exposure, numerical error and nonlinear response. It describes this finite-range experiment rather than a universal *N* ^−1/2^ law.

Indeed, for unbounded positive Cauchy excitability tails, a single-cell stationary rate grows as 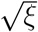, giving a logarithmically divergent quenched second moment. Bounded phase moments and spike-rate averages therefore need not have the same sampling asymptotics. Moreover, E and I counts covary in the primary design; separate E/I error exponents are unidentifiable after saturated condition effects are included. Crossed numerical-count controls and a projected count-noise predictor are reported as diagnostics. The latter performs worse than a minimum-count predictor on held-out stable conditions, consistent with its omission of quenched, cutoff and basin effects. Finite-count shot-noise theory provides a complementary approximation [6, 7], not a substitute for the microscopic convergence controls.

### 5. Finite-count escape: initialization, censoring and inference

The full 12-dimensional neural/synaptic state is simulated with class-specific Poisson counts of intensity *n*_*b*_*r*_*b*_. Each presynaptic class count simultaneously drives both postsynaptic filters with increment 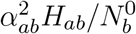. This mesoscopic count approximation is distinct from the finite deterministic QIF network. The midpoint neural step, homogeneous synaptic propagator and end-of-step counts are included. No reported count trajectory required a positivity-floor intervention. Initialization uses an approximate quasi-stationary distribution (QSD).

At *λ*_*I*_ = 0.9, the fold is 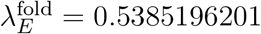; the chosen fold distances set 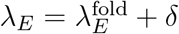 before integer rounding. The achieved distances enter geometry and noise calculations. Equilibrium-escape inference is restricted to the 30 cells in Supplementary Table S4 whose high-rate equilibrium is asymptotically stable. At *δ* = 0.08 that equilibrium exists but is unstable and therefore does not define a metastable well. The smaller-distance, larger-size design independently probes the joint small-noise and near-fold scaling, rather than extending the broad-grid fit without changing its asymptotic regime.

**Supplementary Fig. S2.**
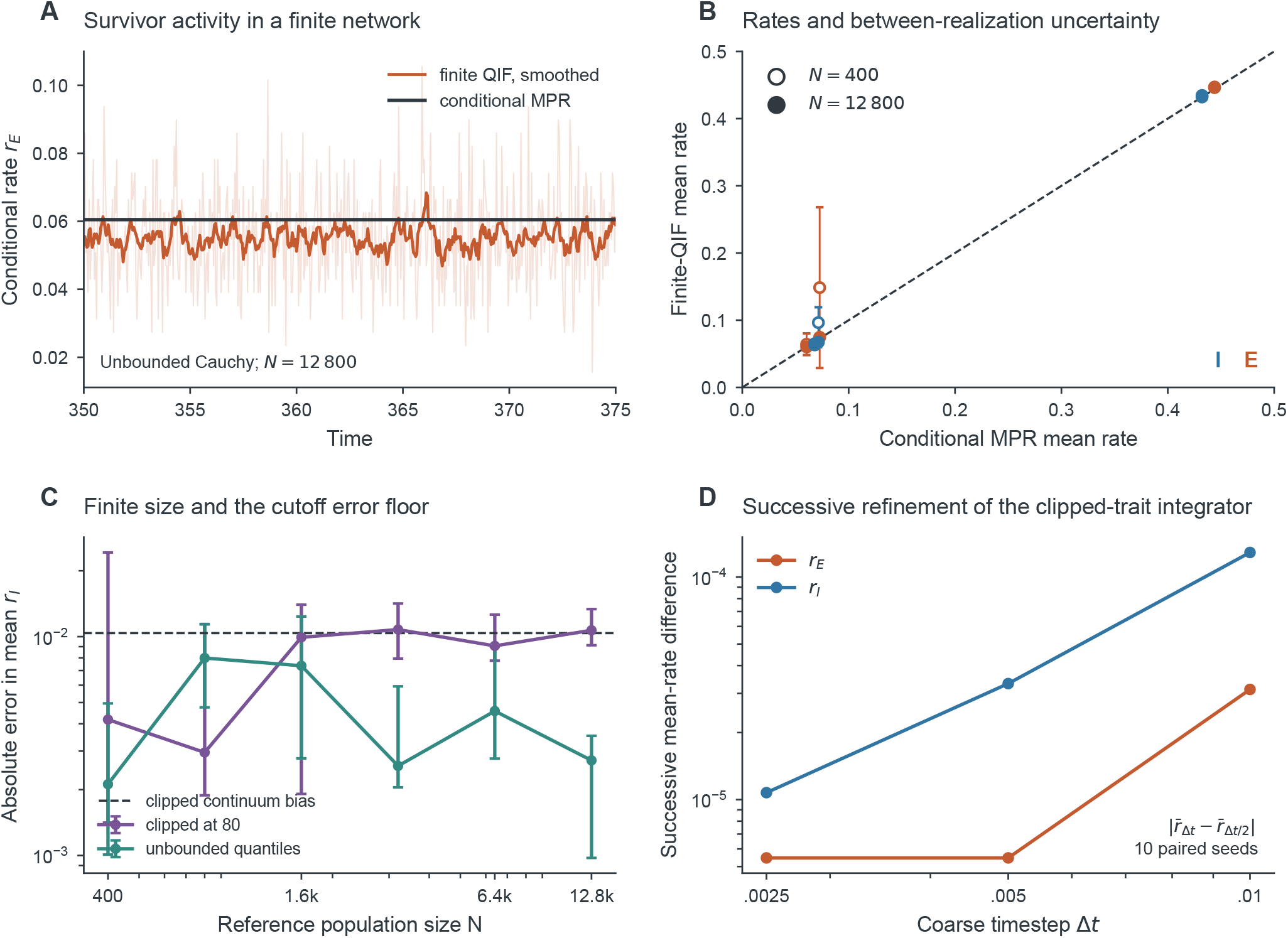
Finite neuronal populations, quenched variability and numerical bias. **(A)** One seeded unbounded-Cauchy QIF realization at *N* = 12800 under E-dominant loss, compared with the achieved-fraction conditional theory. Pale and dark curves show raw 0.05-unit samples and a display-only 0.5-unit average. **(B)** Mean E/I rates at *N* = 400 (open) and 12800 (filled), with nominal 95% between-realization Student-*t* intervals over ten unbounded realizations. **(C)** Median absolute inhibitory-rate errors and interquartile ranges for clipped-trait and unbounded-trait designs. The dashed level is the independently calculated clipped-law bias. **(D)** Measured successive-timestep differences in the clipped-trait integrator over ten paired realizations. All runs have 100 units of burn-in and at least 400 observation units. The clipped and unbounded distributions define distinct continuum comparisons.

**Supplementary Table S4.** Count-process designs and exact observation accounting. Each successful cell has 50 paths. Exposure is summed in model-time units; initialization particle-exposure is separate.

| Design | Cells | Paths | Events | Analyzed exposure |
| --- | --- | --- | --- | --- |
| Admissible grid | 30 | 1500 | 1226 | 1387811.1 |
| Timestep controls | 8 | 400 | 228 | 858716.1 |
| Joint local/weak-noise limit | 6 | 300 | 257 | 81186.9 |
| QSD particle/burn controls | 3 | 150 | 136 | 143478.3 |
| Total | 47 | 2350 | 1847 | 2471192.4 |

Grid: $N = 500, 1000, 2000, 4000, 8000, 16000$ crossed with $\delta = 0.01, 0.02, 0.03, 0.04, 0.06$ . Timestep controls: $(N, \delta) = (1000, 0.02), (4000, 0.04), (8000, 0.06), (16000, 0.04)$ at 0.01 and 0.0025, compared with grid step 0.005. Joint limit: $\delta = 0.005$ at $N = 16000, 32000, 64000$ and $\delta = 0.0025$ at $N = 32000, 64000, 128000$ . Actual simulated observation exposure is 2523219.7; QSD initialization particle-exposure is 1072243.2. Rejected nonstationary cells are not included.

Across 44 principal and refinement conditions and three additional reservoir controls, 2350 paths produce 1847 analyzed events and 2,471,192.4 units of analyzed exposure. Actual simulated first-passage exposure is 2,523,219.7, with a further 1,072,243.2 particle-time units for initialization. Zero-event conditions contribute their full exposure and one-sided bounds.

#### Augmented-state QSD initialization and causal events

Two independent Fleming–Viot reservoirs per cell each contain 64 particles. Interacting-particle approximation of quasi-stationary laws provides the methodological precedent [8]; diffusion convergence theorems do not automatically cover this augmented count-and-detector process. Every particle carries all 12 state variables, the last 21 E-rate samples (spacing 0.1), and its persistence counter. When killed it clones a uniformly chosen survivor’s entire augmented state. The operational escape event occurs after 20 consecutive moving-average samples fall below the computed saddle E-rate; the event is timed at confirmation and never backdated. This is a causal stopping rule, not a computed global separatrix. Reservoir burn-in is at least max(100, 20*t*_rel_), where *t*_rel_ is the leading modal relaxation time of the stable high-rate equilibrium, yielding durations of 100–394 units in the 44 principal and refinement cells. Initialization kills and particle exposure are saved, rather than counted as observation escapes. Late moment drift and between-reservoir distances quantify approximate equilibration. Fifty observation paths sample empirical particles with replacement, 25 per reservoir, and use distinct random streams. They are independent conditional on the estimated reservoirs, whose particles remain correlated through cloning. Three controls double reservoir size to 128 and burn-in to at least max(200, 40*t*_rel_), at (*N, δ*) = (1000, 0.01), (8000, 0.04), (16000, 0.005).

#### Exposure rule and inferential scope

Every observation path is simulated to escape or 4800 units. The analysis horizon is the first of 300, 1200, 4800 containing at least 30 events, or 4800 if none qualifies. All 50 records are included and clipped at that common horizon. This is data-adaptive censoring; fixed-horizon sensitivity at all three values is therefore also reported. Kaplan–Meier product limits [9], Greenwood log-log intervals and Nelson–Aalen hazards use explicit risk sets. Censored Weibull shape diagnostics require at least five events; the nominal shape-one test rejects in 1 of 27 eligible grid cells, which is not proof of exponential survival. Let *D* be the analyzed event count and *T* the summed at-risk exposure. Event/exposure rates 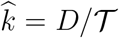 have profile intervals under an exponential working model. Three zero-event grid cells retain one-sided 95% upper bounds − log(0.05)*/ T*; at full exposure *T* = 240000 this is 1.24822 × 10^−5^. Exact finite-sample coverage under the adaptive horizon or full propagation of reservoir uncertainty is not claimed.

The 14-cell near-fold, weak-noise subset consists of main-grid and joint-limit cells with specified *δ* ≤ 0.02 and *N* ≥ 2000. The separate timescale-only selection retains main-grid cells whose estimated mean dwell time is at least ten local relaxation times, without imposing a fold-distance cutoff. Thus the two selections test different asymptotic requirements.

For condition *i*, 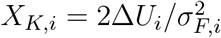 uses the local fold barrier and projected noise variance defined in the main article. Local Kramers scaling [10, 11] is assessed using log *k*_i_ = −*β*_0_ *−β*_1_ *X*_*K,i*_ and the censored exponential likelihood ∑_*i*_(*D*_*i*_ log *k*_*i*_ *− k*_*i*_ *T*_*i*_), including zero-event cells. The full 30-cell grid gives *β*_1_ = 0.5849 with profile interval [0.5657, 0.6045]; its stratified trajectory-bootstrap interval is [0.5625, 0.6089]. The six joint-limit cells give 1.0383 with profile interval [0.9298, 1.1499]. Adding the local prefactor as a fixed offset gives slopes 0.6090 and 1.0677, respectively. Bootstrap intervals condition on the empirical reservoirs. Near-unit local slope does not validate the prefactor, absolute rates, or the full process’s global large-deviation law. Supplementary Fig. S4 displays the survival, exposure, reservoir and integration diagnostics.

**Supplementary Fig. S3.**
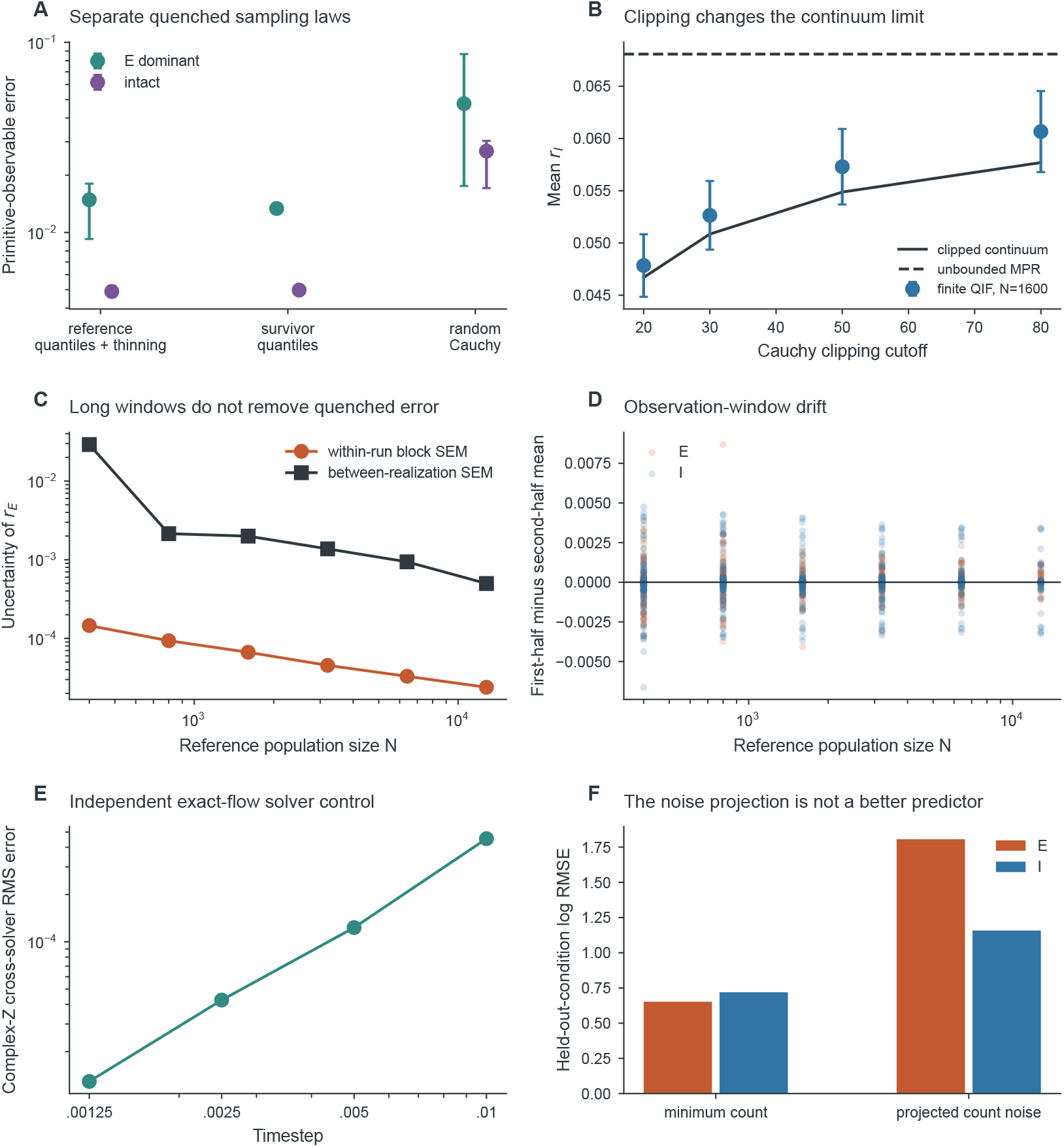
Microscopic error sources and independent controls. **(A)** Primitive mean-observable root-mean-square error (RMSE) for three trait-sampling laws; medians and interquartile ranges over ten realizations. **(B)** Clipped-law stationary continuum, unbounded MPR, and finite-network inhibitory rates with nominal 95% between-seed Student-*t* intervals. **(C)** Within-run block and between-realization standard errors of the mean (SEM) are separate uncertainties. **(D)** Half-window rate differences for all 660 clipped-trait size runs, without excluding drifting realizations. **(E)** Independent theta/Riccati phase-trace differences under refinement. **(F)** Held-out-condition log RMSE: the projected count-noise predictor is worse than minimum count in the three stable conditions tested. No universal finite-size convergence law is inferred from these finite-range results.

### 6. Finite-rate viability control

The matched control decreases from 1 to 0.34 and returns to 1 with 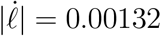. The rate-based transition detector gives a down/up separation of 0.268884. This externally imposed reversal is distinct from irreversible neuronal thinning. The frozen-state comparison uses the equilibrium and periodic branches in main-text Sec. 5.

### 7. Deterministic continuation and observation-design details

#### 7.1. Continuation coordinates and reference values

The arclength metric is Euclidean in the supplied coordinates; rates are log-transformed for equilibrium continuation, and the period is log-transformed for shooting. Residuals and step lengths accompany accepted points. Corrector failure, domain boundaries and maximum-step termination delimit the computed branches; rejected points are not replaced by grid interpolation.

**Supplementary Fig. S4.**
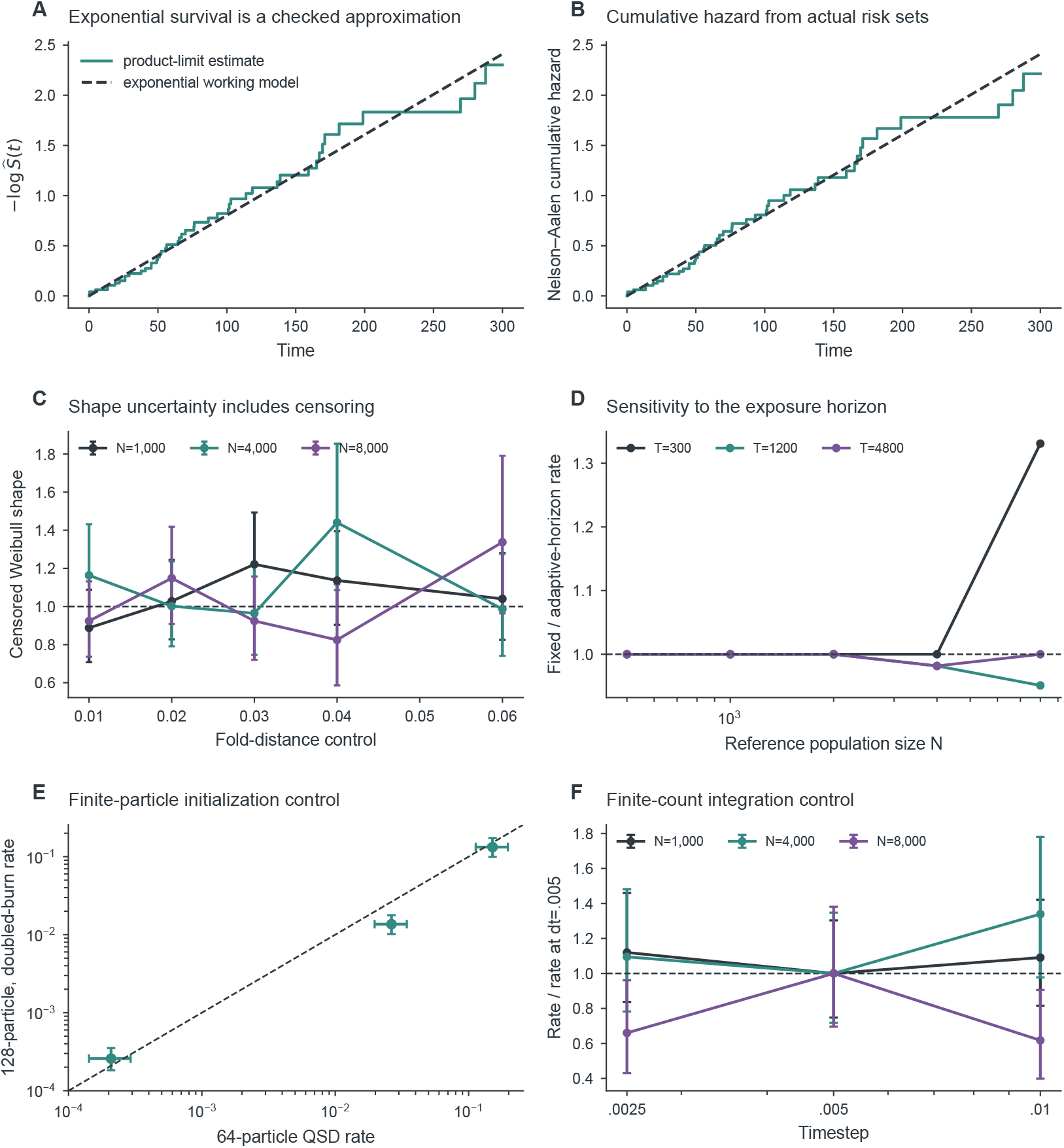
First-passage inference and sensitivity controls. **(A**,**B)** Negative log survival and Nelson–Aalen hazard for *N* = 4000, *δ* = 0.03, compared with the constant-hazard working model. **(C)** Censored Weibull shapes and profile intervals; one denotes exponential survival. **(D)** Fixed/adaptive horizon rate ratios for *δ* = 0.03, using matched paths; zero-rate cases remain in the source table. **(E)** Three 128-particle, doubled-burn controls against the standard 64-particle initialization, with conditional observation-rate profile intervals. **(F)** Rates and intervals normalized by the step-0.005 point estimate. Denominator uncertainty is not propagated in these display ratios; separate condition-level rates and intervals support inference. All panels retain the finite-reservoir and adaptive-exposure qualifications in the text.

Continued folds reproduce the equilibrium count at all 841 exploratory grid points. The high-state fold on *λ*_*I*_ = 0.9 occurs at *λ*_*E*_ = 0.538519620099, with *A*_*F*_ = −3.130248419 and *B*_*F*_ = 1.820226871 under the main-text nullvector convention. The matched, faster-I and faster-E paths cross the reference high-state Hopf boundary at *ℓ* = 0.555341165, 0.879677768 and 0.765173022. All three crossings are transverse and supercritical, with *l*_1_ = −0.484502, −0.00397793 and −0.837798, and frequencies 1.580699, 2.665385 and 1.381404 radians per model time. At *ℓ* = 1, the E-rate peak-to-trough amplitude is 0.666636, the period is 2.801836 and the largest nontrivial Floquet modulus is 0.192398. The 90 physical periodic branch points approach amplitude 6.812 × 10^−4^ near Hopf. At *ℓ* = 0.55, the E-rate transient amplitude decreases from 0.011469 over *t* ∈ [160, 260] to 6.782 × 10^−10^ over [1500, 1600], consistent with leading decay exponent −0.012429 (Supplementary Fig. S1).

**Supplementary Fig. S5.**
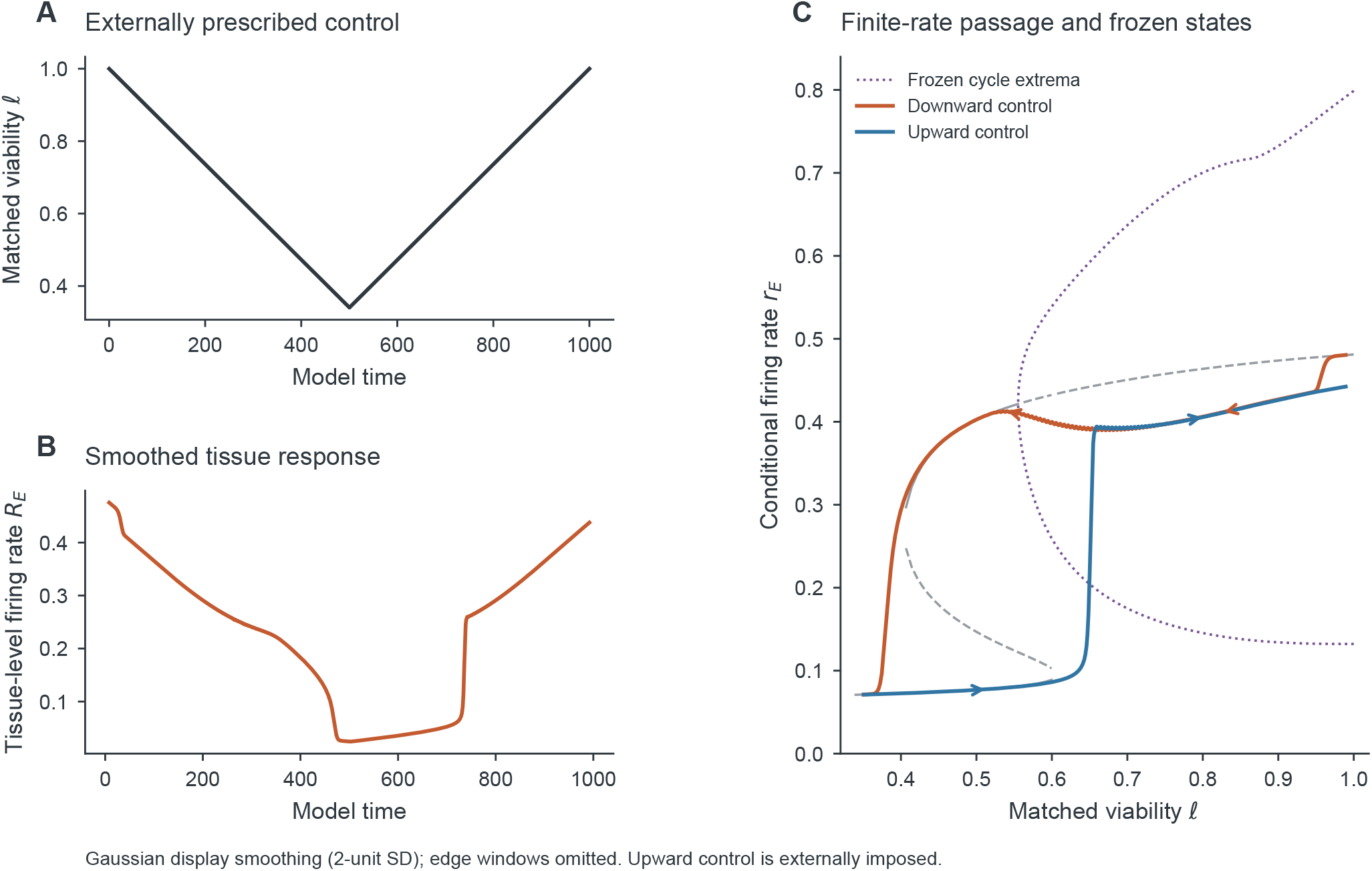
Finite-rate viability control combines frozen-state coexistence with dynamic lag. **(A)** Externally prescribed triangular matched-viability control. **(B)** Tissue-level E firing rate with Gaussian display smoothing (standard deviation two model-time units). **(C)** Conditional rate on the downward (rust) and upward (blue) paths, compared with stable (solid gray) and unstable (dashed gray) frozen equilibria and periodic-orbit extrema (dotted purple). Arrows indicate control direction. Eight-unit edge windows are omitted from smoothed curves; transition estimates use the rate-based detector, not the display smoothing. The upward limb is externally prescribed.

#### 7.2. Parameter-ensemble admission

Coupling magnitudes, heterogeneity widths, intrinsic centers, synaptic rates and the reference density ratio were independently perturbed by up to 10%. Of 157 proposals, 128 passed the admission test: an unstable oscillatory intact high-state equilibrium and a sustained intact trajectory after 180 units of burn-in and 180 observation units. E-rate amplitude had to exceed 0.05, half-window amplitude differences had to be below 5%, and the period coefficient of variation below 3%. Among 87 members with comparable regular Hopf crossings on all paths, faster I loss crossed first in 21 cases (24.14%, bootstrap 95% interval 16.09–33.33%), and faster E loss in 66 (75.86%, 66.67–83.91%). Thirty-eight members had verified path topology changes and three retained unresolved calculations. Marginal regular-path counts were 104, 99 and 99 for matched, faster-I and faster-E paths. This ensemble samples a dynamically selected parameter neighborhood, not a distribution over biological populations.

#### 7.3. Mechanism matches and observation scales

At (*λ*_*E*_, *λ*_*I*_) = (0.6, 0.9) with *ν*_*aI*_ = 0.5, reducing *λ*_*I*_ to 0.81, setting 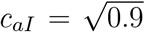, or withdrawing compensation to *ν*_*aI*_ = 0 gives the same effective inhibitory source factor. The conditional rates are (*r*_*E*_, *r*_*I*_) = (0.0615226, 0.0685963) and the leading eigenvalue is −0.606528 with corresponding transformed initial conditions. The tissue-level I rates are 0.0555630 for the viability change and 0.0617367 for the other two. The intrinsic-center match 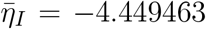 gives the same *r*_*E*_, but *r*_*I*_ = 0.0650761 and leading eigenvalue −0.596556.

The nine-parameter local design uses signed fractional reference coordinates, rate and response scales 0.1, eigenvalue scale one and identity working observation covariance. Stationary summaries plus a relaxation eigenvalue have numerical rank five. Complex responses at *ω* = 0.5, 1, 2, 4 raise the rank to eight, leaving the exact density–coupling null direction (1, −1, −1). For a unit external E-current perturbation, transfer functions use 401 logarithmically spaced frequencies 0.01 ≤ *ω* ≤ 30. The largest absolute difference among product-equivalent conditional responses is below 2.1 × 10^−17^. Full singular-value spectra and refinement controls appear in the observation-design methods.

### 8. First-passage initialization and observation protocol

For each realization record an event indicator *d*_*i*_ and observed time *t*_*i*_ = min(*T*_*i*_, *C*_*i*_). Under a constant-hazard exponential working model,

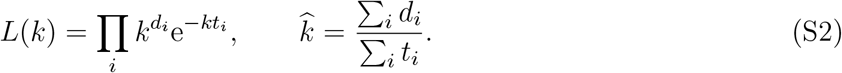

Right-censored non-events contribute exposure. For zero events, the one-sided 95% exponential-model bound is −log(.05)/ ∑_*i*_ *t*_*i*_. For positive event counts, the profile-likelihood interval solves 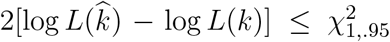. These working-model intervals depend on the constant-hazard assumption and the stopping rule.

At each distinct event time *t*_*j*_, let *d*_*j*_ be the number of events and 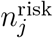 the number of trajectories still at risk immediately before that time. These are trajectory counts, not neuronal counts. The Kaplan–Meier survival and Nelson–Aalen cumulative hazard estimates are

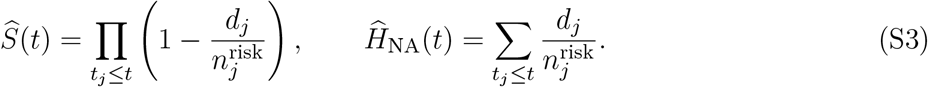

Tied censors remain in the risk set until that time and are not subtracted again at later events. Greenwood/log–log intervals and Weibull-shape diagnostics accompany the curves [9]. Nonexponentiality diagnoses the event process without by itself contradicting the deterministic fold calculation.

The main grid uses *λ*_*I*_ = 0.9, 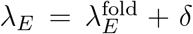, with *δ* ∈ {0.01, 0.02, 0.03, 0.04, 0.06} and *N* ∈ {500, 1000, 2000, 4000, 8000, 16000}. The high equilibrium at *δ* = 0.08 is unstable and is excluded from high-state escape experiments. Each valid cell has 50 first-passage paths, plus independent step, joint-limit and initialization controls.

Initialization approximates a quasi-stationary distribution with two independent Fleming–Viot reservoirs of 64 particles. Each particle carries the full 12-dimensional state, causal rate-averaging memory and crossing persistence counter. A particle is killed when its averaged E rate remains below the saddle E rate for 20 consecutive 0.1-unit samples; its complete augmented state is cloned from a surviving particle. Burn-in is at least 100 units and 20 modal relaxation times. First-passage paths have independent random streams conditional on the estimated reservoirs. The saddle-rate stopping threshold is not equated with the full multidimensional basin boundary.

Each path runs until a confirmed event or 4800 units. The analysis horizon is the first of 300, 1200 or 4800 units with at least 30 events, or 4800 if that target is not reached. All 50 paths contribute, including right-censored paths. Kaplan–Meier and Nelson–Aalen estimates, profile-likelihood intervals, Weibull shape diagnostics, fixed-horizon analyses and doubled reservoir/burn-in controls accompany the estimates. Intervals are nominal working-model intervals conditional on estimated reservoirs. Adaptive horizons and correlated reservoir particles limit an exact-coverage interpretation [9].

The quasi-stationary initializer includes causal detector memory in its state. Particle replacement is a finite-particle approximation to conditioning on no detected escape. Reservoir differences, burn-in extension and particle-number controls assess initialization separately from first-passage uncertainty. Adaptive analysis horizons are also checked against a common fixed horizon.

### 9. Higher-order bifurcation and wave reference values

**Supplementary Table S5.** Reference values at the original reported precision. Main-text coordinates, matrices and eigenvalues are rounded for display; the coefficient conventions and numerical estimates are unchanged.

| Quantity | Reported value |
| --- | --- |
| GH $(\lambda_E, \lambda_I)$ | (0.9382654414, 0.8468556406) |
| GH $\omega$ | 2.680980 |
| HH $(\lambda_E^H, \lambda_I^H)$ | (0.941627337027, 0.851027619454) |
| HH $(\omega_1, \omega_2)$ | (2.681500680, 4.323263742) |
| HH $\omega_2/\omega_1$ | 1.6122553 |
| $\Gamma$ | $\begin{pmatrix} -2.72266658 & 2.19469219 \\ 3.07045276 & -1.45034639 \end{pmatrix}$ |
| $\det \Gamma$ | -2.78988905 |
| $A^{\text{DH}}$ | $\begin{pmatrix} 0.000473166 & -0.109474682 \\ -0.339554885 & -0.016482303 \end{pmatrix}$ |
| $\beta^{\text{DH}}$ | $\begin{pmatrix} -0.122966171 & -0.138855053 \\ -0.060085386 & -0.003879428 \end{pmatrix}$ |
| $\det A^{\text{DH}}$ | -0.03718046 |
| Line-wave transverse pair | $-0.0809198335 \pm 3.5161974710i$ |
| Homogeneous sheet root, $k = 0$ | $0.080564 + 2.199097i$ |

At generalized Hopf, *ω* = 2.680980 and *l*_2_ = −0.00646343. The fifth-order homological residual is below 9 × 10^−16^; the augmented Jacobian and transverse spectral gap remain nonzero. Analytic cubic and quintic normal forms and a nonlinear coordinate shear test coefficient normalization. At *λ*_*I*_ = 0.84975, 111 periodic solutions connect unstable and stable cycles through a fold at *λ*_*E*_ = 0.9405981518, with E-rate amplitude 0.065928 and period 2.344331. Its distance from Hopf is 5.80277 × 10^−7^; the quintic prediction is 5.80511 × 10^−7^, a relative difference of 0.0405%. The augmented residual is 1.51 × 10^−14^, with 18 tolerance controls.

At double Hopf, shooting with initial modal amplitudes 0.02, 0.04 and 0.06 gives six secondary bifurcation points, with residuals below 3.6 × 10^−13^. At amplitude 0.04, the mode-1 point has (*λ*_*E*_, *λ*_*I*_) = (0.942054432, 0.851557071), period 2.343275027 and critical multipliers −0.760554 ± 0.649275i. The mode-2 point has (*λ*_*E*_, *λ*_*I*_) = (0.941740305, 0.851248383), period 1.453377317 and critical multipliers −0.728333 ± 0.685223i. Continued slices contain 118 distinct physical periodic solutions and verify the crossing signs. At amplitude 0.02, the predicted parameter displacements differ from shooting by 0.029% and 0.507%. The finite-amplitude constraint is a linear adjoint projection, not the nonlinear normal-form amplitude.

The torus at (*λ*_*E*_, *λ*_*I*_) = (0.942017140, 0.851555826) has frequencies Ω_1_ = 2.681119958 and Ω_2_ = 4.323472267. Odd angular grids with 9, 13 and 17 nodes reduce the oversampled invariance residual from 6.62 × 10^−8^ to 3.81 × 10^−14^. A full-system trajectory agrees with the surface evolution to 2.33 × 10^−9^ over 4,000 time units and gives 1,707 positive modal-phase section crossings. Real normal exponents 0.0003909296293 and −0.0004195546250 have oversampled eigenfunction residuals below 2.1 × 10^−12^ and remain separated from tangent planes. The tangent-quotiented variational cocycle gives the same signs.

For the line wavetrain, 50 corrected profiles span matched viability approximately 0.70085–0.94745. At *ℓ* = 0.82, the refined speed is *c*_w_ = 0.8718430885. The 96- and 128-point even-grid controls differ by 4.2 × 10^−7^; the 32-point corrector does not converge. Odd grids 65, 97, 129 and 193 support the stability calculation. At 193 points, the full-state residual is 2.47 × 10^−13^ and the translation eigenvalue is approximately 5.1 × 10^−13^. The leading nontrivial pair is −0.0809198335±3.5161974710i and its full-traversal Floquet modulus is 0.5581253. The translation-vector residual is 3.36 × 10^−5^, or 1.28 × 10^−8^ after Jacobian-infinity-norm scaling. Twelve corrected 97-point branch samples have no unstable nontrivial collocation eigenvalues. The complete finite-grid spectra concern the fixed periodic domain; wavelength and Bloch tests address additional perturbation classes.

Linear response about a stable stationary field is tested by applying its eigenmode to that same field and comparing the consistent tangent and nonlinear numerical maps. For initial amplitude *ϵ*, the finite-time nonlinear remainder is *O*(*ϵ*^2^) for a sufficiently smooth underlying flow.

### 10. Spatial discretization and numerical convergence

The plane kernel is periodized with nine image cells, retaining each image’s actual path length; minimum-distance wrapping would define a different model. The continuous omitted-tail bound is below 7.2 × 10^−7^. Spatial midpoint quadrature error is recorded separately: the largest pre-normalization zero-frequency (DC) defect decreases from 0.00587 at 64^2^ to 0.000737 at 128^2^. Discrete weights are normalized, and interpolation between adjacent delay shells preserves total mass and first delay moment. Full-state Heun integration evaluates delayed convolution at both stages. The Fourier ring-buffer implementation is checked against direct small-grid convolution. The simulation design uses 64^2^, 96^2^ and 128^2^ grids, 100 units of burn-in, 300 observation units, step 0.01 and shell width 0.04, with separate 0.005-step and 0.02/0.01-shell controls.

Nonoverlapping 10-, 20- and 50-unit block summaries distinguish estimands and expose drift. A diagnostic screen requires half-window mean change below 1%, standard-deviation change below 10% and frequency difference below 5%; passing it is not a statistical proof of stationarity.

Longer integration uses full-state checkpoints including emitted-source delay history. A restart initialized without that history is treated as a separate initial-value problem with an additional burn-in. A checkpoint test compares uninterrupted and checkpoint-resumed trajectories numerically. Table S6 compares the separate short-horizon refinements; Supplementary Fig. S6 relates these tests to the longer-window summaries and coherent-wave spectral refinement.

**Supplementary Table S6.** Matched finite-horizon spatial convergence at *t* = 8. Errors are actual successive-grid conditional-rate root-mean-square (RMS) differences; no finest-grid zero is plotted.

| Control | Successive comparison | Absolute RMS | Relative RMS |
| --- | --- | --- | --- |
| Time step | 0.01 → 0.005 | $3.240 \times 10^{-5}$ | $7.659 \times 10^{-5}$ |
| Time step | 0.005 → 0.0025 | $8.085 \times 10^{-6}$ | $1.912 \times 10^{-5}$ |
| Spatial grid | $64^2 \rightarrow 96^2$ | $8.377 \times 10^{-5}$ | $1.980 \times 10^{-4}$ |
| Spatial grid | $96^2 \rightarrow 128^2$ | $2.040 \times 10^{-5}$ | $4.823 \times 10^{-5}$ |
| Delay shells | 0.04 → 0.02 | $4.116 \times 10^{-5}$ | $9.731 \times 10^{-5}$ |
| Delay shells | 0.02 → 0.01 | $9.401 \times 10^{-6}$ | $2.223 \times 10^{-5}$ |

The coherent-wave problem uses a phase-conditioned Fourier boundary-value solve, not a fitted space–time slope. Its exact counterpropagating transport representation yields 20 states per collocation point for stability. Complete spectra at 65/97/129/193 odd grids, translation-mode residuals, finite-difference Jacobian checks and 12 continued wave states are reported. The 193-grid leading nontrivial exponent is −0.0809198335±3.5161974710i. Reported Floquet multipliers evolve over a full 2*π* traversal, two minimal periods of this wave-number-two solution. The primitive-period continuation and Bloch calculations below additionally probe wavelength dependence and sidebands; these finite spectral tests are not a proof of stability for every wavelength [12–14].

**Supplementary Fig. S6.**
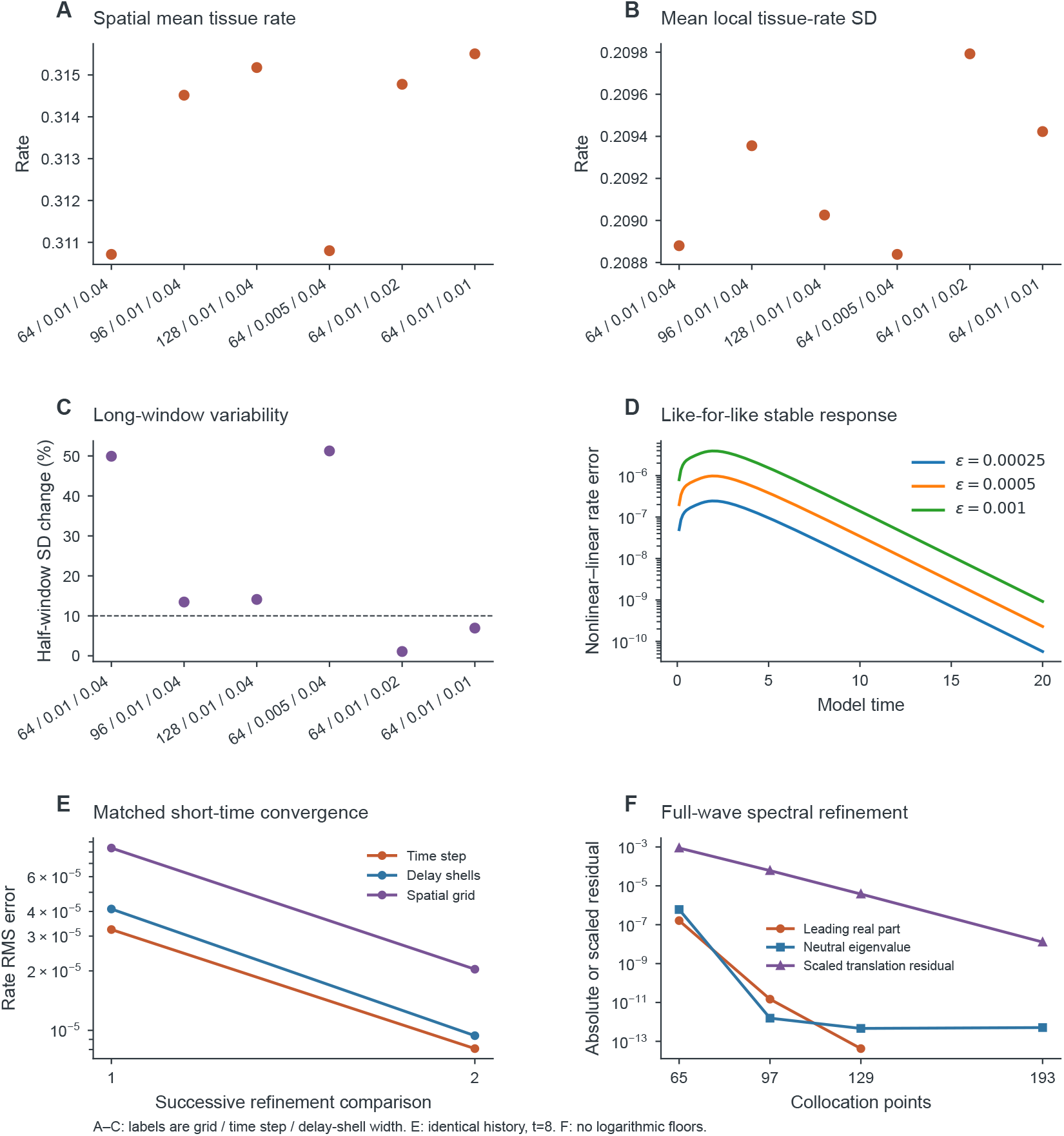
Spatial summaries and consistent linear comparisons. **(A**,**B)** Long-window spatial mean tissue-level E firing rate and spatial mean local tissue-level firing-rate standard deviation. Labels identify grid, timestep and delay-shell width. **(C)** Half-window change in global-rate standard deviation; the dashed 10% line belongs to a descriptive screen, not a stationarity test. **(D)** Nonlinear–linear errors about the same stable stationary ring and the derivative of the same full Heun/history map. The maximum difference has perturbation-amplitude order 2.00282. **(E)** Successive full-field errors at *t* = 8 from identical initial histories: step 0.01/0.005/0.0025, shell width 0.04/0.02/0.01 and grid 64/96/128. **(F)** Coherent-wave spectral refinement on odd collocation grids: leading real part relative to the 193-point value, translation eigenvalue modulus, and translation-vector residual scaled by the generator norm. Exact zero differences are omitted from logarithmic axes rather than floored.

### 11. Double-Hopf and invariant-torus numerical methods

#### 11.1. Double-Hopf normalization and independent controls

The critical pairs are ordered by 0 *< ω*_1_ *< ω*_2_. Set 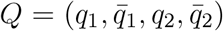 and 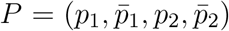, with ∥*q*_*j*_∥_2_ = 1 and *P* ^†^*Q* = I. The measured biorthogonality error is 8.8 × 10^−15^; the equilibrium residual is 4.0 × 10^−15^ and critical right/adjoint residuals are below 6.5 × 10^−14^. The remaining eight eigenvalues are strictly stable and at least 1.1273 from the imaginary axis. Total equilibrium-branch eigenvalue derivatives are evaluated by adjoint projection and checked against separately refined equilibria with parameter steps 2 × 10^−4^, 10^−4^, 2 × 10^−5^, 10^−5^ and 2 × 10^−6^. The common-time complex derivative matrix is

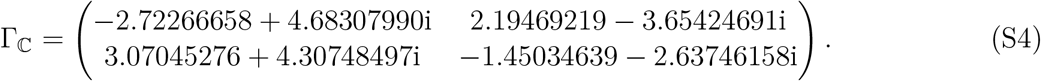

The real part of Eq. (S4) maps physical parameter changes to modal growth rates; the imaginary part supplies the first-order frequency detuning.

The main article gives the four-mode embedding, bordered homological equations and explicit Hessian–resolvent coefficient formulas. All four resonant cubic interactions are retained in the same original model time. The largest homological residual is 7.6 × 10^−16^. In the neuronal model C = 0, but the synthetic tests include nonzero cubic derivatives. All four coefficients agree between the two algorithms to 5.3 × 10^−16^. Coupled Stuart–Landau systems with known complex coefficients, a nonlinear coordinate shear, modal relabeling and positive amplitude rescalings test both algebra and conventions. With unit-norm real-rotation eigenvectors, the synthetic complex coefficients become twice the coefficients written directly in Cartesian complex amplitudes. The tests check this factor explicitly rather than comparing only signs.

For the amplitude classification, set 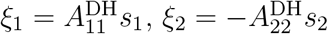 and *τ* = 2*t*. With 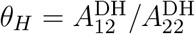 and 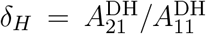, differentiation with respect to *τ* gives 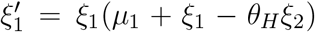 and 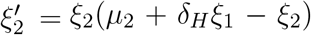. This defines the opposite-self-coupling case-IV label [15, Sec. 8.6, second edition] without depending on a diagram convention. Positive mixed amplitudes require *µ*_2_ *>* 0 and *µ*_2_*/δ*_*H*_ *< µ*_1_ *< θ*_*H*_*µ*_2_. The negative determinant of their radial derivative rules out an attracting mixed state and a radial Hopf bifurcation in this local cubic system. Eliminating the nonzero pure-mode radial coordinate gives secondary cubic coefficients −78.5781 on mode 1 and +2.25578 on mode 2. The first secondary bifurcation is supercritical relative to a cycle that already has a radial unstable direction; the second is subcritical relative to the stable mode-2 cycle. Both yield saddle mixed states locally.

#### 11.2. Full-state periodic branches and invariant torus

Secondary shooting uses unknowns (*y*_0_, log *T, λ*_*E*_, *λ*_*I*_). The 12 periodicity conditions are supplemented by 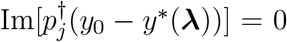 and 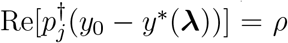. The adjoint *p*_*j*_ is fixed at the double-Hopf point, while the equilibrium and its parameter derivatives move with ***λ***. The last condition is log |Λ_*c*_| = 0 for the tracked, nontrivial complex multiplier. DOP853 tolerances are 2 × 10^−12^ and 2 × 10^−14^; the complete variational matrix and both parameter sensitivities are integrated. Periodicity derivatives are analytic, and the multiplier-condition row is centrally differenced. The amplitudes *ρ* = 0.02, 0.04, 0.06 give three points per mode. The largest augmented residual is 3.56 × 10^−13^ and the largest phase-neutral flow residual is 5.74 × 10^−12^. Two fixed-*λ*_*I*_ periodic slices through the *ρ* = 0.04 points use pseudo-arclength continuation, evaluate all multipliers and terminate at prescribed amplitude or parameter boundaries. Their 120 continuation samples contain 118 distinct physical solutions, because each bidirectional slice stores its initial point twice. Neither termination is claimed as a bifurcation.

The torus Fourier ansatz solves the rigid-angle invariance equation with two unknown frequencies and phase constraints ⟨*Y* − *Y*_ref_, ∂_*j*_*Y*_ref_⟩ = 0 for *j* = 1, 2. The brackets denote the uniform discrete angular inner product. The invariance derivative includes the full state Jacobian at every node, the Fourier differentiation matrices and the two frequency columns. The phase-reference tangents are normalized before the bordered Newton solve. The method follows the invariant-torus boundary-value formulation [16]; it is not a periodic-orbit shooting calculation. Starting from normal-form radii (0.035, 0.030), odd angular grids 9^2^, 13^2^, 17^2^ are solved to a residual below 3 × 10^−12^. Independent 19^2^, 27^2^, 35^2^ grids give invariance residuals 6.62 × 10^−8^, 3.83 × 10^−11^ and 3.81 × 10^−14^. The last frequency change is 2.75 × 10^−14^. On the finest verification grid the minimum pointwise singular value of (∂_1_*Y*, ∂_2_*Y*) is 0.01762, checking immersion rather than only an average tangent rank.

Full ordinary differential equation (ODE) integration uses DOP853 with relative/absolute tolerances 2 × 10^−11^/2 × 10^−13^ for 4,000 time units. State output spacing is 0.1; comparison with the Fourier surface is sampled every five time units. Positive crossings of 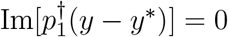 are detected directly by the integrator. The two mean projected modal radii are 0.0349834 and 0.0300290, close to their cubic predictions. Hann-window spectra are reported only as finite-window signatures; the resolved angular frequencies correspond to 0.4267135 and 0.6881020 cycles per model time.

Two independent normal-stability calculations supplement this trajectory comparison. First, a quotient variational cocycle projects out both instantaneous torus tangent directions after each five-time-unit integration block and QR-orthogonalizes two radial-like initial vectors. The base point is reset to the corrected torus at each block, but the projected perturbation vectors are not reset. The maximum base-flow reset error is 7.04 × 10^−12^. Integration to 20,000 time units gives full-window averages (3.7309, −4.0687) × 10^−4^; excluding the first 1,000 time units gives (3.5199, −3.9303) × 10^−4^. These finite-time averages have bounded angular modulation and are not presented as precise asymptotic Lyapunov exponents.

Second, the angular variational eigenfunction problem is solved on all three Fourier grids. Two real eigenfunction exponents converge to 0.0003909296293 and − 0.0004195546250, with changes below 6.2 × 10^−14^ from 13^2^ to 17^2^ nodes. At unit maximum-component normalization, their oversampled residuals are 1.56 × 10^−12^ and 2.06 × 10^−12^; boundary Fourier-energy fractions are below 5.7 × 10^−24^. Their minimum normal components are 0.421 and 0.327, respectively, after tangent projection. Thus the eigenfunctions are smooth resolved normal directions, not unresolved high-wave-number spectral artifacts or phase modes. An independent two-oscillator analytic torus tests the variational operator: its two known radial eigenfunctions have the specified normal exponents and both angular tangents are neutral. The full neuronal values agree closely with the cubic radial predictions 0.0003909233263 and −0.0004194322163. For a torus these are angular eigenfunction/variational calculations, not Floquet multipliers over a fictitious common period. Opposite signs establish the numerical saddle classification; no attracting torus is inferred.

### 12. Two-dimensional patterns and matched-defect methods

#### 12.1 Spatial initial conditions, history matching, and diagnostic thresholds

Writing *g*_0_ for the leading homogeneous growth rate and *g*_*k*_ for the largest searched growth rate at a nonzero admissible wave number, the three selection scores are *g*_0_ − *g*_*k*_, *g*_*k*_ −*g*_0_, and |*g*_*k*_ − *g*_0_|. The first two are maximized; the third is minimized among conditions with max(*g*_0_, *g*_*k*_) *>* 0. The reference condition is always retained. These are growth-rate *difference* scores, not independent maxima of *g*_0_ or *g*_*k*_.

The 2D solver uses the same image-periodized radial kernel, linearly interpolated delay shells, and full-state Heun map as the field convergence calculations. A duplicated circular Fourier-history buffer permits a strided view of past emissions without changing the operator. At 64^2^, an independent 250-step comparison with the direct indexed-buffer implementation has maximum full-state difference 1.71 × 10^−15^.

Let *X* = 2*πx/L, Y* = 2*πy/L*, and normalize a complex seed *F* so that 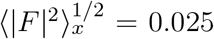. Rate perturbations are multiplicative 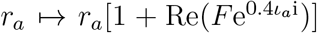 and voltage perturbations are additive 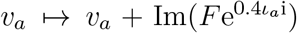, where *ι*_*E*_ = 0 and *ι*_*I*_ = 1 encode the neuronal class. The stripe uses *F* ∝ e^i*X*^ (or the explicitly specified diagonal e^i(*X*+*Y*)^). The radial seed uses 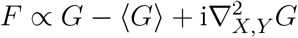, *G* = exp[2(cos *X* +cos *Y* −2)]. The periodic vortex-pair seed is 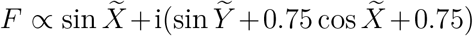, where 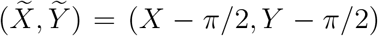. It has two isolated zeros of opposite winding on the torus. Broadband coefficients are independent fixed-seed complex Gaussian draws divided by |***k***|, for 0 *<* |***k***|^2^ ≤ 10 and component magnitudes at most three. Synaptic states begin at the common intact stationary value, and the initial emitted history is constant at the perturbed initial rate.

**Supplementary Fig. S7.**
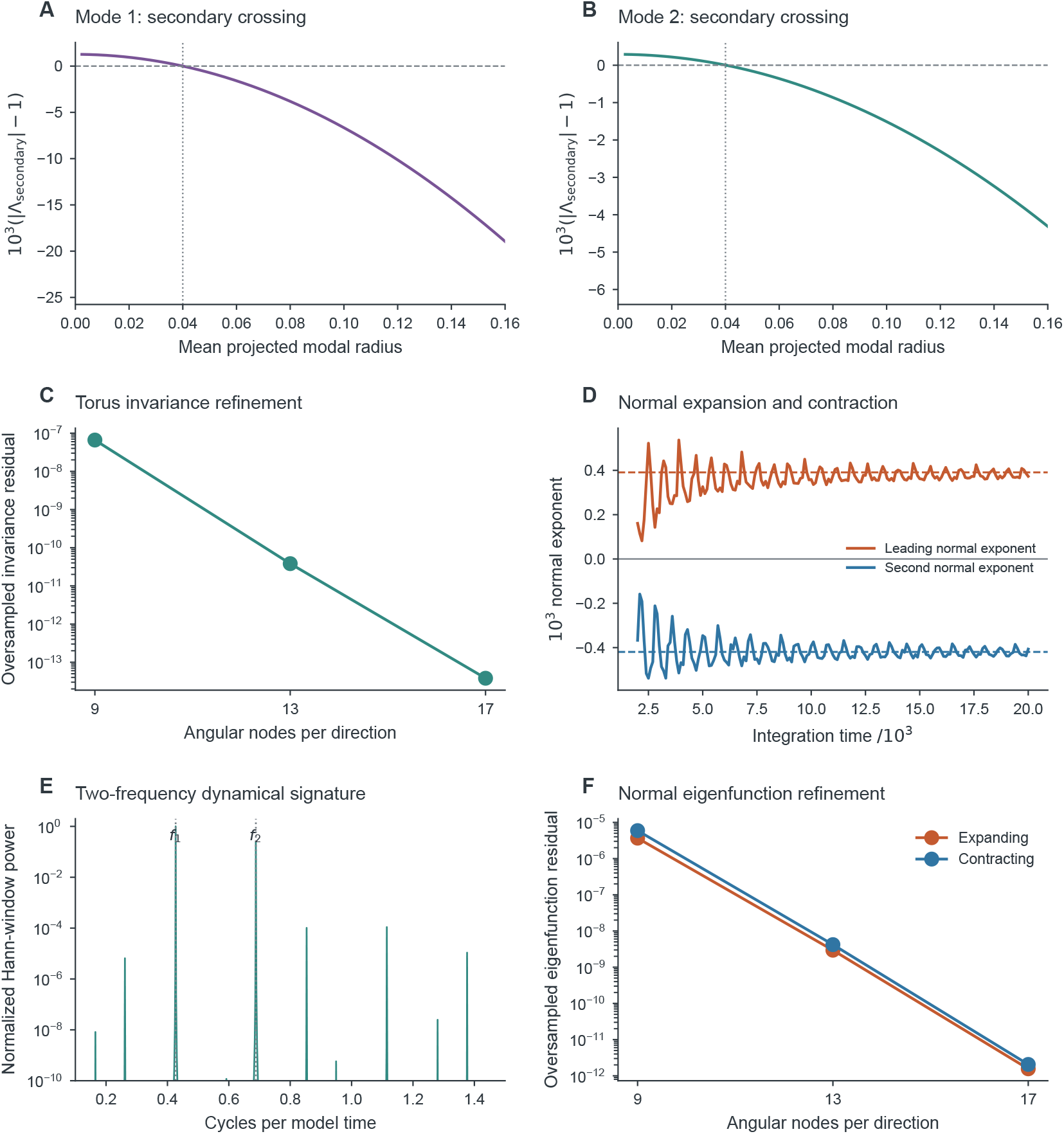
Independent dynamical and numerical controls for higher-order objects. **(A**,**B)** The tracked secondary complex multiplier modulus crosses one on each full-state periodic branch at fixed inhibitory viability; the phase multiplier is excluded. The horizontal coordinate is the mean projected modal radius, close to but not identical to the shooting initial amplitude constraint. **(C)** Independently oversampled torus-invariance residual under angular-grid refinement. **(D)** Running finite-time normal cocycle averages; dashed lines show the cubic radial predictions. These averages are distinct from the grid-refined normal eigenfunction exponents reported in the text. **(E)** A Hann-windowed rate spectrum of the 4,000-time-unit trajectory; marked frequencies come from the invariant-torus boundary-value solution, not spectral peak fitting. **(F)** Off-grid residuals of the expanding and contracting normal eigenfunctions decrease under independent angular-grid refinement. These controls distinguish torus invariance, dynamical signatures and normal stability.

Morphology is assessed over the final 100 time units. Dominant-band phase uses a fourth-order Butterworth bandpass spanning 0.75–1.25 times the dominant frequency, followed by the analytic signal; three cycles at both ends are excluded. Oscillatory RMS must exceed 0.5% of mean rate and the dominant spectral neighborhood must contain at least 50% of the power. A plane additionally requires at least 80% power in one nonzero spatial mode, directional alignment at least 0.90, phase-regression *R*^2^ ≥ 0.98, mode-amplitude coefficient of variation at most 0.30, and normalized median curvature at most 0.15. Radial alignment must exceed 0.90 on at least 60% of a fixed interior annulus. Failed conditions remain unresolved. Half-window mean and oscillatory-RMS changes below 5% and 10% are moment-stability checks, not proofs of an invariant distribution or attractor.

**Supplementary Table S7.** Two-dimensional simulation design. A sum denotes discarded initial transient plus observation time. Both delay-shell and timestep controls are independent, not simultaneous changes. Each depth–width design comprises 20 nonzero deficits and one shared zero-deficit control. All parameters, including initial-condition formulas, class ratios and source histories, are recorded for every condition.

| Experiment | Runs | Grid | Step | Shell | Time |
| --- | --- | --- | --- | --- | --- |
| Four-point, four-seed repertoire | 16 | $48^2$ | 0.01 | 0.04 | 100 + 200 |
| Planar/radial/rotating grid controls | 6 | $64^2, 96^2$ | 0.01 | 0.04 | 100 + 300 |
| Separate step and shell controls | 6 | $64^2$ | 0.005, 0.01 | 0.04, 0.02 | 100 + 300 |
| Matched reference intact/lesion | 2 | $64^2$ | 0.01 | 0.04 | 100 + 300 |
| Exact-history long continuations | 5 | $64^2$ | 0.01 | 0.04 | 400 more |
| Axial incident-crest depth/width design | 21 | $64^2$ | 0.01 | 0.04 | 200 |
| Reference diagonal depth/width design | 21 | $64^2$ | 0.01 | 0.04 | 200 |
| Finite-$k$-only two-seed follow-up | 2 | $64^2$ | 0.01 | 0.04 | 100 + 700 |

Core detection checks concentric loops with radii equal to two and three 48^2^-grid spacings, scaled to fixed physical size on refined grids. The minimum loop amplitude exceeds 5% of the spatial median analytic amplitude. An exactly zero grid vertex is handled by enclosing loops, not assigned an arbitrary phase. Same-charge tracks are matched with periodic distances and a fixed physical displacement cap. Persistence requires five dominant cycles. Unfiltered firing rate is sampled on a moving annulus of radius 0.4; at least five actual angular rotations must be measured, the first angular Fourier mode must contain at least 25% of modes 1–5, its angular regression must satisfy *R*^2^ *>* 0.98, and its speed must agree with the analytic-phase estimate within 25%. Edge-truncated lifetimes are marked as censored. A synthetic stationary phase vortex fails the rotation test, whereas a known rotating rate pattern passes it.

The numerical tests also cover exactly identical zero-deficit pairs, unchanged pre-intervention weighted histories, exact planar phase gradients and zero curvature, a smooth radial phase field with known gradient, opposite vortex charges, and known-speed crest arrival under three sampling intervals. Grid, timestep, and shell controls use identical smooth initial functions; long continuations preserve all 12 state components and the complete delayed history rather than reconstructing missing memory. Table S7 distinguishes the initial-condition repertoire, matched interventions and separate resolution tests.

#### 12.2. First-crest qualifications and complete spatial outcomes

The 79 trajectories in Supplementary Table S7 all complete without a numerical failure. Six additional short first-crest controls bring the total to 85 paths. Numerical completion is distinct from a successful morphology or crest classification; unresolved and transient outcomes are retained.

For first-crest assignment, rate maxima are quadratically interpolated in time from samples separated by 0.1. The reference crest at the most upstream sampled position is chosen after one quarter of the incident period. Peaks have at least one-half period separation and prominence at least 10% of the local peak-to-peak rate range. Subsequent reference maxima must lie within 0.3 periods of the expected arrival under the measured intact phase speed. Each lesioned maximum must lie within 0.4 periods of the corresponding reference. Arrival-time slopes are fitted separately in [−2, −*b*_*E*_], [−*b*_*E*_, *b*_*E*_], and [*b*_*E*_, 2]; all three must have the expected sign and *R*^2^ *>* 0.95. Each zone must contain at least four assigned sampling sites. A phase-equivalent arrival shift is reported as qualified only when all three regions pass. These finite-zone slopes describe crest deformation; they need not equal a single spatially invariant wave speed.

The first-traversal harmonic field uses the least-squares model

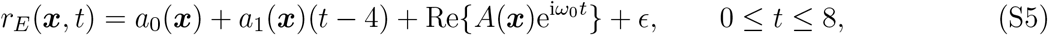

with the measured intact angular frequency *ω*_0_ and cycle frequency *f*_0_ = *ω*_0_/(2*π*). A site is valid when both intact and lesioned fits have *R*^2^ *>* 0.5 and amplitudes exceeding 10% of their respective spatial maxima. The circular mean phase contrast outside radius max(1.5, 1.5*b*_*E*_) is removed before calculating the RMS contrast within radius *b*_*E*_. At least 60% of the central area must be valid. Curvature additionally requires valid immediate neighbors and a nonvanishing phase gradient. The downstream peak-to-peak amplitude ratio uses max(*b*_*E*_, 0.8) ≤ *s* ≤ 2 and 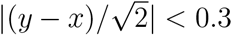 in the diagonal experiment. Tissue and conditional ratios are calculated independently, with the appropriate *λ*_*E*_ weighting. A crest with adequate support is classed as shifted when |Δ*t*|*f*_0_ ≥ 0.02; otherwise it is deformed when central phase-contrast RMS is at least 0.10 rad, and preserved below both thresholds. Missing support or an invalid arrival fit remains unresolved.

The separate axial depth–width experiment uses the transient 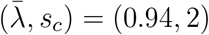 plane at *t* = 400. It gives one intact, ten preserved, nine deformed, and one shifted first-crest condition. Its later intact field becomes mixed, so those results support only the first traversal and do not establish long-time planar propagation. The reference diagonal experiment instead uses the plane that remains coherent through *t* = 1000. Its one unresolved condition is retained even though a candidate downstream maximum can be located: the complete three-zone qualification, not visibility of a maximum, determines validity.

At 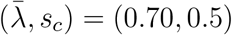, homogeneous linear growth is negative while a searched finite-wave-number mode grows. Two standardized axial stripe seeds, with wave numbers one and two, are followed to *t* = 800. Their final windows are respectively bulk-dominated (98.77% spatial zero-mode band power, *f* = 0.58081) and planar (93.04% dominant spatial-mode power, *f* = 0.30465), and both pass the half-window moment tolerances. These finite-window, seed-dependent outcomes do not establish basins of attraction, a subcritical bifurcation, or certified multistability.

##### Independent short first-crest refinement

An adaptive six-path control tests the representative diagonal interaction (*d*_*E*_, *b*_*E*_) = (0.48, 0.58) over only the first eight units after the intact *t* = 800 checkpoint. The 64^2^, *h* = 0.01 intact/lesion pair reproduces the first 81 rate/voltage snapshots and global observations of the depth–width experiment exactly. Separate pairs use 96^2^ with *h* = 0.01, and 64^2^ with *h* = 0.005; the shell width remains 0.04.

| Control | Downstream shift | Downstream–upstream | Intact speed |
| --- | --- | --- | --- |
| $64^2$, $h = 0.01$ | 0.155953 | 0.041174 | 1.212423 |
| $96^2$, $h = 0.01$ | 0.156633 | 0.041430 | 1.212110 |
| $64^2$, $h = 0.005$ | 0.155923 | 0.041176 | 1.212389 |

The spatial control spectrally prolongs the full state and the physical emitted-source fields, then transforms the latter using the finer fast Fourier transform (FFT) normalization. State and history round-trip errors are below 2.23 × 10^−15^ and 1.34 × 10^−15^; conditional/tissue means are preserved to 1.12 × 10^−16^, and all imported physical sources remain positive. The half-step control linearly interpolates chronological pre-intervention emissions, excluding the circular buffer’s obsolete next-write slot and using the known intact source at *t* = 800 as age zero. The coarse temporal second-difference divided by eight has maximum 8.98 × 10^−4^ and RMS 1.15 × 10^−4^ in physical-source units; these are interpolation-curvature indicators, not measured fine-history error bounds. All crest fits remain qualified. The downstream shift changes by 0.44% under spatial refinement and 0.02% under timestep/history refinement. The grids also sample the averaging zones differently. These imported-history, short-window sensitivities neither establish a convergence order nor replace independently resolved long histories or two-dimensional stability analysis. The downstream–upstream difference removes the contemporaneous upstream phase-equivalent arrival shift; neither quantity is a proof of causal information-transmission delay.

#### 12.3. Conditional and tissue-level activity in a heterogeneous line field

The observation identity is local. A viability deficit can preserve structured conditional activity while attenuating tissue-level firing at the same position and time. Supplementary Fig. S8 displays this distinction in a one-dimensional delayed field with class-specific localized loss. The paired rate maps share their coordinates and color limits, so the weaker tissue-level signal follows from multiplication by viable mass rather than a change in the plotting scale.

**Supplementary Fig. S8.**
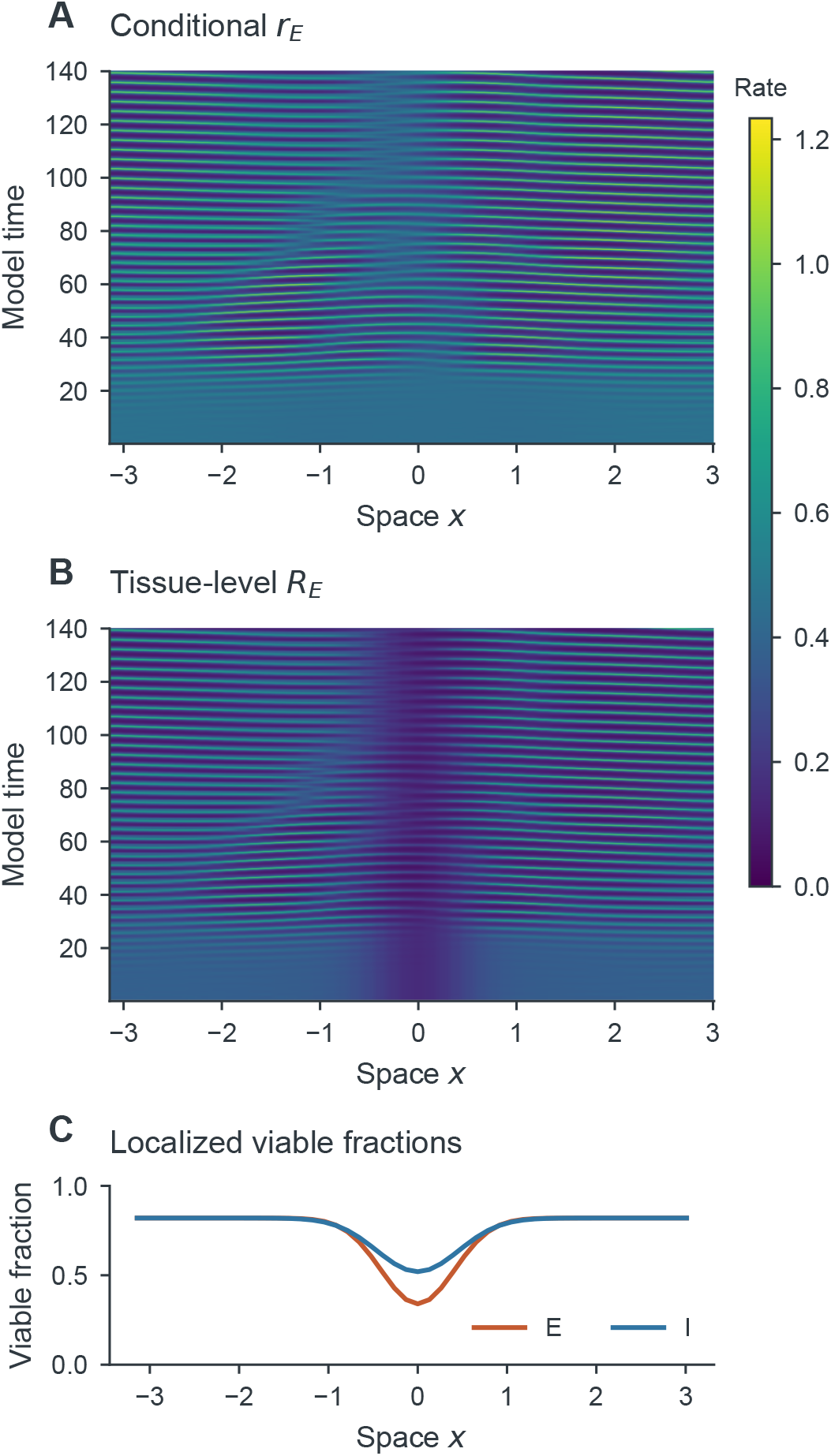
Conditional activity remains structured through a localized viability deficit. **(A)** Conditional E firing rate *r*_*E*_(*x, t*). **(B)** Tissue-level firing rate *R*_*E*_(*x, t*) = *λ*_*E*_(*x*)*r*_*E*_(*x, t*) from the same trajectory, with the same color scale. **(C)** E/I viable-fraction profiles. Conditional activity remains spatially structured through the deficit, while the local viable fraction attenuates its tissue-level firing rate.

This paired observation holds the conditional trajectory fixed when comparing normalizations. A comparison with an intact field must additionally account for the change in recurrent input caused by the deficit itself.

### 13. Paired local observations and sheet refinements

The 128^2^ trajectory in Supplementary Fig. S9 compares conditional and tissue-level fields at identical times and color limits. At the lesion center the observation-window mean conditional E rate is 0.406890, whereas its tissue-level firing rate is 0.138343. The corresponding mean local phase-moment magnitudes are 0.380949 and 0.129523.

Mean tissue-level firing rate and mean local tissue-level firing-rate standard deviation change by 0.210% and −0.157% between the 96^2^ and 128^2^ simulations. The selected mode frequency changes by 0.211% to 0.303124 cycles per model time. These are finite-window summaries. At 128^2^, the half-window global-rate mean and standard deviation differ by 1.19% and 14.1%. Other controls show larger global-amplitude changes and changes in dominant propagation direction. Longer 64^2^ continuation retains amplitude drift. The evolving field has local summaries that vary less under refinement than its global amplitude and propagation direction; these results do not identify a stationary two-dimensional traveling wave.

Local harmonic phase gradients define phase velocities and wavefront curvature where amplitude and gradient thresholds are satisfied. A lesion phase residual is measured relative to a fitted planar carrier outside the lesion, not as a causal delay relative to an intact-sheet experiment. Phase-based regions follow the selected propagation direction. Matched intact-field comparisons are reported in main-text Sec. 8.5.

Short-horizon controls compare full fields at *t* = 8, separating discretization from long-time state selection. Halving the timestep twice gives rate-error order 2.00244; halving shell width twice gives order 2.13040. Spatial 64^2^/96^2^/128^2^ comparisons use Fourier interpolation to a common physical grid. These tests assess discretization, while long-window diagnostics assess the stability of summary statistics.

### 14. Spatial repertoire and matched-field measurements

#### 14.1. Planar, radial and rotating activity

We classify planar, radial and rotating activity using separate spatial and temporal criteria. Byrne and colleagues demonstrated planar, radial, and rotating activity in a next-generation field with gap junctions and a balanced wizard-hat kernel[17]. Their two-dimensional brain-wave equation is a long-wavelength approximation; the E/I field studied here instead retains four positive radial exponential propagation kernels and their distributed delays, with no electrical coupling. Pattern classes in the two models therefore need not have the same parameter dependence.

We screen viable fractions 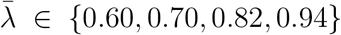, matched across the homogeneous E/I populations, and a common conduction-speed multiplier *s*_*c*_ ∈ {0.5, 1, 2}.

Growth-rate comparisons use the admissible square-domain wave numbers 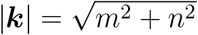, along-side the continuous dispersion curves. The reference point (0.82, 1) and extrema of the homogeneous-growth, finite-wave-number-growth, and competitive-growth scores select (0.94, 0.5), (0.94, 2), and (0.94, 1). Four standardized initial conditions—band-limited broadband, an axial stripe, a smooth radial seed, and a periodic vortex–antivortex pair—give 16 conditions at 48^2^, with 100 transient units and 200 observation units. The perturbations have the same complex root-mean-square amplitude, 0.025.

**Supplementary Fig. S9.**
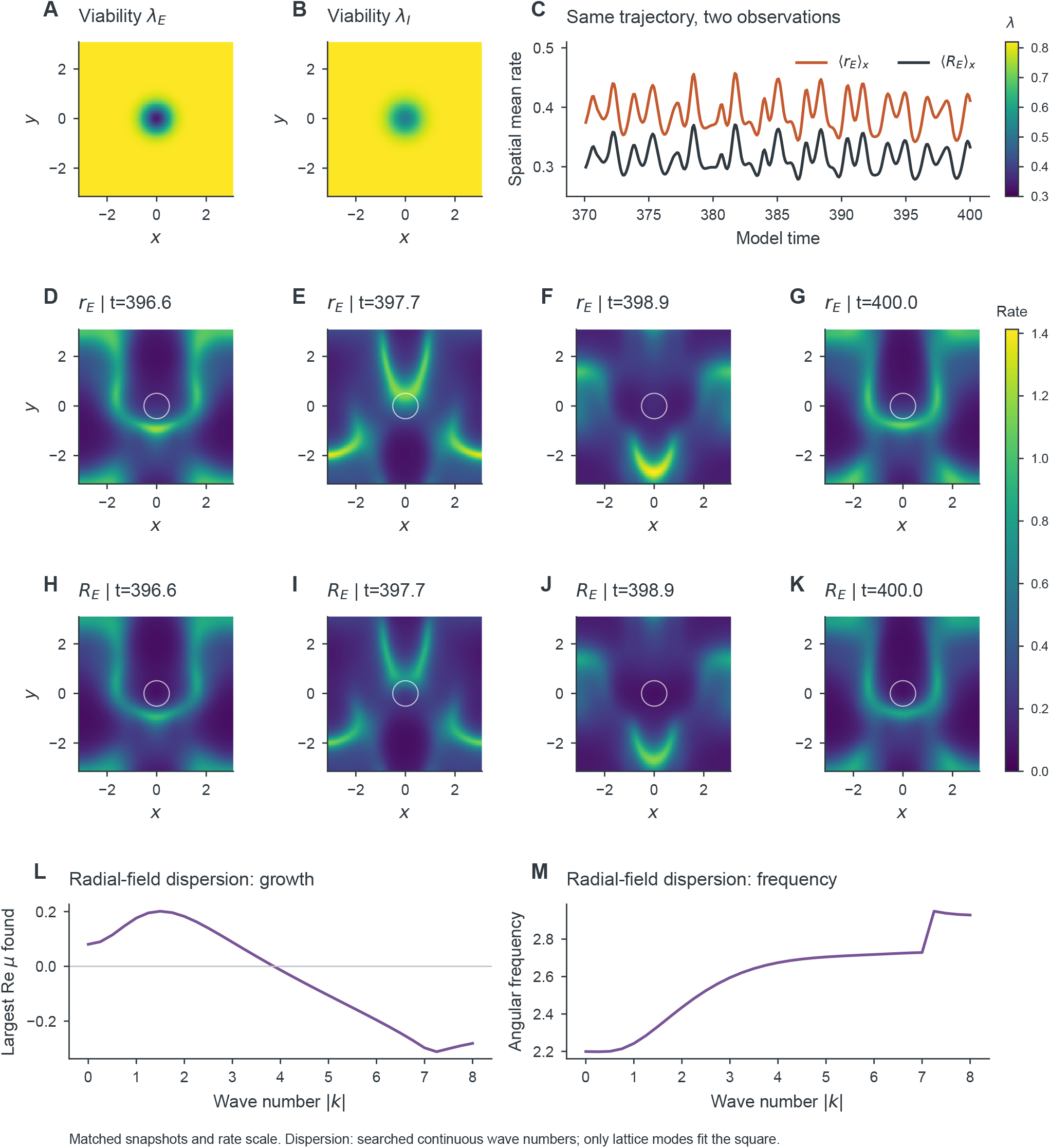
Survivor activity persists through a viability deficit while its tissue-level contribution is attenuated. **(A**,**B)** E/I viable-fraction maps. **(C)** Spatial means of local conditional E rate and tissue-level E firing rate from the same 128^2^ trajectory on a common axis. **(D–G)** Four conditional-rate snapshots selected by a fixed rule in the late window. **(H–K)** Corresponding tissue-level rates *R*_*E*_ = *λ*_*E*_*r*_*E*_ at identical times and a common rate scale; white contours mark *λ*_*E*_ = 0.6. The radial distributed-delay field uses 100 burn-in and 300 observation units. Its evolving patterns are distinct from the coherent wavetrain in main-text Fig. 8. **(L**,**M)** Largest growth rate found and its angular frequency in the continuous wave-number search about the homogeneous equilibrium. Only lattice wave numbers fit the square; the search is not exhaustive over characteristic roots.

Oscillation phase is obtained from the dominant temporal band of the conditional firing-rate fluctuation, rather than identifying it with arg *Z*_*a*_. With the convention 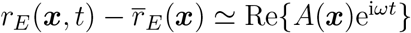 and *ω >* 0, the phase is *ϕ*(***x***) = arg *A*(***x***) and

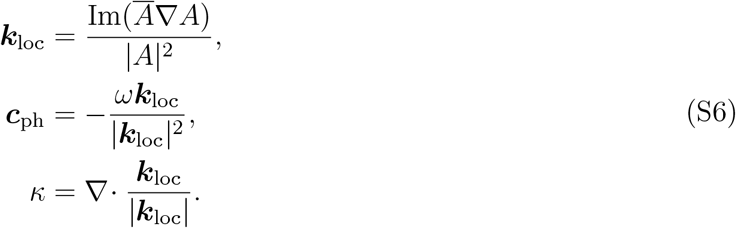

Here *ω* is angular frequency, ***c***_ph_ is a local phase velocity, and *κ* is wavefront curvature. These quantities are masked where amplitude or phase gradient is too small. A phase core requires integer winding

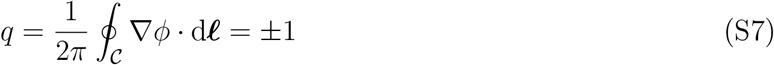

on two reliable surrounding loops. A vanishing amplitude at the center is compatible with, rather than evidence against, a genuine core. Opposite-charge tracks are followed on the periodic domain. A rotating-activity classification additionally requires at least five cycles of persistence and rotation of the first angular mode of the *unfiltered* firing rate around the moving core, consistent with the analytic-phase rotation. Thus a time-independent phase singularity alone does not establish rotating activity.

The windowed classification yields seven rotating-core conditions, two planar conditions, one radial condition, and six mixed or unresolved conditions (Supplementary Fig. S10). The homogeneous-growth-favored conditions are almost spatially uniform, but their temporal complexity fails the narrowband criterion. They are not relabeled as simple bulk oscillations. The distinction between finite-window morphology and persistence is important: the (0.94, 2) axial and radial seeds lose their original classification by *t* = 800. The radial seed also develops qualified rotating cores at 96^2^, so a resolution-independent radial attractor is not established.

In contrast, rotating activity from the reference vortex-pair seed is present at 64^2^ and 96^2^, with separate timestep and delay-shell refinements. At 96^2^, six tracks exceed the five-cycle threshold, with lifetimes up to 57.2 units (17.39 dominant cycles). Their unfiltered angular speeds have magnitudes 1.938–1.968, angular regressions explain more than 99.95% of the phase variance, and the first angular mode contains approximately 88–90% of the resolved annular angular power. The qualified total topological charge remains zero. Core creation, motion, and disappearance preclude identifying this activity with one rigidly rotating spiral solution. Rotating activity remains detectable in the 700–800 window of the exact continuation, including tracks lasting more than 15 cycles. The result is persistent rotating activity in the sampled field, not a spectral-stability proof for a two-dimensional spiral branch.

#### 14.2. Local loss can increase whole-sheet activity

To measure the effect of localized loss, the lesioned and intact reference fields start from exactly the same full neuronal and synaptic state and the same emitted-source history. Only *λ*_*a*_(***x***) changes at the intervention. Signals emitted before the intervention retain their original source weighting while they remain in flight. This prevents a lesion-specific equilibrium or a rescaled past history from becoming an uncontrolled second intervention.

A diagonal seed with wave vector (1, 1) at 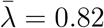 produces an intact plane that remains coherent through *t* = 800: the dominant analytic mode contains 99.991% of the band power and its phase speed is 1.21251. The corresponding lesion uses (*d*_*E*_, *d*_*I*_) = (0.48, 0.30) and (*b*_*E*_, *b*_*I*_) = (0.58, 0.65). Over *t* ∈ [400, 800], the spatially averaged conditional E rate changes from 0.344406 to 0.373541, while its tissue-level counterpart changes from 0.282413 to 0.301449. The lesion therefore increases both the spatial mean conditional E rate and the whole-sheet tissue-level E firing rate in this matched comparison. Locally, *R*_*a*_ = *λ*_*a*_*r*_*a*_ still attenuates the tissue-level firing rate at fixed conditional state. Globally, the state itself changes because the viable recurrent source has changed. Lower local viability thus need not imply lower whole-sheet mean firing.

**Supplementary Fig. S10.**
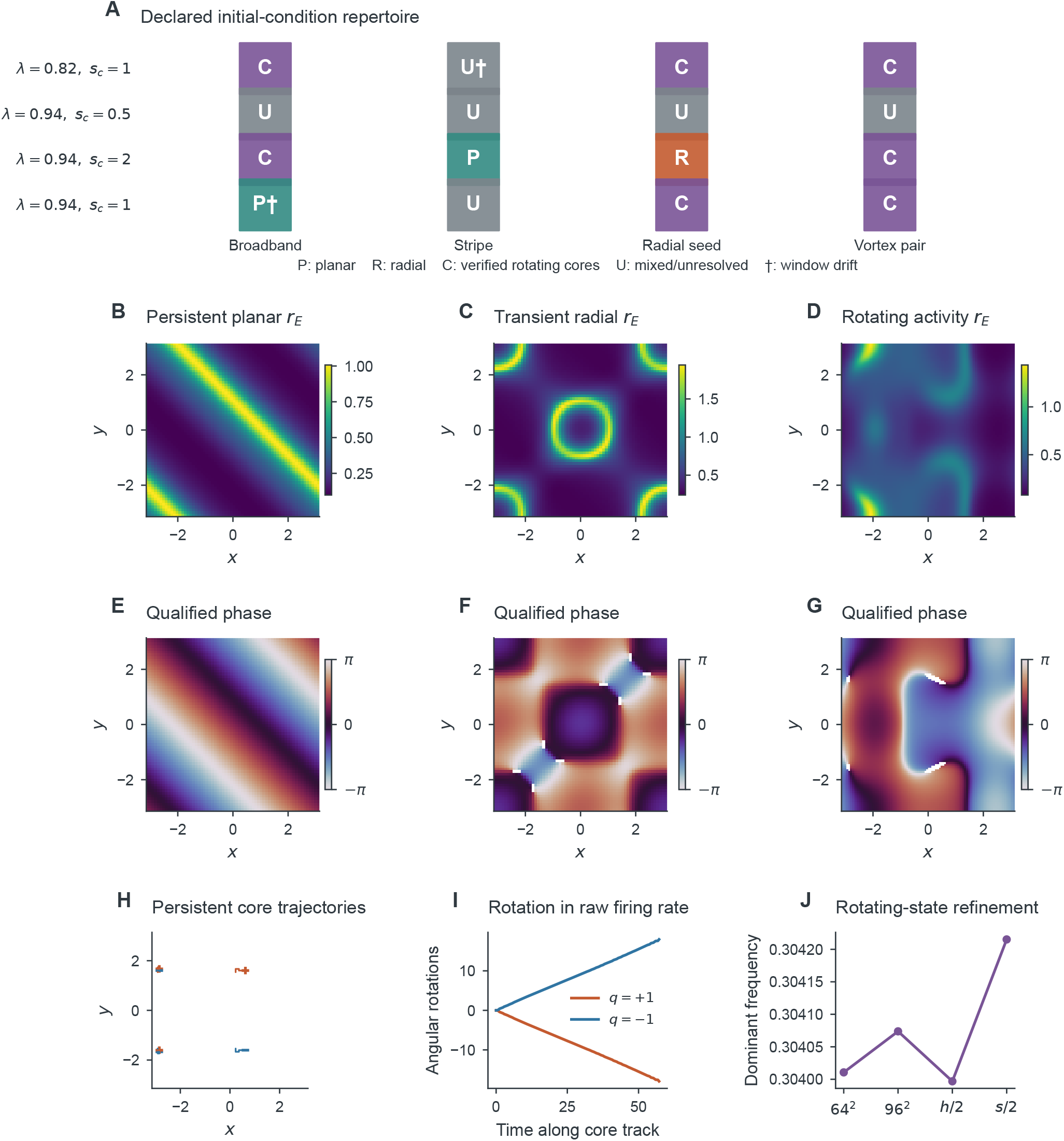
Persistent rotation is supported in selected conditions, whereas radial morphology can be transient. **(A)** All 16 parameter/initial-condition outcomes in the observation window; a dagger marks a failed half-window moment-stability criterion. P, planar; R, radial; C, verified rotating activity; U, mixed or unresolved. **(B–G)** Conditional-rate and simultaneous band-phase maps for the persistent reference diagonal plane at *t* = 780, the transient radial condition at *t* = 380, and the refined reference rotating condition at *t* = 380, respectively. Rate color limits are independent across these different conditions. Phase is masked below 10% of the frame’s maximum analytic amplitude. The radial map is not evidence of a radial attractor. **(H)** Persistent opposite-charge core tracks on the torus; apparent breaks at the boundary are periodic wrapping. **(I)** Angular rotations measured in the unfiltered firing rate along two opposite-charge tracks. **(J)** Dominant frequency under separate grid, timestep, and delay-shell refinements of the rotating condition. Long-time morphology and short-time discretization error are assessed separately.

The distinction extends beyond firing. The time mean of the magnitude of the spatially averaged conditional I phase moment, 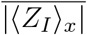, changes from 0.488517 to 0.413865, whereas 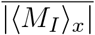 changes from 0.400584 to 0.335066. Main-text Fig. 7 separates simultaneous field snapshots from differences in time-averaged local observables. Spatial means of local conditional quantities remain distinct from whole-sheet survivor-normalized observables. The lesioned field passes the stated half-window mean and root-mean-square (RMS) tolerances in this late window, but its mixed spatial morphology is not classified as a stationary traveling wave.

### 15. Deficit depth–width results and crest measurements

A persistent intact plane provides a matched reference for measuring how deficit depth and width affect crest timing, amplitude and phase geometry. At *t* = 800, the full intact state and emitted history are shared among 21 conditions. The radial deficit has 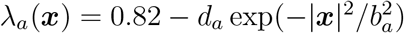, with *d*_*E*_ ∈ {0, 0.16, 0.32, 0.48, 0.64}, *b*_*E*_ ∈ {0.30, 0.45, 0.58, 0.80, 1.10}, *d*_*I*_ = 0.625*d*_*E*_, and *b*_*I*_ = (0.65/0.58)*b*_*E*_; one zero-deficit field suffices for all widths. The intact field also meets the planar criteria in the final 100-unit window of the further observation, ending at *t* = 1000.

The phase velocity points along (1, 1), so arrival is measured on the physical coordinate 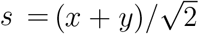 along *y* = *x*, rather than along an oblique Cartesian cut. A common incident crest is followed during the first eight units after the intervention. Let *t*_*d*_(*s*) and *t*_0_(*s*) denote its lesioned and intact arrival times. The two timing estimands are defined in main-text Eq. 88. The downstream phase-equivalent shift Δ*t*_+_ includes any upstream shift caused by the simultaneous intervention; it is not itself additional transit time across the deficit. Both quantities concern an identified oscillatory crest, not a causal information-transmission delay.

Of the 20 nonzero deficits, seven preserve the crest within the specified phase and timing tolerances, twelve exceed the 0.02-cycle downstream-shift threshold, and one is unresolved (Supplementary Fig. S11). For the unresolved (*d*_*E*_, *b*_*E*_) = (0.64, 0.80) condition, the within-deficit arrival fit fails the linearity criterion, *R*^2^ = 0.9486 *<* 0.95; it is not called propagation failure. Qualified downstream shifts range from 0.0151 to 0.5403 time units. The downstream tissue peak-to-peak amplitude ratio ranges from 0.9891 to 0.5666 across all nonzero deficits, while conditional amplitude and local phase geometry provide separate descriptions of the surviving population. The zero-deficit comparisons have exactly zero state/history differences, zero arrival shift, and unit amplitude ratios.

For (*d*_*E*_, *b*_*E*_) = (0.48, 0.58), the downstream and upstream shifts are 0.155953 and 0.114780, giving Δ*t*_cross_ = 0.041174. Conditional and tissue downstream amplitude ratios are 0.8962 and 0.8856, and the supported central phase-contrast RMS is 0.1197 rad after removing the common far-field offset. Separate short-window controls at 96^2^ and half timestep change the downstream shift by 0.44% and 0.02%. The refinement tests use the same intact signal history with spatial or temporal interpolation. They measure short-window sensitivity rather than the convergence order of a long-time attractor. Within this bounded depth–width set, the resolved effects are arrival shifts, tissue-amplitude attenuation and local phase deformation; no condition meets the criteria for pinning or loss of the incident crest. The unresolved condition remains distinct from propagation failure.

**Supplementary Fig. S11.**
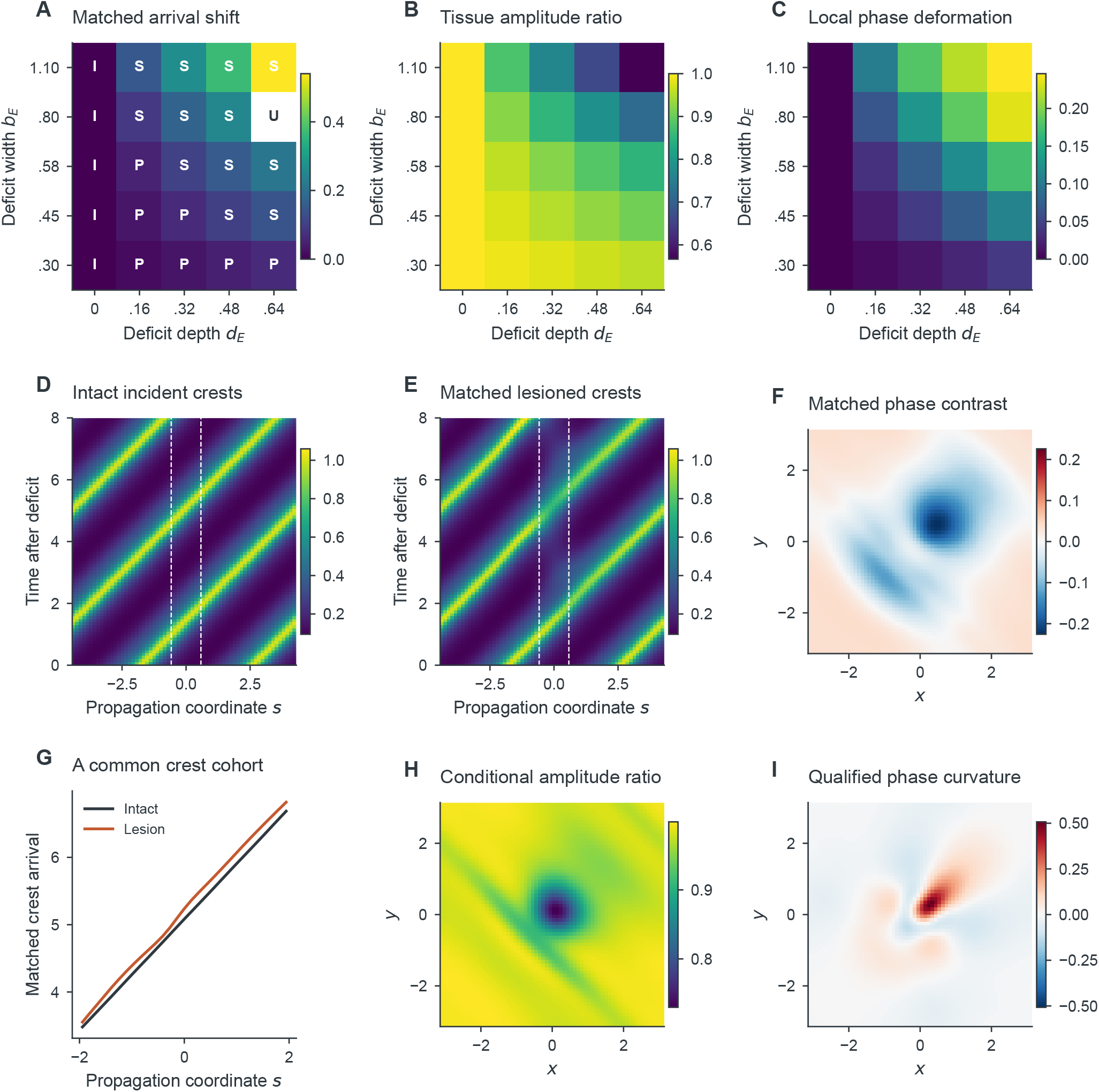
Viability deficits shift crest timing and attenuate tissue amplitude without qualified propagation failure in the tested range. **(A)** Downstream phase-equivalent arrival shift over the 21-condition design. I, intact; P, preserved crest; S, shifted crest; U, unresolved. The unresolved arrival estimate is masked, and the identical zero-deficit column is repeated only for display. **(B)** Downstream tissue peak-to-peak amplitude relative to the intact field over the same first eight units. **(C)** RMS central phase contrast after removal of a common far-field phase offset; only sites meeting the amplitude and harmonic-fit criteria enter this statistic. **(D**,**E)** Intact and lesioned conditional-rate cuts along *y* = *x* at (*d*_*E*_, *b*_*E*_) = (0.48, 0.58), plotted against physical distance 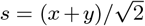, with common rate color limits; dashed lines mark *s* = ±*b*_*E*_. **(F)** Qualified local phase contrast in radians. **(G)** Matched phase-equivalent crest arrivals; their downstream difference must not be confused with the additional crossing shift in main-text Eq. 88. **(H)** Conditional harmonic-amplitude ratio and **(I)** qualified phase curvature over the first traversal. Signed maps use symmetric color limits containing all qualified values. Phase and curvature are descriptors of the oscillatory rate field, not direct measures of information transmission.

### 16. Coherent-wave wavelength and Bloch calculations

The traveling wave is corrected on its primitive spatial period (wavelength) *P* = *π*. Here *λ* = *λ*_*E*_ = *λ*_*I*_ = *ℓ* is the matched viable fraction, as in the main-text Bloch calculation and figure. The relative half-period translation defect of the 193-point doubled-cell state is 1.57 × 10^−8^, and its odd-harmonic power fraction is 6.17 × 10^−17^. Even Fourier coefficients supply a primitive-cell initial guess, followed by nonlinear correction; no uncorrected restriction is used as a wave solution. Five profiles are connected to the *λ* = 0.82 state using viability increments no larger than 0.01, and corrected at 129 and 193 points.

The wavelength continuation uses 65 nodal points, an integral phase condition and a weighted Euclidean arclength in the four neuronal profiles, wave speed and period. Each profile coordinate has weight 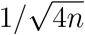; speed and period have unit weights. The initial period increment is 0.03, the maximum arclength step is 0.04, and failed correctors trigger step halving. The bounded search uses *P* ∈ [*π/*2, 2*π*] and at most 80 steps in either direction, producing 123 accepted points with positive rates and rate range exceeding 0.01. The largest boundary-value residual is 1.24 × 10^−11^. Nine wavelengths spanning the branch are corrected at 97 and 129 points; the maximum successive speed change is 3.39 × 10^−11^ and the maximum relative neuronal-profile change is 3.46 × 10^−6^. The bounds and step budget delimit the reported family; branch exhaustion and stability of the whole wavelength family are not inferred.

The full linearization retains 20 dynamical components and uses Toeplitz Fourier-Galerkin multiplication. Coefficient grids resolve every retained mode difference, without circular wrap-around convolution. Bloch derivatives act on every component, with comoving velocities 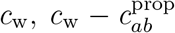 and 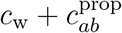 as appropriate. All eigenvalues of every retained generator are computed by dense eigendecomposition, rather than a single spectral shift. Table S8 summarizes the spectral sampling and convergence tests; Supplementary Fig. S12 compares spectral, translation-mode and wave-speed convergence.

**Supplementary Table S8.** Bloch design and independent numerical controls. All entries use matched E/I viability; no time integration or stochastic sampling is involved.

| Quantity | Design or result |
| --- | --- |
| Viabilities | 0.705, 0.76, 0.82, 0.88, 0.94 |
| Primitive wavelength | $P = \pi$ |
| 49-mode Bloch mesh | 19 Floquet numbers plus 18 interval midpoints per viability |
| 65-mode controls | $qP/\pi = 0, .005, .01, .02, .04, .0625, .125, .5, 1$ |
| 97-mode controls | $qP/\pi = 0, .005, .01, .02, .04, .5, 1$ |
| Complete Bloch spectra | $185 + 45 + 35 = 265$; dimensions 980, 1300, 1940 |
| Largest abscissa change, 49 to 97 modes | $7.78 \times 10^{-9}$ |
| Largest abscissa change, 65 to 97 modes | $3.51 \times 10^{-11}$ |
| 97-mode translation eigenvalue modulus | At most $3.65 \times 10^{-13}$ |
| Physical-cell parity-block matrix defect | $1.23 \times 10^{-15}$ |
| Physical-cell spectral set distance | $2.14 \times 10^{-11}$ |

The initial scaled Floquet mesh is 0, .02, .04, .0625, .125, .1875, …, 1; both it and every interval midpoint are evaluated at 49 modes. The smallest nonzero control *qP/π* = .005 resolves the translation-derived long-wave branch. The maximum positive-*q* spectral abscissa is negative (−8.25 × 10^−7^). At *q* = 0, only the eigenvalue closest to zero is removed for the transverse-abscissa statistic; its interpretation is checked against the projected profile derivative. No shift or positive floor is added to spectral errors for logarithmic plotting.

The physical cell-folding test compares a doubled-cell truncation with modes −48, …, 48 against primitive periodic modes −24, …, 24 and primitive antiperiodic modes −24, …, 23. This matched cutoff gives exactly 97=49+48 modes per component. After even/odd permutation the off-diagonal blocks vanish to roundoff, and the full doubled-cell spectrum equals the union of the two primitive spectra to the stated numerical tolerance. An independent nodal collocation calculation at 49, 65 and 97 points, with the same all-component Bloch derivative, converges to the same rightmost eigenvalues at *qP/π* = 0, .5, 1 for *λ* = .82.

Table S9 gives the adjoint phase coefficients at each viable fraction, separating comoving drift from diffusive relaxation.

**Supplementary Table S9.** Phase-diffusion coefficients from the adjoint bordered solve at 97 Fourier modes. The coefficient *a*_1_ is measured in the comoving frame.

| $\lambda$ | $c_w$ | $a_1$ | $D_{ph}$ |
| --- | --- | --- | --- |
| 0.705 | 0.8608396682 | 0.5718366724 | 0.0329848437 |
| 0.760 | 0.8574190582 | 0.6190217308 | 0.0938429834 |
| 0.820 | 0.8718430885 | 0.6805188408 | 0.1304126652 |
| 0.880 | 0.9012035651 | 0.7439236290 | 0.1481280171 |
| 0.940 | 0.9409923979 | 0.8018852873 | 0.1560899324 |

For the translation-derived branch, we fit Re *µ*(*q*) with an even quartic and Im *µ*(*q*) with an odd cubic. Both fits use *q* = 0, .005, .01, .02, .04; restriction to *q* ≤ .02 changes the diffusion coefficient by at most 2.50 × 10^−8^. The independent adjoint calculation solves the phase-normalized bordered system for the derivative of the translation eigenvector. Its relative linear-system residual is at most 1.74 × 10^−15^, and its diffusion coefficient agrees with the spectral fit within 2.32 × 10^−8^. The imaginary parts of the nominally real adjoint coefficients are below 2.02 × 10^−13^. Symmetric wavelength corrections at *λ* = .82 with period steps .002, .001, .0005 give *P* d*c*_w_/d*P* differing from the adjoint phase coefficient by 2.16 × 10^−8^, 5.41 × 10^−9^ and 1.35 × 10^−9^, respectively, consistent with the second-order difference formula. Synthetic tests additionally verify the all-component Bloch shift, *q* ↔ − *q* conjugacy, exact transport transfer functions and the projected Jacobian against finite differences of the independent nonlinear transport implementation. The difference between these two Jacobian estimates is 9.91 × 10^−11^.

**Supplementary Fig. S12.**
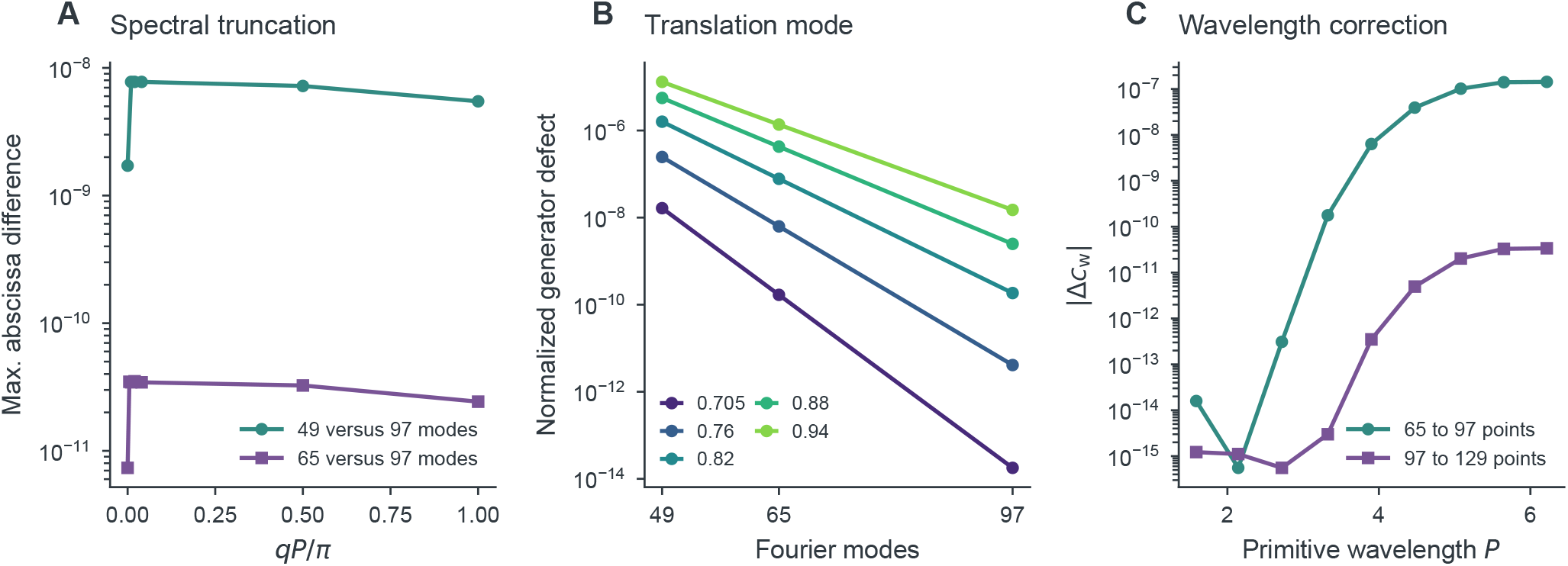
Coherent-wave numerical convergence. **(A)** Maximum rightmost transverse-abscissa difference across the five viabilities, comparing 49 or 65 Fourier modes with 97 modes at common Floquet numbers. At *q* = 0 the translation eigenvalue is excluded. **(B)** Norm of the generator applied to the projected wave derivative, normalized by the derivative norm and the generator infinity norm. **(C)** Successive wave-speed changes when selected wavelengths are corrected at 97 and 129 nodal points. Only strictly positive numerical differences are displayed on logarithmic axes; no artificial floor is introduced.

## Notes

### Competing Interest Statement

The authors have declared no competing interest.

## References

[1] Hugh R. Wilson and Jack D. Cowan. “Excitatory and Inhibitory Interactions in Localized Populations of Model Neurons”. In: Biophysical Journal 12.1 (1972), pp. 1–24. doi: 10.1016/s0006-3495(72)86068-5. URL: https://doi.org/10.1016/s0006-3495(72)86068-5.

[2] Edward Ott and Thomas M. Antonsen. “Low dimensional behavior of large systems of globally coupled oscillators”. In: Chaos: An Interdisciplinary Journal of Nonlinear Science 18.3 (2008), p. 037113. doi: 10.1063/1.2930766. URL: https://doi.org/10.1063/1.2930766.

[3] Ernest Montbrió, Diego Pazó, and Alex Roxin. “Macroscopic Description for Networks of Spiking Neurons”. In: Physical Review X 5.2 (2015), p. 021028. doi: 10.1103/physrevx.5.021028. URL: https://doi.org/10.1103/physrevx.5.021028.

[4] Christian Bick et al. “Understanding the dynamics of biological and neural oscillator networks through exact mean-field reductions: a review”. In: The Journal of Mathematical Neuroscience 10.1 (2020), p. 9. doi: 10.1186/s13408-020-00086-9. URL: https://doi.org/10.1186/s13408-020-00086-9.

[5] Stephen Coombes. “Next generation neural population models”. In: Frontiers in Applied Mathematics and Statistics 9 (2023), p. 1128224. doi: 10.3389/fams.2023.1128224. URL: https://doi.org/10.3389/fams.2023.1128224.

[6] Carlo R. Laing. “Derivation of a neural field model from a network of theta neurons”. In: Physical Review E 90.1 (2014), p. 010901. doi: 10.1103/physreve.90.010901. URL: https://doi.org/10.1103/physreve.90.010901.

[7] Áine Byrne, Daniele Avitabile, and Stephen Coombes. “Next-generation neural field model: The evolution of synchrony within patterns and waves”. In: Physical Review E 99.1 (2019), p. 012313. doi: 10.1103/physreve.99.012313. URL: https://doi.org/10.1103/physreve.99.012313.

[8] Áine Byrne et al. “Next-generation neural mass and field modeling”. In: Journal of Neurophysiology 123.2 (2020), pp. 726–742. doi: 10.1152/jn.00406.2019. URL: https://doi.org/10.1152/jn.00406.2019.

[9] Edward Ott and Thomas M. Antonsen. “Long time evolution of phase oscillator systems”. In: Chaos: An Interdisciplinary Journal of Nonlinear Science 19.2 (2009), p. 023117. doi: 10.1063/1.3136851. URL: https://doi.org/10.1063/1.3136851.

[10] Bastian Pietras and Andreas Daffertshofer. “Ott–Antonsen attractiveness for parameter-dependent oscillatory systems”. In: Chaos 26 (2016), p. 103101. doi: 10.1063/1.4963371.

[11] Bastian Pietras, Rok Cestnik, and Arkady Pikovsky. “Exact finite-dimensional description for networks of globally coupled spiking neurons”. In: Physical Review E 107 (2023), p. 024315. doi: 10.1103/PhysRevE.107.024315.

[12] Ronaldo García Reyes and Eduardo Martinez Montes. “Modeling neural activity in neurodegenerative diseases through a neural field model with variable density of neurons”. In: bioRxiv (2022). Preprint; not peer reviewed. doi: 10.1101/2022.08.23.504980. URL: https://doi.org/10.1101/2022.08.23.504980.

[13] Agus Hartoyo et al. “Parameter estimation and identifiability in a neural population model for electro-cortical activity”. In: PLOS Computational Biology 15.5 (2019), e1006694. doi: 10.1371/journal.pcbi.1006694.

[14] A. Raue et al. “Structural and practical identifiability analysis of partially observed dynamical models by exploiting the profile likelihood”. In: Bioinformatics 25.15 (2009), pp. 1923–1929. doi: 10.1093/bioinformatics/btp358. URL: https://doi.org/10.1093/bioinformatics/btp358.

[15] Oana-Teodora Chis, Julio R. Banga, and Eva Balsa-Canto. “Structural Identifiability of Systems Biology Models: A Critical Comparison of Methods”. In: PLoS ONE 6.11 (2011), e27755. doi: 10.1371/journal.pone.0027755. URL: https://doi.org/10.1371/journal.pone.0027755.

[16] Willem de Haan et al. “Activity Dependent Degeneration Explains Hub Vulnerability in Alzheimer’s Disease”. In: PLoS Computational Biology 8.8 (2012), e1002582. doi: 10.1371/journal.pcbi.1002582. URL: https://doi.org/10.1371/journal.pcbi.1002582.

[17] Willem de Haan et al. “Altering neuronal excitability to preserve network connectivity in a computational model of Alzheimer’s disease”. In: PLOS Computational Biology 13.9 (2017), e1005707. doi: 10.1371/journal.pcbi.1005707. URL: https://doi.org/10.1371/journal.pcbi.1005707.

[18] Anisleidy González Mitjans et al. “Accurate and Efficient Simulation of Very High-Dimensional Neural Mass Models with Distributed-Delay Connectome Tensors”. In: NeuroImage 274 (2023), p. 120137. doi: 10.1016/j.neuroimage.2023.120137. URL: https://doi.org/10.1016/j.neuroimage.2023.120137.

[19] G. B. Ermentrout and N. Kopell. “Parabolic Bursting in an Excitable System Coupled with a Slow Oscillation”. In: SIAM Journal on Applied Mathematics 46.2 (1986), pp. 233–253. doi: 10.1137/0146017. URL: https://doi.org/10.1137/0146017.

[20] Paul So, Tanushree B. Luke, and Ernest Barreto. “Networks of theta neurons with time-varying excitability: Macroscopic chaos, multistability, and final-state uncertainty”. In: Physica D: Nonlinear Phenomena 267 (2014), pp. 16–26. doi: 10.1016/j.physd.2013.04.009. URL: https://doi.org/10.1016/j.physd.2013.04.009.

[21] Bastian Pietras and Ernest Montbrió. “Heterogeneous populations of quadratic integrate-and-fire neurons: on the generality of Lorentzian distributions”. In: arXiv (2024). Preprint; version 1, 26 September 2024. doi: 10.48550/arXiv.2409.18278. eprint: 2409.18278. URL: https://arxiv.org/abs/2409.18278.

[22] Rok Cestnik and Arkady Pikovsky. “Hierarchy of Exact Low-Dimensional Reductions for Populations of Coupled Oscillators”. In: Physical Review Letters 128.5 (2022), p. 054101. doi: 10.1103/physrevlett.128.054101. URL: https://doi.org/10.1103/physrevlett.128.054101.

[23] Irina V. Tyulkina et al. “Dynamics of Noisy Oscillator Populations beyond the Ott-Antonsen Ansatz”. In: Physical Review Letters 120.26 (2018), p. 264101. doi: 10.1103/physrevlett.120.264101. URL: https://doi.org/10.1103/physrevlett.120.264101.

[24] Carlo R Laing. “Numerical Bifurcation Theory for High-Dimensional Neural Models”. In: The Journal of Mathematical Neuroscience 4.1 (2014), p. 13. doi: 10.1186/2190-8567-4-13. URL: https://doi.org/10.1186/2190-8567-4-13.

[25] James Rankin et al. “Continuation of Localized Coherent Structures in Nonlocal Neural Field Equations”. In: SIAM Journal on Scientific Computing 36.1 (2014), B70–B93. doi: 10.1137/130918721. URL: https://doi.org/10.1137/130918721.

[26] W. Govaerts, Yu. A. Kuznetsov, and B. Sijnave. “Numerical Methods for the Generalized Hopf Bifurcation”. In: SIAM Journal on Numerical Analysis 38.1 (2000), pp. 329–346. doi: 10.1137/s0036142999352552. URL: https://doi.org/10.1137/s0036142999352552.

[27] Carlo R. Laing and Oleh E. Omel’chenko. “Periodic solutions in next generation neural field models”. In: Biological Cybernetics 117. 4-5 (2023), pp. 259–274. doi: 10.1007/s00422-023-00969-6. URL: https://doi.org/10.1007/s00422-023-00969-6.

[28] O E Omel’chenko. “Periodic orbits in the Ott–Antonsen manifold”. In: Nonlinearity 36.2 (2023), pp. 845–861. doi: 10.1088/1361-6544/aca94c. URL: https://doi.org/10.1088/1361-6544/aca94c.

[29] Yuri A. Kuznetsov. Elements of Applied Bifurcation Theory. 2nd ed. Vol. 112. Applied Mathematical Sciences. New York: Springer, 1998. URL: https://wwwf.imperial.ac.uk/~dturaev/kuznetsov.pdf.

[30] Yuri A. Kuznetsov. “Numerical Normalization Techniques for All Codim 2 Bifurcations of Equilibria in ODE’s”. In: SIAM Journal on Numerical Analysis 36.4 (1999), pp. 1104–1124. doi: 10.1137/S0036142998335005. URL: https://epubs.siam.org/doi/10.1137/S0036142998335005.

[31] Yuri A. Kuznetsov et al. “Switching to nonhyperbolic cycles from codim 2 bifurcations of equilibria in ODEs”. In: Physica D: Nonlinear Phenomena 237.23 (2008), pp. 3061–3068. doi: 10.1016/j.physd.2008.06.006. URL: https://doi.org/10.1016/j.physd.2008.06.006.

[32] Frank Schilder, Hinke M. Osinga, and Werner Vogt. “Continuation of Quasi-periodic Invariant Tori”. In: SIAM Journal on Applied Dynamical Systems 4.3 (2005), pp. 459–488. doi: 10.1137/040611240. URL: https://epubs.siam.org/doi/10.1137/040611240.

[33] Vladimir V. Klinshov and Sergey Yu. Kirillov. “Shot noise in next-generation neural mass models for finite-size networks”. In: Physical Review E 106.6 (2022), p. L062302. doi: 10.1103/physreve.106.l062302. URL: https://doi.org/10.1103/physreve.106.l062302.

[34] Si-Wei Qiu and Carson C. Chow. “Finite-size effects for spiking neural networks with spatially dependent coupling”. In: Physical Review E 98.6 (2018), p. 062414. doi: 10.1103/physreve.98.062414. URL: https://doi.org/10.1103/physreve.98.062414.

[35] Bastian Pietras, Valentin Schmutz, and Tilo Schwalger. “Mesoscopic description of hippocampal replay and metastability in spiking neural networks with short-term plasticity”. In: PLOS Computational Biology 18.12 (2022), e1010809. doi: 10.1371/journal.pcbi.1010809. URL: https://doi.org/10.1371/journal.pcbi.1010809.

[36] Paul C. Bressloff and Jay M. Newby. “Metastability in a Stochastic Neural Network Modeled as a Velocity Jump Markov Process”. In: SIAM Journal on Applied Dynamical Systems 12.3 (2013), pp. 1394–1435. doi: 10.1137/120898978.

[37] H.A. Kramers. “Brownian motion in a field of force and the diffusion model of chemical reactions”. In: Physica 7.4 (1940), pp. 284–304. doi: 10.1016/s0031-8914(40)90098-2. URL: https://doi.org/10.1016/s0031-8914(40)90098-2.

[38] Peter Hänggi, Peter Talkner, and Michal Borkovec. “Reaction-rate theory: fifty years after Kramers”. In: Reviews of Modern Physics 62.2 (1990), pp. 251–341. doi: 10.1103/revmodphys.62.251. URL: https://doi.org/10.1103/revmodphys.62.251.

[39] E. L. Kaplan and Paul Meier. “Nonparametric Estimation from Incomplete Observations”. In: Journal of the American Statistical Association 53.282 (1958), pp. 457–481. doi: 10.1080/01621459.1958.10501452. URL: https://doi.org/10.1080/01621459.1958.10501452.

[40] Irmantas Ratas and Kestutis Pyragas. “Macroscopic oscillations of a quadratic integrate-and-fire neuron network with global distributed-delay coupling”. In: Physical Review E 98.5 (2018), p. 052224. doi: 10.1103/physreve.98.052224. URL: https://doi.org/10.1103/physreve.98.052224.

[41] Federico Devalle, Ernest Montbrió, and Diego Pazó. “Dynamics of a large system of spiking neurons with synaptic delay”. In: Physical Review E 98.4 (2018), p. 042214. doi: 10.1103/physreve.98.042214. URL: https://doi.org/10.1103/physreve.98.042214.

[42] Jose M. Esnaola-Acebes et al. “Synchrony-induced modes of oscillation of a neural field model”. In: Physical Review E 96.5 (2017), p. 052407. doi: 10.1103/physreve.96.052407. URL: https://doi.org/10.1103/physreve.96.052407.

[43] Pau Clusella and Ernest Montbrió. “Exact low-dimensional description for fast neural oscillations with low firing rates”. In: Physical Review E 109 (2024), p. 014229. doi: 10.1103/PhysRevE.109.014229.

[44] Zachary P. Kilpatrick, Stefanos E. Folias, and Paul C. Bressloff. “Traveling Pulses and Wave Propagation Failure in Inhomogeneous Neural Media”. In: SIAM Journal on Applied Dynamical Systems 7.1 (2008), pp. 161–185. doi: 10.1137/070699214. URL: https://doi.org/10.1137/070699214.

[45] Paul C. Bressloff. “Traveling fronts and wave propagation failure in an inhomogeneous neural network”. In: Physica D: Nonlinear Phenomena 155.1–2 (2001), pp. 83–100. doi:10.1016/S0167-2789(01)00266-4. URL: https://doi.org/10.1016/S0167-2789(01)00266-4.

[46] Áine Byrne et al. “Mean-Field Models for EEG/MEG: From Oscillations to Waves”. In: Brain Topography 35.1 (2022), pp. 36–53. doi: 10.1007/s10548-021-00842-4. URL: https://doi.org/10.1007/s10548-021-00842-4.

[47] Ronald Garcia-Reyes, Julien Vezoli, and Pedro A. Valdes–Sosa. “Spatial transcriptomic programs relate to spectrolaminar rhythms across macaque cortex”. In: bioRxiv (2026). Preprint; not peer reviewed. doi: 10.64898/2026.07.08.737187. URL: https://doi.org/10.64898/2026.07.08.737187.

[48] Ronaldo Garcia Reyes et al. “Lifespan development of EEG alpha and aperiodic component sources is shaped by the connectome and axonal delays”. In: National Science Review 13.7 (2026), wag076. doi: 10.1093/nsr/nwag076. URL: https://doi.org/10.1093/nsr/nwag076.

[49] Ying Wang et al. “The influence of nonlinear resonance on human cortical oscillations”. In: Communications Biology 9.1 (2026), p. 605. doi: 10.1038/s42003-026-10164-5. URL: https://doi.org/10.1038/s42003-026-10164-5.

## Supplementary References

[1] W. Govaerts, Yu. A. Kuznetsov, and B. Sijnave. “Numerical Methods for the Generalized Hopf Bifurcation”. In: SIAM Journal on Numerical Analysis 38.1 (2000), pp. 329–346. doi: 10.1137/s0036142999352552. URL: https://doi.org/10.1137/s0036142999352552.

[2] Carlo R Laing. “Numerical Bifurcation Theory for High-Dimensional Neural Models”. In: The Journal of Mathematical Neuroscience 4.1 (2014), p. 13. doi: 10.1186/2190-8567-4-13. URL: https://doi.org/10.1186/2190-8567-4-13.

[3] A. Raue et al. “Structural and practical identifiability analysis of partially observed dynamical models by exploiting the profile likelihood”. In: Bioinformatics 25.15 (2009), pp. 1923–1929. doi: 10.1093/bioinformatics/btp358. URL: https://doi.org/10.1093/bioinformatics/btp358.

[4] Oana-Teodora Chis, Julio R. Banga, and Eva Balsa-Canto. “Structural Identifiability of Systems Biology Models: A Critical Comparison of Methods”. In: PLoS ONE 6.11 (2011), e27755. doi: 10.1371/journal.pone.0027755. URL: https://doi.org/10.1371/journal.pone.0027755.

[5] Ernest Montbrió, Diego Pazó, and Alex Roxin. “Macroscopic Description for Networks of Spiking Neurons”. In: Physical Review X 5.2 (2015), p. 021028. doi: 10.1103/physrevx.5.021028. URL: https://doi.org/10.1103/physrevx.5.021028.

[6] Vladimir V. Klinshov and Sergey Yu. Kirillov. “Shot noise in next-generation neural mass models for finite-size networks”. In: Physical Review E 106.6 (2022), p. L062302. doi: 10.1103/physreve.106.l062302. URL: https://doi.org/10.1103/physreve.106.l062302.

[7] Si-Wei Qiu and Carson C. Chow. “Finite-size effects for spiking neural networks with spatially dependent coupling”. In: Physical Review E 98.6 (2018), p. 062414. doi: 10.1103/physreve.98.062414. URL: https://doi.org/10.1103/physreve.98.062414.

[8] Denis Villemonais. “Interacting Particle Systems and Yaglom Limit Approximation of Diffusions with Unbounded Drift”. In: Electronic Journal of Probability 16.61 (2011), pp. 1663–1692. doi: 10.1214/EJP.v16-925. URL: https://www.maths.tcd.ie/EMIS/journals/EJP-ECP/article/view/925.html.

[9] E. L. Kaplan and Paul Meier. “Nonparametric Estimation from Incomplete Observations”. In: Journal of the American Statistical Association 53.282 (1958), pp. 457–481. doi: 10.1080/01621459.1958.10501452. URL: https://doi.org/10.1080/01621459.1958.10501452.

[10] H.A. Kramers. “Brownian motion in a field of force and the diffusion model of chemical reactions”. In: Physica 7.4 (1940), pp. 284–304. doi: 10.1016/s0031-8914(40)90098-2. URL: https://doi.org/10.1016/s0031-8914(40)90098-2.

[11] Peter Hänggi, Peter Talkner, and Michal Borkovec. “Reaction-rate theory: fifty years after Kramers”. In: Reviews of Modern Physics 62.2 (1990), pp. 251–341. doi: 10.1103/revmodphys.62.251. URL: https://doi.org/10.1103/revmodphys.62.251.

[12] Carlo R. Laing and Oleh E. Omel’chenko. “Periodic solutions in next generation neural field models”. In: Biological Cybernetics 117. 4-5 (2023), pp. 259–274. doi: 10.1007/s00422-023-00969-6. URL: https://doi.org/10.1007/s00422-023-00969-6.

[13] O E Omel’chenko. “Periodic orbits in the Ott–Antonsen manifold”. In: Nonlinearity 36.2 (2023), pp. 845–861. doi: 10.1088/1361-6544/aca94c. URL: https://doi.org/10.1088/1361-6544/aca94c.

[14] Áine Byrne, Daniele Avitabile, and Stephen Coombes. “Next-generation neural field model: The evolution of synchrony within patterns and waves”. In: Physical Review E 99.1 (2019), p. 012313. doi: 10.1103/physreve.99.012313. URL: https://doi.org/10.1103/physreve.99.012313.

[15] Yuri A. Kuznetsov. Elements of Applied Bifurcation Theory. 2nd ed. Vol. 112. Applied Mathematical Sciences. New York: Springer, 1998. URL: https://wwwf.imperial.ac.uk/~dturaev/kuznetsov.pdf.

[16] Frank Schilder, Hinke M. Osinga, and Werner Vogt. “Continuation of Quasi-periodic Invariant Tori”. In: SIAM Journal on Applied Dynamical Systems 4.3 (2005), pp. 459–488. doi: 10.1137/040611240. URL: https://epubs.siam.org/doi/10.1137/040611240.

[17] Áine Byrne et al. “Mean-Field Models for EEG/MEG: From Oscillations to Waves”. In: Brain Topography 35.1 (2022), pp. 36–53. doi: 10.1007/s10548-021-00842-4. URL: https://doi.org/10.1007/s10548-021-00842-4.

